# A sibling study of variation in parental mutation rates

**DOI:** 10.64898/2026.05.10.724105

**Authors:** Vanesa Getseva, Pelin Poyraz, Anastasia Stolyarova, Ipsita Agarwal, Molly Przeworski

## Abstract

The number of germline *de novo* mutations (DNMs) differs among people, depending on chance and parental ages. To explore additional causes, we developed an approach to call DNMs from differences between siblings in genomic regions inherited identical by descent from both parents. Applying it to whole genome sequences from 28,985 sibling pairs of diverse genetic ancestries in the UK Biobank and All of Us datasets, as well as 2,330 trios, we identified >850K autosomal DNMs in 27,645 parental sets. Across genetic ancestry groups, there are subtle differences in the spectrum but no difference in the total DNM rates, accounting for age. Testing for associations between parental mutation phenotypes and their burden of loss-of-function and deleterious missense variants in a set of 180 candidate genes, we found evidence for mutator alleles in *REV1* and *LIG1*. This study illustrates how relatedness in biobanks can be used to uncover determinants of germline mutation.

## Introduction

Germline mutations are found in every cell of an offspring’s body. Most are probably without fitness effects and a tiny fraction are advantageous, but a non-negligible proportion is deleterious. That subset underlies genetic diseases, notably developmental disorders, and contributes to risk for numerous cancers. In that regard, not everyone faces the same risk: while on average, a person is born with on the order of 100 *de novo* mutations (DNMs) [1, 2], the number varies among individuals. Identifying sources of this variation, and the extent to which they may be modifiable, is important for human and cancer genetics; it is also central to our understanding of how mutation rates evolve [3, 4].

Beyond chance fluctuations, the main reason for differences in the number of germline DNMs is parental age at reproduction: the number of mutations that a person inherits increases by (on average) ∼1.5 point mutations per year of their father’s age and ∼0.4 of their mother’s [1, 5], and (after accounting for stochastic variation) these age effects together account for ∼70% of inter-individual variance [4]. Given that mutation rates are a property of parents and very hard to estimate precisely, identifying additional sources of variance has proven elusive.

The largest study to date examined whole genome sequences for approximately 14K trios ascertained in the United Kingdom (UK) for a child with a rare disease, as well as exome sequences in an additional 8K trios [4]. In a dozen cases, the child inherited an unusually high number of mutations given the parental ages. In half, the cause was likely a course of chemotherapy in the father, indicating that, although the germline is well protected from exogenous damage [6, 7], mutagens can have an effect on germline mutations [4]. The same study also found evidence for likely genetic effects: in two cases in which the number of DNMs in the genome of the child was several-fold higher than expected, the fathers were homozygous for a LoF or deleterious missense mutation in a DNA repair gene (*XPC* and *MPG*). Elevated mutation rates have also been reported for offspring of fathers with a single missense mutation in *POLE*, *POLD1* and of mothers with compound heterozygous missense mutations in *MUTYH* [8–10]. Additional examples have been identified in other species: in particular, modifier alleles have been mapped to *OGG1* in yeast [11] and to *MUTYH* and *OGG1* in mice [12, 13], where they influence the rate of C*>*A mutations. An elevated mutation rate of C*>*T mutations has also been associated with disruptions of *MBD4* in rhesus macaques [14].

In humans, after controlling for parental ages, the number of mutations inherited by a child is estimated to have a heritability of 0.53 for fathers (for mothers, the point estimate is lower and not significantly different from 0) [4]. While this result indicates that there are genetic effects to be found, no common variants were significantly associated with mutation rates in a genome-wide association study [15]. The lack of findings could reflect low power, especially given the noisiness of the phenotype. Alternatively, it might suggest strong selection against most mutator alleles, owing to the additional mutations that they cause [16, 17], which leads them to be maintained at low frequency in the population. Accordingly, an analysis of heritability stratified by allele frequency suggested that the paternal heritability resides exclusively among alleles at low minor allele frequency (*<* 1%; [4]) and mutator alleles identified to date segregate at frequency well below 1% [18]. Taken together, these findings suggest that the known mutator alleles represent only the large-effect tail of rare variants that influence germline mutation rates in humans. If so, burden tests may be more powerful than GWAS for identifying additional cases.

Further evidence that mutator alleles or environmental effects play a role in shaping human germline mutation rates comes from analyses of polymorphism data, which highlight differences across ancestry groups in the number and types of mutations [19–22], as well as from differences in the mutation spectrum among primate species [23, 24]. Notably, one mutation type (TCC*>*TTC) was discovered to be more common in people with recent European genetic ancestry relative to those with East Asian or African ancestry, due to a mutational pulse in their ancestors tens of thousands of years ago [24, 25]. Differences between genetic ancestry were also found in a reanalysis of UK trios: accounting for parental ages, the mean DNM rate was elevated in 198 parental sets of recent African ancestry compared to those of European (8,102 parental sets), South Asian (1,249) or “Admixed American” (215) ancestry, and significant differences in mutation spectra were also detected between the European and South Asian ancestry groups [15]. As the authors note, these differences may reflect environmental exposures rather than genetic effects.

The only environmental data available for these samples, smoking history, had a statistically significant effect on the number of DNMs, but the mutational signature known to be associated with tobacco damage was not detected among the DNMs, so the finding remains to be confirmed [15]. Nonetheless, that germline mutation rates have been shown to be heritable in males and associated with chemotherapy and possibly smoking suggests that there are genetic and environmental influences yet to be identified.

## Results

### Our approach

To characterize variation in germline mutation rates among humans and test for such influences, we developed a method that leverages whole-genome sequencing data from siblings. Closely related individuals share large segments of their genomes that are identical by descent (IBD). Differences within these segments can be used to identify mutations that have arisen since their most recent common ancestor [21, 26]. Here, we relied on the fact that siblings share both of their chromosomes (i.e., are IBD2) in a quarter of their genome on average. In these regions, any genetic difference between siblings must result either from a germline DNM that occurred in the transmission from a parent to one child but not the other (Figure 1a), or, more rarely, from a gene conversion event in one of the parents (Figure 1b).

**Fig. 1.**
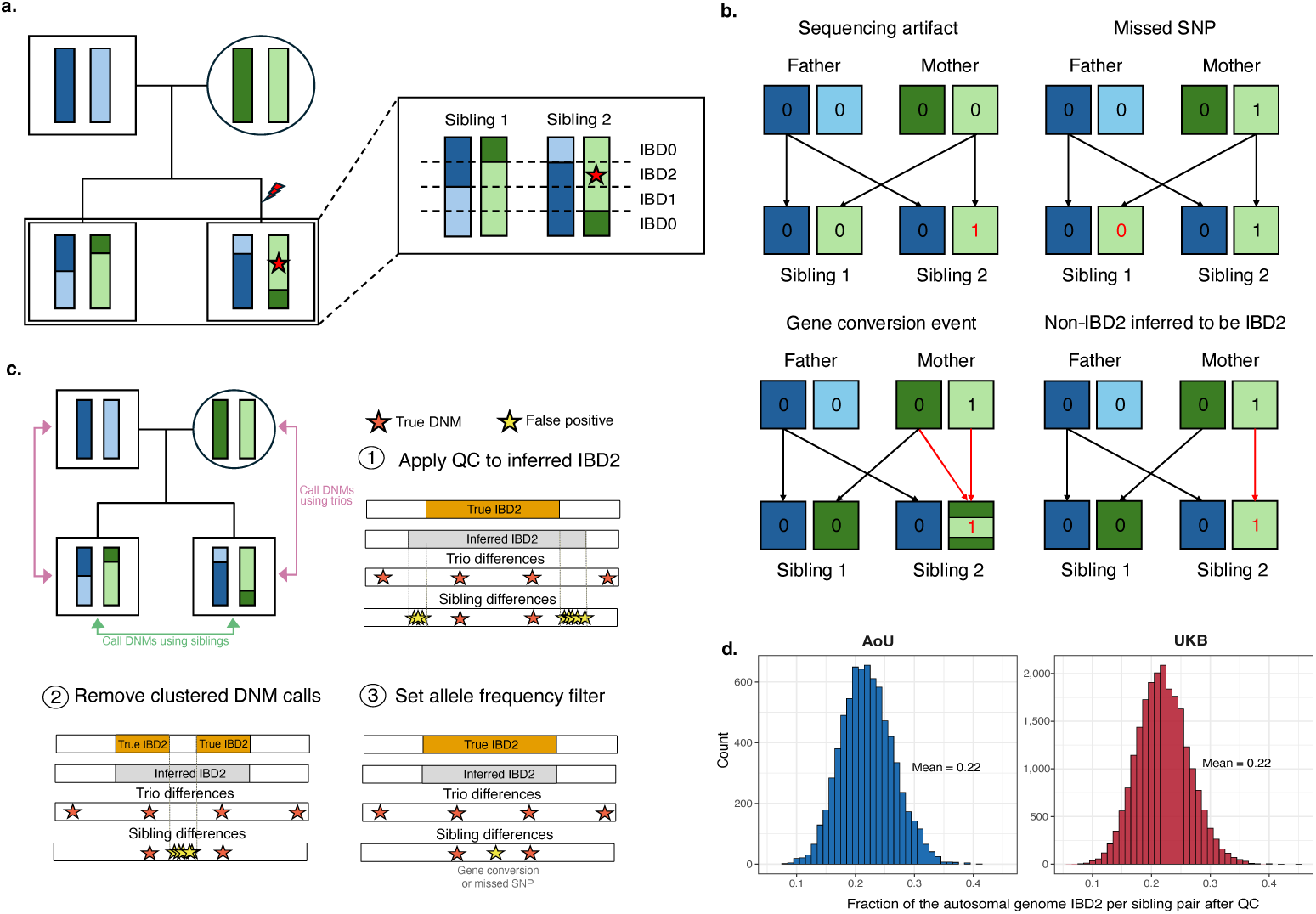
Overview of the sibling-based DNM calling approach. (a) DNMs appear as differences in genotype calls between siblings in IBD2 regions (with paternal haplotypes in blue and maternal in green). A red star represents a DNM in Sibling 2. (b) Alternative sources of genotype differences between siblings in IBD2 segments. Boxes indicate parental haplotypes at a given locus, with paternal haplotypes in blue and maternal in green. The numbers inside represent genotypes: 0 is the reference allele and 1 the alternative. Arrows indicate the transmission of haplotypes to offspring. (c) Data from quads (parents and two offspring) inform QC steps. False positive DNM calls (yellow stars) resulting from errors in IBD2 inference, missed SNPs and gene conversion events appear in the sibling approach but not the trio approach. (d) Distribution of the fraction of the autosomal genome retained as IBD2 per sibling pair after QC in UKB and AoU.

Sibling genomes also allow one to impute parental genotypes [27] and test for an effect of modifiers on mutation rates, as discussed below. In this regard, we note that when the goal is to estimate the mutation rate of a set of parents, *µ*, a sibling pair is roughly half as informative as a trio. Ignoring the separate contribution of the two sexes, the offspring of a trio provides one estimate of *µ* based on the whole genome via comparison of the offspring’s entire genome with the genomes of the parents. In turn, a sibling pair provides two (quasi-)independent estimates of *µ* via comparison of their IBD2 segments, which cover 1*/*4 of the genome in expectation. There are two estimates available for this quarter of the genome because each sibling can serve as a comparison for the other. Thus, a given sibling pair provides approximately half the information of a trio study, without the need for data from the parents.

In practice, apparent differences between siblings can also arise from sequencing artifacts, missed heterozygous calls in one sibling, or IBD inference errors (Figure 1b). We therefore identified DNMs by inferring which regions of the genome are IBD2 for every sibling pair and applied filters to minimize the contributions of errors and gene conversion events before identifying differences between them.

We applied this approach to two large population cohort studies: the UK Biobank (henceforth UKB), which enrolled ∼500K individuals between the ages of 39 and 69 living in the United Kingdom, and the All of Us study (AoU), which in 2026 included genomes from ∼415K adults recruited across the United States. Both studies collected genotype array data for these samples as well as Illumina whole genome sequencing (WGS) data from whole blood or saliva at a mean fold coverage of ∼30X [28, 29] (Methods 1). Using the genetic data, we identified identical genomes, which could be accidental duplicates or, in a subset of cases, monozygotic twins; trios consisting of mother, father and offspring; quads consisting of both parents and two offspring; and sibling pairs (with or without parental data available) (see Methods 2; Table 1).

**Table 1.** Sample size of sets of a given kind.

|  | UKB | AoU |
| --- | --- | --- |
| Monozygotic twin and<br>genomic duplicate pairs | 178 | 2,242 |
| Parent-offspring trios | 1,038 | 1,292 |
| Quads | 35 | 192 |
| Sibling pairs | 21,660 | 7,325 |
| Parental sets | 20,262 | 7,383 |

### A new approach to call DNMs

Before calling DNMs from sibling differences, we characterized the error profile of the WGS data in the two datasets and tuned our filters accordingly. To that end, we applied standard filters, then identified nucleotide differences between the 2,420 monozygotic twin and genomic duplicate pairs (henceforth referred to as “duplicates”). These differences should overwhelmingly reflect errors, given that no such differences are expected for true duplicates and approximately one constitutive mutation is expected to differ between monozygotic twins (see Methods 4). In the process, we discovered a much higher false positive rate for indels than point mutations, especially in AoU (Methods 4). In what follows, we therefore focused only on point mutations. To remove additional artifacts, we compiled a set of error-prone regions for the two biobanks and excluded differences within these regions, which substantially reduced the error rate (Methods 4).

Next, we called DNMs from the WGS data for parent-offspring trios, using a standard approach ([1]; see Methods 5). Applying the same quality filters and error-prone region exclusions as for duplicates, we identified a mean of 43.5 DNMs in the UKB (corresponding to a mean mutation rate of 9.48 × 10*^−^*^9^ per bp) and a mean of 56.7 DNMs in AoU (mean mutation rate of 1.24 × 10*^−^*^8^ per bp) (see Methods 5), for a total of 118,321 DNMs across 2,330 trios. The smaller number of DNMs in the UKB is expected from the younger ages of the parents, given the age enrollment criteria (see Methods 5).

Some of these trios are part of nuclear families in which there are also siblings. We used these 227 quads to decide on additional filters for the sibling-based approach (Figure 1c, Methods 7). Our goal was to exclude gene conversion events and minimize missed heterozygous calls and IBD inference errors (Figure 1b), all of which can produce differences between siblings due to inherited mutations rather than DNMs. To do so, we masked the parental data in the quads and treated them as we would siblings, inferring IBD2 segments and imposing filters on these segments and the putative differences within them. We then asked what fraction of DNM calls made solely based on siblings were also identified using the parental information. This comparison indicated that 6.7% in AoU and 10.3% in the UKB are found only in siblings, reflecting either false positive calls in the siblings or missed DNMs in the trio (see Methods 7).

As a complementary approach to estimating error rates of DNM calls based on siblings, we identified a subset of siblings with a parent or offspring in the sample and applied the same IBD2-inference approach and set of filters established in the quad analyses (see Methods 9). Data from parents and offspring for these siblings allowed us to quantify two sources of error in DNM calls: the rate at which a variant is missed in a sibling, leading an inherited variant to appear as a DNM, and the rate at which a sequencing artifact leads to a spurious DNM call. Summing up these two sources of error yields an estimated false discovery rate of 9.7% (95% CI: 6.9%-12.5%) in the UKB and 7.5% (95% CI: 6.5%-8.5%) in AoU, compared to an analogous rate of 4.5% for trio-based DNM calls [30]. Together, these analyses indicate that the vast majority of DNMs identified based on sibling differences are genuine.

We therefore proceeded to identify DNMs across all sibling pairs. As a first step, we identified an average of 21.9% of the genome shared IBD2 per pair in the UKB and 22.2% in AoU after quality control (when 25% is expected without filtering), with substantial variance among sibling pairs, as expected (Figure 1d). Since mutation rates are properties of parents, we do not expect large differences in mutation rates between siblings after accounting for their age difference; such cases are more likely to reflect elevated sequencing errors or clonal hematopoiesis than a true germline signal. We used this criterion to flag and exclude 3.1% of pairs in the UKB and 4.8% in AoU (see Methods 8). In the remaining 28,985 pairs of siblings, we identified a mean of 25.9 DNMs in the UKB (corresponding to a mean mutation rate of 1.28 × 10*^−^*^8^ per bp) and a mean of 27.0 in AoU (mean mutation rate of 1.34 × 10*^−^*^8^ per bp) (Figure 2a-b). As expected, the mean difference in the number of DNMs between siblings increases linearly with their difference in age, reflective of parental age effects (*p <* 10*^−^*^16^; Supplementary Figure 1). Furthermore, the fraction of DNMs within meiotic double-strand break hotspots [31] (1.26% in the UKB and 1.27% in AoU) closely matches the expectation from trio studies of DNMs [32] (1.26%), indicating that they are not enriched for gene conversion events (Methods 7). The number of DNMs varies substantially across sibling pairs, reflecting chance differences in the fraction of their genomes that is IBD2, stochasticity in the number of mutations in those regions, and potentially differences in parental mutation rates (Figure 2a-b).

**Fig. 2.**
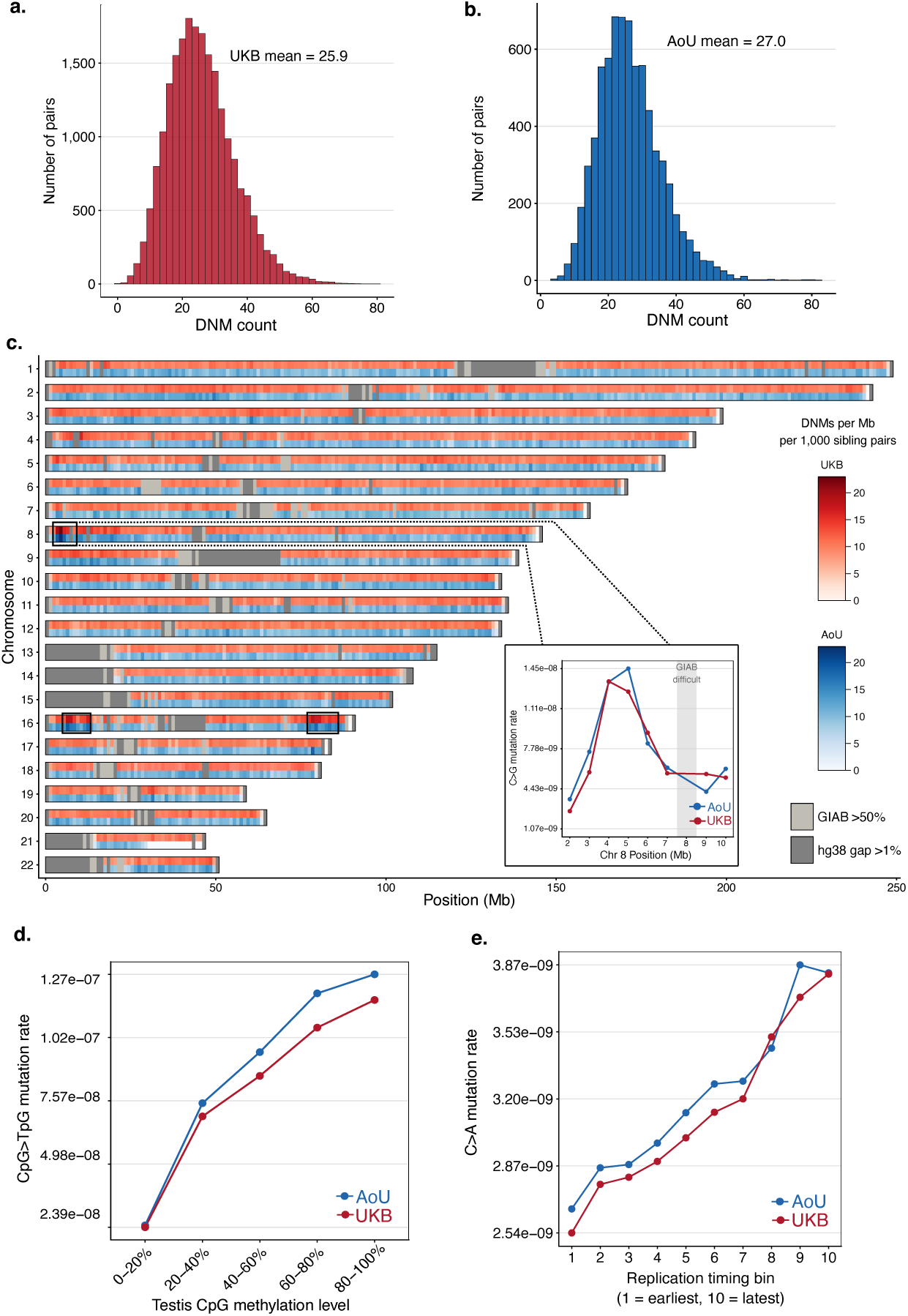
Features of germline mutagenesis are captured by sibling-based DNM calls. (a-b) DNM counts per sibling pair in the UKB and AoU. (c) DNM density in 1 Mb windows across the genome in AoU (blue) and UKB (red). Gray boxes are 1 Mb windows in which there is a *>*1% gap in hg38 or which overlap by *>* 50% with GIAB error-prone regions (Methods 4). Rectangles indicate regions previously reported to harbor high densities of C*>*G on chromosomes 8 and 16, especially in older mothers [1, 33–35]. Inset: C*>*G rates across the chromosome 8 enriched region. (d) CpG*>*TpG mutation rate across bins of CpG methylation level in testis. CpG sites were grouped into bins by methylation level and the average CpG*>*TpG mutation rate was calculated for each bin (Methods 10). (e) C*>*A mutation rates across bins of replication timing in lymphoblastoid cell lines. Bins were generated by calculating mean replication timing in 1 Mb windows and assigning a replication timing decile to each window (Methods 10). Points represent the average C*>*A mutation rate in each bin.

In total, we identified 734,715 DNMs from sibling pairs, distributed across all autosomes (with the exception of one segment on chromosome 21, in which no IBD2 sharing is identified in AoU; Figure 2c). The C*>*G mutation rate is elevated in specific regions of chromosomes 8 and 16 previously reported to show an enrichment of maternal C*>*G mutations [1, 33–35] (Figure 2c). In turn, the CpG*>*TpG mutation rate increases with local methylation level in testis (Figure 2d) and the C*>*A mutation rate increases with replication timing in lymphoblastoid cell lines (Figure 2e, Methods 10), as expected from previous analyses of germline mutations [5, 36]. More-over, the mutational spectrum of the DNM calls from siblings is highly consistent with DNMs previously identified in complete families [32], with a cosine similarity exceeding 0.99 (Supplementary Figure 2). In accordance with prior work indicating that germline point mutations are predominantly assigned to COSMIC SBS signatures SBS5 and SBS1 [37–39], most DNMs identified in siblings are attributed to these two signatures (Supplementary Figure 2). Thus, DNMs called from IBD2 segments in siblings recapitulate known features of germline mutations.

### Differences in germline mutation rates among genetic ancestry groups

The DNMs were identified in families with a diverse set of recent genetic ancestries. To stratify them accordingly, we assigned individuals in the UKB and AoU to ancestry groups based on genetic similarity to HGDP-1000 genomes individuals [40]. Specifically, we ran a principal component analysis (PCA) on the 3400 HGDP-1000 genome samples, then used k-means clustering to generate 10 genetic ancestry clusters (Figure 3a; Methods 11). Next, we assigned each offspring (from trios or sibling pairs) in UKB and AoU to the closest cluster. For the purposes of comparing mutation profiles across ancestry groups, we removed 352 genomes from the UKB and 1,894 from AoU with distances to the nearest cluster greater than the 95th percentile of HGDP-1KG distances, i.e., people whose genetic ancestry was likely a recent mixture of clusters. By this approach, 40,645 individuals in the UKB and 14,190 in AoU were assigned to a single genetic ancestry group.

**Fig. 3.**
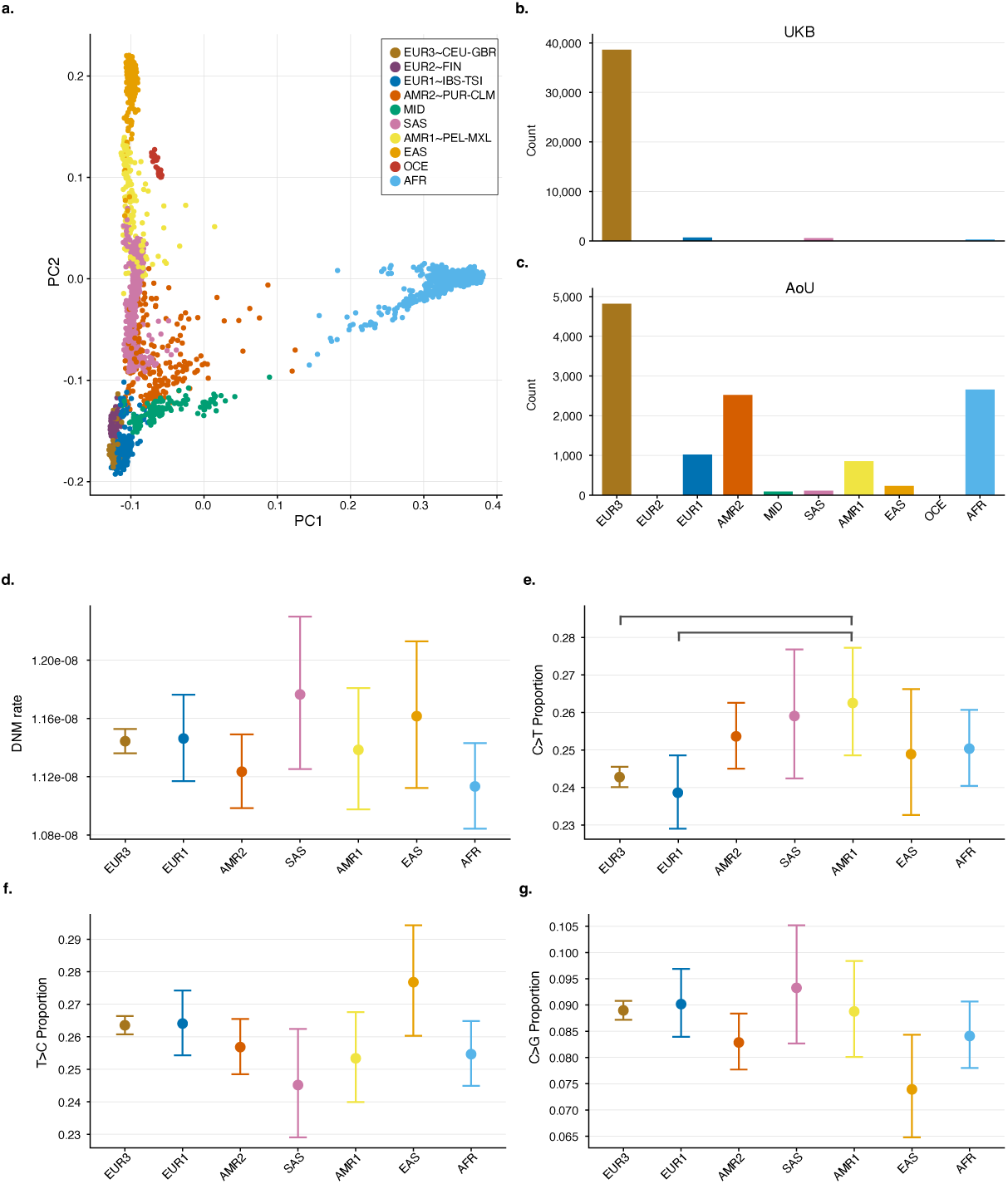
DNM rates and spectra across ancestry groups in UKB and AoU. (a) Principal component analysis of HGDP-1KG reference populations used to generate genetic ancestry groups. (b-c) Number of siblings and trio offspring assigned to each ancestry group in UKB and AoU (see Methods 11). Ancestry groups are ordered by the distance of their centroid to the centroid of the largest group (EUR3). (d) DNM rate per ancestry group, controlling for paternal ages and batch effects. Rates and 95% CIs shown are model predictions for UKB siblings at a paternal age of 29. Note that the y-axis is truncated (see Methods 11). (e-g) Proportion of non-CpG C*>*T, T*>*C, and C*>*G mutations in each ancestry group, controlling for paternal ages and batch effects (see Methods 11). Proportions and 95% CIs shown are model predictions for UKB siblings at paternal age 29. Brackets indicate significant pairwise differences (at an FDR *<*0.05).

In contrast to previous reports [15, 22, 33], we found total mutation rates to be very similar across ancestry groups. Using the seven largest groups (each of which includes more than 60 individuals with paternal age information), there is no discernible effect of ancestry on mean DNM rates, controlling for paternal ages and batch effects (p = 0.15; Methods 11). A previous study of DNMs reported a ∼4% higher mean number of DNMs in Africans than non-Africans, accounting for parental ages [15]. With 90% power to detect such a difference or greater (Figure 3d, Methods 11), we instead found a 2.4% *lower* mutation rate in the group labeled AFR compared to all others (at the same mean paternal age, *p* = 0.06; Supplementary Figure 3). Our finding is in the same direction (albeit of smaller magnitude; see Methods 11) as an inference based on substitution data, which suggested a ∼5% lower mutation rate per year in Africans compared to non-Africans [22].

We also examined differences in the mutation spectrum among genetic ancestry groups, by considering the proportion of seven mutation types (i.e., the six strand-collapsed, single-base substitutions, with C*>*T mutations split into two categories depending on CpG context.) Genetic ancestry is a significant predictor for C*>*T mutations outside of a CpG context (p = 0.04, after correction for seven tests) and marginally significant for C*>*G (p = 0.07) and T*>*C (p = 0.08) mutations, accounting for paternal ages and batch effects (Figure 3e-g, Methods 11). Comparing pairs of ancestry groups, the proportion of C*>*T mutations is significantly higher in AMR1 than in EUR3 (ratio = 1.09, p = 0.046, after FDR correction) and EUR1 (ratio = 1.07, p = 0.046) (Supplementary Table 1). In principle, these observations could arise from differences in maternal age effects among groups or from the allele frequency filter that we imposed on the DNMs called by the sibling approach (given that the proportion of true mutations excluded by this filter will depend on demographic history and the mutation rate of each specific type [41]). However, ancestry remains a significant predictor for the proportion of C*>*T mutations when using only DNMs identified in trios (where no allele frequency filter was applied), after accounting for batch effects and maternal as well as paternal ages (p = 0.007) (Methods 11).

The shifts in proportions of C*>*T among ancestry groups do not appear to be driven by specific trinucleotide contexts (Supplementary Table 2). In particular, we did not detect an elevated rate of TCC*>*TTC in EUR relative to other ancestries, as expected from the estimated time range for this mutation pulse, which is thought to have ended thousands of years ago [24, 25] (see Methods 11). We also did not find the enrichment of CCG*>*CTG in SAS relative to EUR reported by [21] for DNMs: in UKB siblings, for example, the proportion of this mutation type was 4.6% (95% CI: 4.4-4.9%) in SAS and 4.8% (95% CI: 4.7-4.9%) in EUR3 (Methods 11). In this regard, we note that despite similar labeling of continental-level groups, there may be salient differences between studies in the precise ancestry composition or set of environmental exposures within a group, e.g., within “AMR” or “SAS”.

### No effect of smoking on germline mutation rates

Given previous reports of a small (2.4%) but statistically significant increase in germline mutation rates when both parents were current or past smokers versus non-smokers [15], we tested whether we could detect a similar difference in our data. To that end, we used only the DNMs in 2,042 trios, as we do not know the smoking status of parents of siblings. By our approach, we did not find an effect of smoking, with 77% power to detect a 2.4% increase or greater. We also did not detect mutational signatures associated with tobacco damage among offspring of smokers (see Methods 12).

### Genetic effects on germline mutation phenotypes

To test for genetic variants that influence mutation phenotypes, we identified loss-of-function (LoF) variants (due to point mutations or indels) and putatively deleterious missense variants within a curated set of 210 genes involved in DNA repair and maintenance [42] (see Methods 13). To minimize annotation errors, we focused on LoF variants with an allele frequency ≤ 1% [43, 44]; to home in on deleterious missense variants, we considered the subset with REVEL score *>* 0.75, a computational predictor of pathogenicity [45].

When examining genetic effects on germline mutations, the relevant genotypes are those of the parents. In complete nuclear families, those data are available; when we based ourselves only on siblings, they are not, so we instead imputed them. Specifically, for all potential LoF or missense mutations in a gene, we used all the siblings with the same parents to infer the sum of parental genotypes on the basis of the sibling genotypes, whether the siblings are inferred to be IBD0, IBD1 or IBD2 at the site, and, where needed, population allele frequencies (following the approach of [27]; see Methods 13). For consistency, we used the same approach in complete nuclear families, encoding alleles as 0 (reference) and 1 (alternative), such that the sum represents the total number of alternative alleles carried by both parents. We then aggregated these sums across all LoF and deleterious missense sites within a gene to obtain an estimate of the number of disrupting variants per gene for 20,262 parental sets in the UKB and 7,383 in AoU.

The set of 210 genes includes a number of cases in which a single copy of a disrupting mutation is known to produce characteristic mutational signatures in one or more somatic tissues, including *BRCA1*, *BRCA2*, *EXO1*, *PALB2*, *PAXIP1*, *RIF1*, and *MLH1* [46]. Testing for an influence of disrupting variants (LoF or deleterious missense mutations) in these genes on the germline, we found no significant increase in the associated signatures (with the possible exception of *PAXIP1* ; see Methods 13 and Supplementary Table 3) and no elevation in the mutation rates of carriers (see Methods 13; Supplementary Table 4). These results lend support to to notion that the germline may be better protected against compromised repair activity than are specific somatic tissues [4]. We also tested two genes (*POLE* and *POLD1*) in which missense mutations were previously reported to increase germline mutations [8, 9]; neither gene showed elevated mutation rates in carriers relative to non-carriers (see Methods 13; Supplementary Table 4), with the important caveat that the exact variants studied previously were not present in the parents we analyzed.

Next, we tested systematically for an association between parental mutation phenotypes and the burden of disrupting mutations in the set of 210 DNA repair and maintenance genes (see Methods 13). To minimize sequencing artifacts, we excluded 30 genes that substantially overlap with error-prone GIAB regions (see Methods 13). To validate the imputation procedure for the remaining 180 genes, we compared imputed parental genotypes with the genotypes of parents in families with two or more offspring (see Methods 13). Defining parents as carriers if the sum of parental genotypes at a gene was ≥ 1 and non-carriers otherwise, parents are inferred to be carriers when they are not in 0.01% of cases (95% CI: 0.00%–0.02%) and true carriers are missed in 20.6% of cases (95% CI: 16.9%–24.3%) on average, across cohorts and annotations (Supplementary Table 13). In other words, by this procedure, we were very rarely wrong when imputing that a parent is a carrier, but misclassified a fifth of carriers as non-carriers, somewhat decreasing our power to detect associations. Across the 180 genes, we estimated that each pair of parents carries 1.14 LoF and 0.66 deleterious missense variants, corresponding to an average of roughly one disruptive variant per individual.

Prior work decomposing germline mutations into COSMIC single base substitution (SBS) signatures indicated that the majority of DNMs are assigned to SBS5, a signature with a relatively flat trinucleotide spectrum, and about one-sixth are assigned to SBS1, which primarily consists of transitions at CpG dinucleotides (i.e., CpG*>*TpG) [37, 39]. Smaller proportions of germline mutations are attributed to SBS39, which is characterized by C*>*G transversions, a substitution class known to increase steeply with maternal age [32] and a handful of signatures enriched for C*>*T and T*>*C transitions (e.g., SBS12, SBS16, and SBS30) [37, 39]. In addition, known mutator alleles in yeast and mice influence C*>*A germline mutation rates [11–13]; this type is also enriched in early embryonic mosaic variants relative to constitutive mutations in humans [30, 47]. On the basis of these findings, we conducted burden tests for six mutation rate phenotypes: the total mutation rate, as well as the rates of CpG*>*TpG, C*>*G, C*>*T not in a CpG context, T*>*C, and C*>*A (see Supplementary Figure 5 for the raw phenotype distributions).

Specifically, for each phenotype and gene, we tested whether families with a higher burden of LoF and deleterious missense variants in that gene have elevated mutation rates. To this end, we considered 27,645 parental sets, of which 2,121 include data from both parents and offspring, and 25,524 are based solely on siblings and thus provide a noisier estimate of parental mutation rates and genotypes. We combined LoF and deleterious missense mutations to maximize power [48] and required a minimum of 20 carriers, thus reducing the set of genes tested to 138. Parental ages at conception (reported or imputed), batch effects for the biobank and source, and six PCs were added as covariates (see Methods 13). We considered an additive model and, given the differences in power across phenotypes, we adjusted p-values for each phenotype separately using a Benjamini-Hochberg false discovery rate (FDR) correction.

Three of our mutation phenotypes showed signals of associations. Considering genes for which the rate of false discovery, q, is estimated to be less than one in five, *REV1* is associated with total mutation rates (q= 0.0806); *LIG1* (q = 0.0045) and *RAD51C* (q = 0.1507) with CpG*>*TpG mutation rates; and SLX4 (q = 0.1536) for C*>*T mutation rates at non-CpG sites (Supplementary Table 5; see the results for all genes in Supplementary Table S1). For *RAD51C*, this effect seems to be driven primarily by missense rather than LoF mutations (p = 2.5 × 10*^−^*^5^; Methods 13). For the other three genes, the difference between annotations was not significant, but with very limited power, in particular for SLX4 (Supplementary Table 6).

Of the five previously reported germline modifiers in humans, one is recessive in its effects (*MUTYH* [8, 49]) and two others (*MPG* and *XPC*) may be, as associations were only observed in one homozygous parent [4]. Consistent with a recessive mode of action, we detected no additive effects of disruptions in these three genes (Supplementary Table 4; *MPG* was originally excluded because of overlap with GIAB error-prone regions, so we examined it separately). To test for recessive effects, we considered the parent to be a carrier only when the parental genotype sum was ≥ 2 and a non-carrier otherwise, with the same covariates as in the additive model (see Methods 13). This approach has low power, both because two heterozygous parents will be misclassified as a homozygous carrier and because in this population cohort, few parents are truly recessive for a mutator allele (see Methods 13). Accordingly, our only finding is a tentative signal at *DNA2* (q = 0.1828) for T*>*C mutation rates. Focusing on *MPG*, *XPC*, *MUTYH*, three genes in which recessive mutator alleles have been reported [4, 8, 49], we identified no carriers (sum ≥ 2) in *MPG* or *XPC*, and fewer than 20 in *MUTYH*, for which there is no evidence of increased mutation rates (*p* = 0.29; Supplementary Table 4).

Overall, the application of additive and recessive models led to five candidate genes. Of these, three (*SLX4*, *RAD51C*, and *DNA2*) are involved in homologous recombination and the processing of complex DNA intermediates [50–53]; *DNA2* also processes DNA intermediates of Okazaki fragment maturation and BER [54, 55]. To our knowledge, there is no independent evidence linking them to the mutation types to which they are associated. By contrast, the genes with the strongest statistical evidence, *REV1* and *LIG1*, are plausible candidates for the specific phenotypes to which they are associated.

Specifically, our top hit for total mutation rates, *REV1*, is a Y-family DNA polymerase [56, 57] that is known to play two roles during translesion synthesis: its primary function is to recruit other Y-family polymerases and Pol *ζ* to bypass replication-blocking DNA lesions [58]. The REV1-Pol *ζ* complex has further been linked homologous recombination repair in human cells [59]. Less frequently, *REV1* has a catalytic role, incorporating deoxycytidines across damaged template G or abasic sites [57]. Under an additive model, we estimated that the total autosomal mutation rate in five carriers of LoF mutations and 21 carriers of deleterious missense variants in *REV1* is 16% higher on average than in non-carriers. This increase is seen at six of the seven mutation types (all but T*>*C; Figure 4a and Supplementary Figure 6a), seemingly independent of replication timing (Supplementary Figure 8, Methods 10). Based on a relatively small number of DNMs in carriers, the proportion assigned to SBS3, a signature associated with defective homologous recombination-based repair, is increased relative to non-carriers (from an estimated 0 to 9.2%; two-tailed *p* = 0.034; Supplementary Table 7; Methods 13). There is also weaker evidence for a small increase in SBS15 (0 to 1.8%, *p* = 0.061), a putative signature of defective DNA mismatch repair [38], as well as for a decrease in the contribution of SBS5 in carriers relative to non-carriers (58% instead of 83%, *p* = 0.067), consistent with findings linking SBS5 to translesion synthesis [6, 60, 61].

**Fig. 4.**
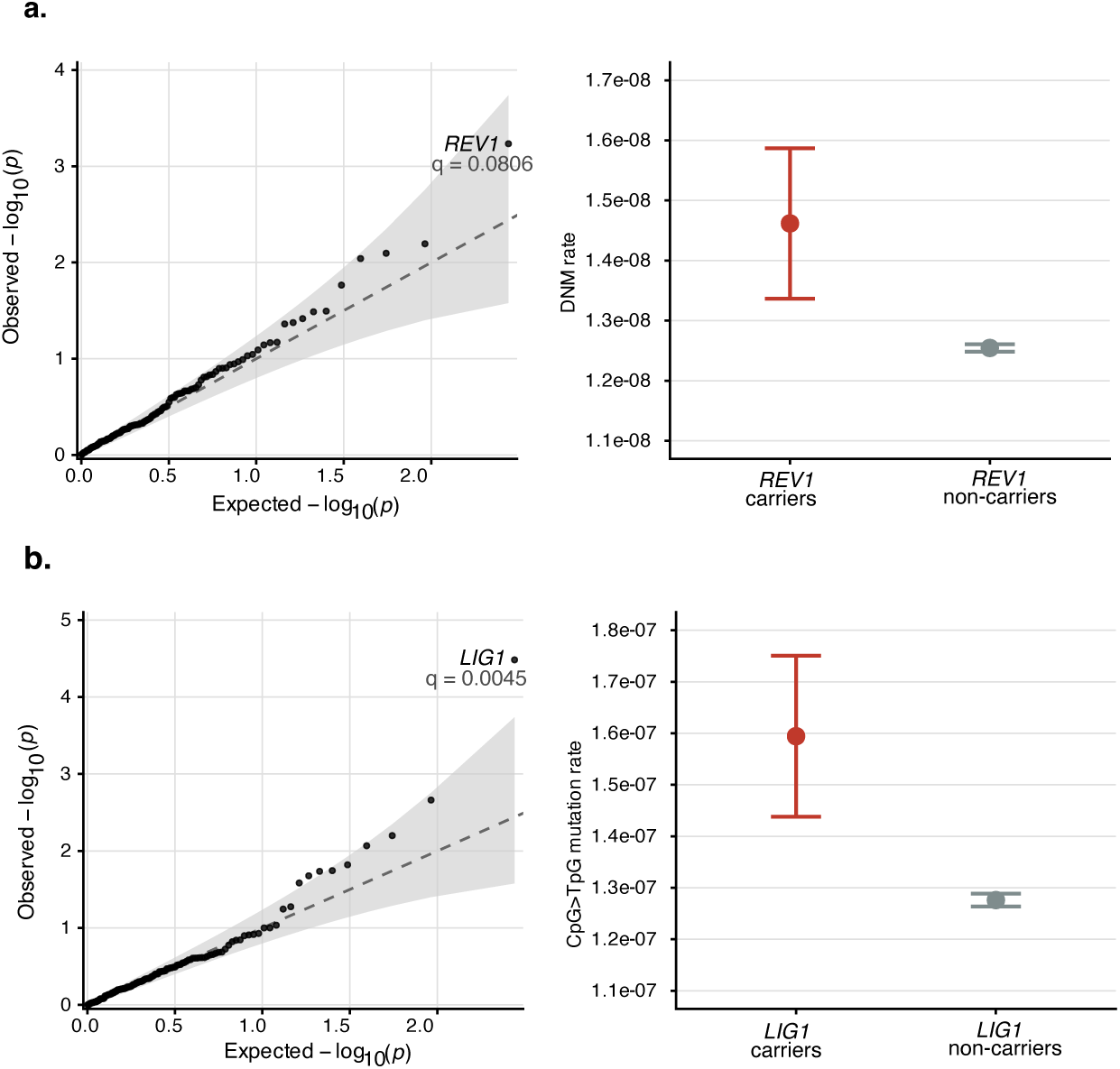
Burden test results for loss-of-function and deleterious missense variants in DNA repair and maintenance genes, for (a) the total autosomal DNM rate and (b) the autosomal rate of CpG*>*TpG mutations, under an additive model (see Methods 13). In the first column are Q-Q plots for 138 DNA repair genes with at least 20 carrier families in the UKB and AoU combined. The gray band represents the 95% CI under the null. The name of the gene with the strongest signal is indicated. In the second column are estimates of (a) the total autosomal DNM rate per bp and (b) the autosomal rate of CpG*>*TpG per bp in carriers versus non-carriers of disruptive variants. Shown are mean rates and 95% CIs for carriers (set to have a parental genotype sum = 1) and non-carriers (set to a sum = 0) based on predictions of an additive model for UKB siblings with genetic ancestry PCs from an EUR3 individual, at paternal age 29.0 and maternal age 27.1. Sample sizes for *REV1* : 26 carriers, 27,619 non-carriers; for *LIG1* model: 85 carriers, 27,560 non-carriers. Note that the y-axis is truncated.

In turn, the top association for the CpG*>*TpG mutation rate is *LIG1*, a DNA ligase involved in the last steps of both DNA replication and repair [62]. During DNA replication, its role is to ligate Okazaki fragments on the lagging strand. In DNA repair, LIG1 seals nicks generated after base excision repair (BER), nucleotide excision repair and mismatch repair (MMR) [62, 63]. LIG1 contributes to the fidelities of both DNA replication and repair by validating correct DNA base pairing and discriminating against damaged DNA [64]. CpG*>*TpG mutations arise at least in part from the deamination of 5-methylcytosine and are repaired through BER or, when due to replication errors, by MMR [65, 66]. Carriers of LoF and deleterious missense variants in *LIG1* exhibit a 24% increase in their CpG*>*TpG mutation rate on average (and an 8% increase in total mutation rate) (Figure 4b; Supplementary Figure 6b). The increase in carriers is not specific to methylated CpG sites (Supplementary Figure 8, Methods 10) and is also seen, to a lesser degree, at non-CpG sites (Supplementary Figure 6b). When DNMs in carriers are assigned to SBS signatures, there is a small increase in the proportion of SBS21 (from 0 to 1.1%, two-tailed *p* = 0.042), a DNA mismatch repair deficiency signature, as well as in SBS16 (0 to 2.9%, *p* = 0.050), the etiology of which is unknown; there is also tentative evidence for a decrease in SBS5 (83.1 to 71.8%, *p* = 0.067) (Supplementary Table 8; Methods 13) [38].

These gene associations are unlikely to result from population structure confounding, both because burden tests, which combine effects over many independent variants, are expected to be relatively robust to such confounding [67], and, in this instance, because no differences in total or CpG*>*TpG mutation rates are detected across ancestry groups (Figure 3). Nonetheless, as an additional check, we applied genomic control to adjust the false positive rate [68]: the non-centrality parameter was close to 1 in both cases (*λ* = 1.04 for total mutation rate and *λ* = 1.01 for CpG*>*TpG mutation rate) and accordingly, the evidence of association remained strong (*REV1*, q = 0.1045; *LIG1*, q = 0.0048; Supplementary Table S1). Moreover, as expected from the inclusion of batch effect in the burden tests, the associations at *REV1* and *LIG1* are not significantly different between UKB and AoU (see Methods 13; Supplementary Table 9). A further concern might be that these associations do not reflect germline effects in the parents, but rather clonal hematopoiesis in an offspring, with carriers of the disrupting mutations having increased somatic mutations in blood. This seems not to be the case: considering 16 families for *REV1* and 57 for *LIG1* in which one or more offspring inherited a disruptive variant and one or more did not, there is no significant difference in the mutation rates between offspring carrying the variant and those not carrying it (Mann-Whitney U test, p ≥ 0.1346; see Methods 13). These additional checks indicate that the associations at *REV1* and *LIG1* reflect disruptive effects on germline mutagenesis.

## Discussion

We conducted the first systematic test of the effects of rare variants on human germline mutation rates, focusing on a set of 138 candidate genes. At *REV1*, a single copy of a disrupting allele is associated with an additional ∼11 mutations genome-wide, and in *LIG1* with a couple of transitions at CpG sites and ∼6 mutations genome-wide. These estimated effect sizes are much smaller than those of known modifiers, which were found by examining candidate genes or identifying children with several-fold more mutations than expected [4, 8]. A caveat is that burden test estimates are an average over mutations that may have variable effect sizes. Nonetheless, these results suggest that, while the germline is well protected from the effects of smoking and gene disruptions that increase mutation rates in specific somatic tissues (Supplementary Table 3 and 4), there are modifier alleles segregating in population cohorts that only mildly increase the number of DNMs.

The diverse genetic ancestries included in the two biobanks also provide the opportunity to examine whether there is a difference in the total number of DNMs among groups, as previously reported [15, 22, 33]. Despite substantial power to detect relatively small differences, we found none (Methods 11). These analyses therefore establish that, accounting for parental ages, germline mutation rates are similar across broad-scale ancestry designations.

There are significant shifts in the mutational spectrum, however, notably in the proportion of C*>*T (Figure 3). Since such shifts must reflect differences in rates of specific mutation subtypes relative to each other, small differences in the total mutation rate among ancestry groups should become discernible in sufficiently large samples, short of strong stabilizing selection on the number of DNMs at reproduction [69].

The evolutionary processes that underlie these shifts—and the divergence in mutation spectrum among primate species [23, 24]—remain unclear. Even modifiers that introduce only a handful of additional mutations in the genome are expected to evolve under relatively effective purifying selection in recent human evolution [18]. As a consequence, such mutators are expected to be rare in all ancestry groups [70, 71], as is the case for LoF mutations in *LIG1* or *REV1* for instance [72], and therefore should contribute little or not at all to differences in mean mutation rates.

One possibility is that many mutator alleles are strictly recessive in their deleterious fitness effects [18, 73]. While such alleles could reach higher frequencies in some populations and contribute to ancestry differences [70, 74], they are unlikely to fix and lead to differences among species. Similarly, the observed differences among ancestry groups could arise from varying environmental exposures, but multiple lines of evidence suggest that the germline is well protected from mutagens [7] (Supplementary Figure 17) (with the exception of chemotherapy [4]), and environmental exposures are unlikely to explain why the divergence among species tracks the phylogeny. Alternatively, there may exist a much larger number of mutators with tiny effects, which evolve under very weak selection and hence can reach appreciable frequencies. While existing studies of common variants in humans suggest otherwise [4, 15], with sample sizes of ∼14K, they may be underpowered, especially for an imprecisely measured phenotype. Yet another explanation is that turnover in the mutation spectrum is nearly neutral, so long as the total mutation rate at reproduction remains roughly constant [75, 76]. To our knowledge, these scenarios remain to be modeled explicitly.

Much more also remains to be learned about the genetic architecture of variation in germline mutation. In this regard, this study provides a template, illustrating how sources of variation can be investigated without additional sequencing, by taking advantage of IBD sharing among genomes in publicly available genomic databases. We focused on IBD2 segments, in which DNMs can be identified without haplotype phasing; where such information is available, IBD1 segments also can be used. While in humans, the low genetic diversity levels mean that only a small proportion of DNMs called from siblings or trios can be phased with short read sequencing data, in a number of other species, including other primates, this fraction should be much higher [77]. Moreover, as long reads become more readily available, it will be feasible to phase almost all DNMs [2], opening the door to analyses of more distant relatives. Therefore, approaches such as the one developed here should enable systematic analyses of the determinants of germline mutation rates in humans, or any species in which large numbers of closely related individuals have been sequenced.

## Methods

### 1 Datasets used

#### UK Biobank

We analyzed the genotype array data (Data-Field 22418), the population-level (Data-Field 24308) and individual-level (Data-Field 24051) WGS data from the UKB 500K cohort [28]. We used the array data to identify genetic relatedness and to infer identity-by-descent (IBD) states between sibling pairs. In turn, we used the WGS data to identify nucleotide differences between apparent monozygotic twins and to call putative DNMs in trios and siblings.

To align the array data (which relied on hg19) with the WGS data (hg38), we performed quality control and liftover from hg19 to hg38, as described in https://odelaneau.github.io/shapeit5/docs/tutorials/ukb_snp_array. We subsetted the population-level WGS data to the 484,145 individuals who also had array data and individual-level WGS data and limited our analysis to variants that passed the machine-learning-based variant recalibration (DRAGEN-ML) using PLINK with the `plink2 --keep --var-filter` options. All analyses were conducted on the UKB Research Analysis Platform.

#### All of Us

We analyzed individuals in the AoU CDRv8 data release, of which 414,830 had WGS data. We used the Illumina Global Diversity Array capture data provided in the PLINK 1.9 binary format to infer genetic relatedness and to identify IBD segments between sibling pairs. We then relied on the AoU short read whole genome sequencing (srWGS) dataset provided as a Hail VariantDataset (VDS) to examine differences across apparent monozygotic twins and call DNMs in trios and siblings. All analyses were performed on the AoU Researcher Workbench.

### 2 Relatedness inferences

#### UK Biobank

To identify first-degree relatives (kinship ≥ 0.177), we inferred relatedness using the array data (see Methods 1) and KING with the `--related --degree 1` options [78]. Among individuals who also had population-level and individual-level WGS data, we identified 178 apparent monozygotic twin pairs, 5,143 parent-offspring pairs, and 22,358 sibling pairs, categorized based on the *InfType* column of the KING output. We confirmed that all monozygotic twin pairs were of the same sex. For each parent-offspring pair, we used age (Data-Field 21003) to assign the younger individual as the offspring. We then used the parents’ sex (reported in the population-level WGS data) to map each offspring to their father and mother. Finally, we retained only individuals with both parents assigned, resulting in 1, 041 parent-offspring trios. Within this set, we further identified 35 quads for which there were whole genome sequences for both parents and two offspring.

We then identified nuclear families across the parent-offspring trios and sibling pairs as follows. Among the 1,041 parent-offspring trios, 2 involved a monozygotic twin; in these cases, we randomly chose one individual from the twin pair, yielding 1,040 trios. Of these, 970 families had a single offspring (1 trio) and 35 families had two offspring (2 trios), resulting in 1,005 nuclear families. Nuclear families were identified based on shared parentage, and no parent appeared in more than 1 family.

Among the 22,358 sibling pairs, we identified 80 cases in which one of the siblings was also a monozygotic twin of another individual in the dataset or was duplicated. In these cases, we retained only one of the twins/duplicates, which reduced the number of such pairs to 40, and therefore left us with 22,318 sibling pairs in total. Sets of parents were identified by treating sibling pairs as edges in a graph and identifying connected components. These sibling pairs were offspring of 19,893 parents, and for 35 of these, the parental data were also available. Including the 1,005 parents from trios, there are a total of 20,863 parental sets represented in the UKB.

Parental age at birth was estimated using one of two approaches. For parent-offspring pairs with both the parent and offspring present in the cohort, it was calculated as the difference in their years of birth (Data-Field 34). For offspring with no parents in the cohort, it was obtained by subtracting the offspring’s age at recruitment (Data-Field 21003) from the age that they reported for their parents at recruitment (Data-Fields 2946 and 1845), where available. Where parental age was missing for some offspring within a family, it was imputed from a sibling who had reported the parent’s age at recruitment, adjusted for differences in year of birth. If no sibling had reported the parent’s age, parental age was left missing and imputed with the mean of available values within the corresponding biobank where required in downstream analyses (see Methods 13).

#### All of Us

We identified pairs of parents-child or siblings (kinship ≤ 0.354 and ≥ 0.177) and monozygotic twin or genomic duplicate pairs (kinship ≥ 0.354) using the kinship scores calculated by AoU with the Hail pc relate function for all samples with srWGS data [79]. From this set, we removed any pairs involving an individual flagged by AoU in the joint call set QC process [79]. We then subsetted the array data to the remaining parent-offspring or sibling pairs using PLINK 1.9.

To distinguish parent-offspring pairs from siblings, we ran KING with the `--related --degree 1` option and categorized pairs based on the inferred relationship at the *InfType* column of the output file [78, 80]. Some members of parent-offspring or sibling pairs also had a monozygotic twin or duplicate in the sample. In these cases, as described for the UKB, we removed one individual from each monozygotic twin or genomic duplicate pair at random. This step resulted in the removal of 87 sibling and 241 parent-offspring pairs.

We calculated ages using the self-reported birth dates from the AoU Basics Survey [29]. We removed 16 parent-offspring pairs with an age difference of ≤ 10 years and 5 sibling pairs with an age difference of ≥ 30 years, which we suspect might be errors in KING’s relationship inference or incorrect self-reported birth years. For the remaining parent-offspring pairs, we classified the younger person as the child.

These parent-offspring and sibling pairs comprised 7,706 sets of parents. If an individual was a parent in multiple nuclear families (i.e., they had offspring with more than one other individual in our sample), we randomly chose one of the nuclear families (in practice always trios), resulting in a total of 7,700 sets of parents for DNM identification. In the end, we identified 2,242 monozygotic twins or genomic duplicate pairs, 7,692 sibling pairs from 6,735 parents, and 1,292 trios from 1,119 nuclear families. Of these 1,119 nuclear families, 964 had one offspring, 138 had two, 16 had three, and 1 had four. In total, this yields a total of 7,383 unique sets of parents with DNMs in AoU.

We subsetted the AoU srWGS dataset provided as a Hail VariantDataset to these individuals, split multiallelic sites, and densified the VariantDataset to a Hail MatrixTable for all subsequent analyses.

### 3 Allele frequency estimation

#### UK Biobank

We used the population-level WGS data (see Methods 1) to obtain allele counts across the cohort of 439,591 individuals, excluding all monozygotic twin pairs, parent–offspring trios, and sibling pairs (*n* = 44,078); we also excluded first-degree relatives (kinship ≥ 0.177, *n* = 476), identified in Methods 2, using PLINK with the `plink2 --keep --freq` counts options. We refer to this set as “unrelated.”

#### All of Us

In order to avoid densifying a large Hail VDS, we started with the pre-calculated allele counts (number of filter-passing alternative alleles) and allele numbers (total number of filter-passing genotyped alleles) included under the columns *gvs all ac* and *gvs all an* in the AoU Variant Annotation Table (VAT) [81]. Variants with more than 50 alternative alleles were not included in the VAT, and accordingly also excluded from our analyses; the number of excluded variants in the trio and sibling analyses are included in Methods 5 and Methods 8. We then calculated allele counts and allele numbers for all genotype calls that passed the Variant Extract-Train-Score filters (mt.FT is “PASS” or NA) in all twins, siblings, and trios (n = 20,771), by applying the Hail `hl.agg.call_stats` function to the Hail MatrixTable described in Methods 2. Similarly, we calculated allele counts and numbers for all individuals with kinship ≥ 0.177 to any twins/siblings/trios that were not already included in the above set (n = 1,558). We calculated the final allele frequency values in the “unrelated” set of 392,514 individuals by subtracting the allele counts and allele numbers for the twins/siblings/trios and their first-degree relatives from the VAT (total-set of AoU) allele counts and allele numbers.

### 4 Analysis of monozygotic twins and genomic duplicates

Based on our reanalysis of the data from [47], monozygotic twins are expected to differ by approximately five early embryonic mutations on average [47], of which ∼ 1.44 differences remain after retaining only variants with an allele frequency (*V AF*) consistent with 0.5 (i.e., *pbinom*(*V AF* × *DP, DP,* 0.5) *>* 0.05) in the twin carrying the mutation. In turn, duplicate samples should not differ at all, in the absence of sequencing errors. Therefore, almost all differences identified by comparing duplicates and monozygotic twin pairs are errors. With these considerations in mind, we analyzed the genomes of 178 monozygotic twin or genomic duplicate pairs in the UKB and 2,242 pairs in AoU to characterize the error profiles of the two data sets and tune our filters accordingly for subsequent analyses.

To this end, in the UKB, we used the population-level WGS data. For each pair, we generated a list of all autosomal sites at which one individual was heterozygous and the other was homozygous reference (0*/*1 and 0*/*0). We considered each difference between pairs to be an error or mutation in the individual with the heterozygous genotype (the “proband” in what follows). We then applied standard quality control filtering to these differences. Specifically, we required a read depth (*DP*) ≥ 20 and genotype quality (*GQ*) ≥ 30 in the pair, *DP* ≤ 100 and a *V AF* consistent with the expected 0.5 for constitutive mutations (i.e., *pbinom*(*V AF* × *DP, DP,* 0.5) *>* 0.05) in the proband, and 0 reads supporting the alternative allele in the non-proband genome, all obtained from the individual-level WGS data (Methods 1). These steps resulted in an average of 23.9 SNP and 172.4 indel differences per duplicate/monozygotic twin pair.

In AoU, we subsetted the previously described Hail MatrixTable to the individuals in monozygotic twin or genomic duplicate pairs. For each pair, we generated a list of all autosomal sites in the MatrixTable at which one individual was heterozygous and the other was homozygous reference (0*/*1 and 0*/*0). We filtered sites to those that passed the AoU variant hard threshold filters (the column *filters* in the Hail MatrixTable is an empty set). Furthermore, we only kept sites where the genotype calls in both members of the pair passed Variant Extract-Train-Score filtering (mt.FT is “PASS” or NA) and had a GQ ≥ 30 [81]. We were not able to apply filtering steps related to read depth in the analysis of the AoU dataset, given that read depth information is not provided in the AoU srWGS VDS. These steps resulted in an average of 85.8 SNP and 1528.6 indel differences per monozygotic twin or genomic duplicate pair. We removed one twin pair that was a clear outlier (with more than 1 million SNP and indels) in terms of the number of differences; this pair also had the lowest kinship score (0.35401) across all monozygotic twin or genomic duplicate pairs (all other pairs had kinship ≥ 0.456.)

These analyses reveal a clear excess of differences relative to what would be expected from monozygotic twins or duplicates, indicative of sequencing errors. The 96-type mutation spectra of the SNP differences identified in the UKB and AoU were fairly similar to one another, with a cosine similarity of 0.945 (Supplementary Figure 9). Consistent with a large proportion of differences identified in twins and duplicates representing likely sequencing errors, in both data sets, 44.7% of the differences were attributed to signatures previously associated with sequencing artifacts (i.e., SBS27, SBS43, SBS45, SBS46, SBS47, SBS48, SBS49, SBS50, SBS51, SBS52, SBS53, SBS54, SBS55, SBS56, SBS57, SBS58, SBS59, SBS60, SBS95 [38]) (Supplementary Figure 9).

In what follows, we rely on these differences to set our filters in order to decrease spurious calls. In that regard, we note a high error rate of indels (∼ 8-fold more indel differences compared to point mutations, when the mutation rate of indels is ∼ 10-fold lower than that of point mutations [2]) in the UKB and especially AoU (∼ 18-fold more indel differences). We therefore focused all subsequent analyses on SNPs.

To further decrease error rates, we considered that genomic contexts such as repetitive regions may be particularly prone to sequencing artifacts [82]. To identify such regions, we relied on annotations from the Genome in a Bottle Consortium (GIAB), which describe various contexts across the genome for benchmarking purposes, including “difficult” regions that may have higher rates of sequencing errors [82]. We downloaded all version 3.6 stratifications for GRCh38 under “all difficult regions”, “functional technically difficult regions”, and “other difficult regions” from the GIAB FTP [82–86]. We calculated the number of differences between pairs of monozygotic twins or duplicates in the UKB and AoU that fell within each stratification provided by GIAB using the `intersect` function in `bedtoolsr` [87]. If sequencing artifacts were uniform along the genome, the proportion of differences that fall within a stratification should be given by the fraction of the autosomal genome covered by that stratification. For each stratification, we therefore calculated a one-tailed p-value for a binomial distribution using the binom.test function in R with *p* equal to the proportion of autosomal reference genome covered, *n* equal to the number of AoU/UKB SNP monozygotic twin/duplicate differences, *k* equal to the number of AoU/UKB differences that fall within the focal stratification and *alternative* = “greater”. We applied a Benjamini-Hochberg correction to all p-values using the `p.adjust` function in R.

We then excluded any stratification for which the FDR corrected p-value was ≤ 0.05 (Supplementary Table 10), which was true for: 1) all low mappability regions; 2) segmental duplications; 3) tandem repeats with a 5 base pair slop; 4) homopolymers ≥ 7 base pairs and imperfect homopolymers ≥ 11 base pairs with a 5 base pair slop; 5) regions with GC content *<* 25% or *>* 65% with a 50 base pair slop; 6) transcription start sites or first exons that have systematically low coverage; 7) the highly polymorphic killer-cell immunoglobulin-like receptor (KIR) region; 8) collapsed duplications in GRCh38; 9) regions with copy-number variants 10) regions with variants with excess heterozygosity flagged by gnomAD; 11) the highly polymorphic major histocompatibility complex (MHC) region; 12) the highly polymorphic, frequently rearranged VDJ segments of T and B cell receptors; 13) reference assembly contigs smaller than 500 kilobases; and 14) regions within 15 kilobases of a reference gap (Supplementary Table 10). The union of these 14 types of regions covered 20.4% of the autosomal reference genome. Filtering out these regions from a dataset of 180,154 autosomal DNMs identified in 2,976 trios by [32] excluded 17.9% of mutations, roughly as expected under the null. In contrast, considering monozygotic twin/duplicate pairs, 86.0% of identified differences in AoU and 73.9% in the UKB fell in such regions. We therefore filtered out GIAB “difficult regions.”

After applying this filter, the average number of SNP differences between monozygotic twin/duplicate pairs in the remaining ∼ 79.8% of the genome was 12.0 in AoU and 6.2 in UKB; the two-fold higher number in AoU likely reflects slight differences in filtering due to available quality measures. When we again modeled the 96-type mutation spectra as a linear combination of COSMIC SBS signatures [38, 39], the error-associated signature proportion went down to 15.2% in the UKB and 13.0% in AoU, indicating that this filter was effective in removing differences resulting from sequencing errors (Supplementary Figure 9).

Based on these analyses, which suggested additional steps effectively reduce the false positive rate, we applied the same GIAB filters to our analyses of trio and sibling data.

### 5 *De novo* mutations identified in trios

#### Expected number of *de novo* mutations for trios

We calculated the expected number of DNMs per trio based on what was reported in [32]. Specifically, we modeled the number of DNMs in each proband in [32] as a function of parental ages using the R function `glm(number_of_dnms ∼ father_age + mother_age + offset(log(callable_genome)), family = quasipoisson(link = ‘‘log’’))`. We used a quasi-Poisson family to account for mean-variance overdispersion in DNM counts (the dispersion parameter was 2.23). We included an offset of log(2 × 2,682,890,000), which was the denominator used as the length of the callable genome in [32]. We used this model and mean parental ages at birth in AoU and UKB to calculate age-adjusted expected DNM count for the trios in each dataset using the `R predict()` function with `se.fit = TRUE`. We used 2 times the length of the autosomal reference genome after removing difficult regions (see Methods 4) (2 × 2,294,489,308) as the length of the callable genome in these predictions. For the AoU trios, the average paternal age at birth is 30.2 and maternal age is 27.3, and the expected number of DNMs per trio after GIAB filtering is 54.2 (95% CI: 53.8-54.6). For the UKB trios, ages are younger (because of the recruitment criteria for ages): the average paternal age is 24.3 and maternal age is 22.8, leading to an expected number of DNMs per trio after filtering of 47.0 (95% CI: 46.6-47.5).

#### UK Biobank

We subsetted the population-level WGS data to SNPs that did not fall within GIAB stratification (Methods 4, Supplementary Table 10) using PLINK with the `plink2 --snps-only --exclude` options. For each of the 1,040 parent–offspring trios (after excluding 1 trio from which individuals withdrew during the course of our analysis), we generated a list of all autosomal sites at which the offspring was heterozygous (0*/*1) and both parents were homozygous reference (0*/*0). We applied quality control filtering to these differences as described in Methods 4. This pipeline resulted in an average of 46.23 putative SNP differences per offspring. After excluding two outlier trios with *>* 900 SNPs, which appeared to be largely somatic mutations (see Methods 5), the average number of putative SNP DNMs per offspring was 43.45, consistent with the age-adjusted expectation from Methods 5. These trios came from 1,002 nuclear families (see Methods 2).

#### All of Us

For each of the 1,292 trios, we used the Hail MatrixTable to identify all loci for which both parents were homozygous reference (0*/*0) and the child was heterozygous for one alternative allele (0*/*1), then applied quality control filters as previously described for twins/duplicates (Methods 4). After removing GIAB stratifications with high error rates (Methods 4, Supplementary Table 10) using the `intersect` function in `bedtoolsr` [87], the average number of mutations per proband was 56.7. Finally, we removed 37 variants (0.05% of all variants) with more than 50 alternative alleles in the entire AoU cohort, because they were not included in the Variant Annotation Table provided by AoU (Methods 3). The final mean number of mutations per trio was 56.7. These trios came from 1,119 nuclear families (see Methods 2).

#### Outliers

We investigated two trios in the UKB that had highly elevated numbers of putative DNMs (see Methods 5). Because of privacy considerations, we do not provide further details, other than to note that closer inspection of the data suggested that in one case, the elevated count reflects an elevated burden of somatic mutations, and in the other, sequencing artifacts. We excluded both from further consideration.

### 6 Inference of IBD2 segments in sibling pairs

To infer IBD2 segments between sibling pairs in the UKB and AoU, we applied a hidden Markov model (HMM) implemented in the snipar package [27], which is tailored for sibling data and models the joint distribution of genotypes at each SNP conditional on the IBD state. The HMM identifies genomic regions of IBD0, IBD1, and IBD2 under the expected genome-wide proportions of 25% IBD0, 50% IBD1, and 25% IBD2. We used the default transition probabilities based on the deCODE sex-averaged genetic map in GRCh38 coordinates [32].

#### UK Biobank

We used the array data to infer IBD segments shared between all sibling pairs. We ran snipar’s ibd.py on all autosomes, using the KING file generated in Methods 2 as the input to `--king` and an age-and-sex file populated with data obtained from Data-Field 21003 and the population-level WGS data as the input to `--agesex`. The mean genotyping error probability estimated by snipar was 0.000808. We trimmed 0.75 cM from the ends of each IBD2 segment based on the same genetic map and excluded all IBD2 segments with length *<* 6 cM, based on our analysis of quads, as described in Methods 7.

#### All of Us

We split the array data pertaining to all sibling and parent-child pairs as described in Methods 2 into individual chromosomes using PLINK 1.9 [80]. We ran `snipar’s` *ibd.py* on each chromosome with the *–king* input as the KING file described in Methods 2 and *–agesex* populated with self-reported information from the AoU Basics survey. Individuals who did not self-report sex were excluded from the –agesex file, which only affected the genotyping error probability estimation step and not the IBD2 inference. The mean genotyping error probability estimated by snipar was 0.000493. We trimmed 0.75 cM from the ends of each IBD2 segment based on the same genetic map and excluded all IBD2 segments with length *<* 6 cM, as described in Methods 7.

### 7 Analysis of quads

#### IBD2 quality control

We used the 35 quads in the UKB and 192 in AoU to set quality control filters for the IBD2 segment calls made by snipar. Given that the IBD calls are based on sparse genotyping data, the edges of the IBD2 segment calls by snipar may extend too far, to regions that are actually IBD0 or IBD1, and thus include many differences that are not DNMs but inherited variants. Such DNMs will not be included in the DNM calls made using the trio approach. We therefore trimmed the edges of the IBD2 segments. To decide on how broad a region to trim, we calculated (#Sibling DNM calls)/(#Sibling DNM calls also called using trio approach) for 0 cM (i.e., no trim), 0.25 cM, 0.75 cM and 1 cM and considered the false discovery and false negative rates of the sibling approach for each of these choices. In the end, we chose a trim of 0.75 cM (Supplementary Figure 10a), which reduced the average proportion of the autosomal reference genome inferred to be IBD2 between siblings from 24.2% to 22.4% in AoU and 24.1% to 22.4% in the UKB. Putative DNMs in siblings were identified as described in Methods 8.

We further expected that shorter IBD2 segments may be less reliable [88] and therefore determined a minimum length cutoff for IBD2 segments. We calculated the overlap between sibling DNM calls and trio DNM calls for IBD2 segments as described above for multiple choices of minimum lengths. Given the relatively small number of IBD2 segments in the 35 UKB quads, we only used the 191 AoU quads for this analysis, calculating overlap for DNM calls within IBD2 segments of length *<* 2cM (896 sibling DNM calls), 2-4cM (367), 4-6cM (385), 6-10cM (621), 10-20cM (1999), *>* 20cM (7011). In the end, we chose to remove any segment *<*6cM (Supplementary Figure 10b), which reduced the average proportion of the autosomal reference genome inferred to be IBD2 between siblings from 22.4% to 22.2% in AoU and 22.4% to 21.9% in the UKB.

After applying a trim to the IBD2 regions and removing shorter segments, the proportion of sibling differences also identified using the trio approach was 50.8% for the AoU quads and 53.9% for the UKB quads.

#### Distance filter

We observed that DNM calls in siblings not found in trios clustered within the genome more than differences that were also found in trios: within an individual, the distance from one DNM to the nearest DNM was smaller than for DNMs identified by both approaches (Supplementary Figure 11a-b). We suspected that this observation may be the result of clustered errors within individual-specific, error-prone regions. Alternatively, it could arise from mistakes in IBD2 inference that do not fall at the ends of segments, such as in cases where two adjacent IBD2 segments are joined. Yet some mutation types, such asde novo CpG transversions, have been shown to be in clusters of two (and rarely three) variants within distances of up to 20 kb [89]. In order to choose a filter that would exclude errors but minimize the number of true mutations that are missed, we decided to remove clusters with ≥ 3 variants in 100 kb sliding windows (sliding by 1 bp along the genome) (Supplementary Figure 11c). This approach increased the proportion of sibling differences identified using the trio approach to 81.7% for the AoU quads and to 75.2% for the UKB quads. This filter removed 1.1% of the DNMs identified by [32], indicating that it would not lead to a high false negative rate. We did not apply this filter to the DNMs identified using the trio approach, as there is no reason to expect and indeed no evidence that these mutation calls are more clustered than expected from previous trio studies (Supplementary Figure 11d).

#### Allele frequency filter

In addition to DNMs, differences in the genotype calls made for siblings in IBD2 regions may represent sequencing errors or meiotic non-crossover (i.e., gene conversion) events, which lead one sibling to carry an inherited mutation rather than a DNM. Cases in which one sibling is incorrectly called as homozygous reference (0*/*0) but is actually heterozygous (0*/*1) will do the same. In contrast, DNM calls based on the trio approach, which include information about parental genotypes, should not include these two types of errors. On the other hand, trio-based DNM calls may include cases where at least one parent is incorrectly genotyped as homozygous reference (0*/*0) when they are actually heterozygous (0*/*1), while in that case, the sibling-based approach will be unaffected. Mutations identified using both the trio and the sibling approaches in quads will not be affected by any of these three types of errors, and thus should have a lower false positive rate.

Cases in which inherited variants are mistaken for DNMs are likely to involve alternative alleles at higher frequencies than true DNMs (where alternative alleles are only present in unrelated individuals due to recurrent mutations). We therefore applied an allele frequency filter to minimize sequencing errors and non-crossover gene-conversion events (see Methods 3). Using the quad data, we calculated the allele frequencies of variants identified as differences between siblings but not identified using the trio approach; variants identified in trios but not identified using the sibling approach; and those identified using both the trio and the sibling approaches. As expected, the frequencies of the variants identified using both the trio and the sibling approaches were lower than variants found only using siblings or only using a trio, in both the UKB and AoU (Supplementary Figure 12a,c).

Next, we calculated the number of sibling DNM calls also identified using the trio approach divided by the number of sibling DNM calls, for allele frequency filters of 10%, 5%, 1%, 0.1%, 0.01%, and 0.001%. Similarly, we calculated the number of trio DNM calls also called using sibling approach divided by the number of trio DNM calls, for the same set of allele frequency filters. On that basis, we chose an allele frequency cutoff of 0.1% (Supplementary Figure 12b,d), which increased the proportion of sibling DNM calls that overlap with trio calls from 81.7% to 93.3% in AoU, and from 75.2% to 89.7% in the UKB. Of the 6.7% of the sibling calls in AoU that were not made by the trio approach, 61% (95% CI: 58.5%-63.5%) of the alternative alleles were observed in parents prior to the FT and GQ filters (see Methods 4). Of the 10.3% of the sibling calls in UKB that were not made by the trio approach, 37.1% (95% CI: 25.8%-48.5%) of the alternative alleles were observed in parents prior to the DP and GQ filters (see Methods 4). Treating calls made using both trios and siblings as the ground truth, the cutoff of 0.1% led to a 2.9% false negative rate in AoU and 1.3% in the UKB. We did not apply the allele frequency filter to the calls made by the trio approach, given that we would not expect it to substantially improve calls yet it could lead to false negatives; indeed, the overlap with sibling-based calls would barely shift (by 1.2% in AoU and 0.9% in the UKB).

To assess what fraction of putative DNMs identified using the sibling approach may in fact be due to meiotic gene conversion events, we examined the proportion that falls within meiotic double-strand break hotspots, as estimated from DMC1 Chip-Seq, before and after the allele frequency filter [31]. After excluding GIAB difficult regions and applying the 100-kb sliding window filter for clustered errors, we expected that approximately 1.26% (95% CI: 1.21% − 1.32%) of true autosomal DNMs should lie in such DMC1 hotspots, based on trio data in [32]. Before applying our allele frequency filter, we instead estimated that 5.17% (95% CI: 5.08%−5.26%) of all sibling differences in AoU, and 5.46% (95% CI: 5.41% − 5.51%) in the UKB occur in DMC1 hotspots, suggesting that, as expected, our calls also include some recombination events. Reassuringly, the allele frequency filter decreased this proportion to 1.27% (95% CI: 1.22% − 1.31%) in AoU and 1.26% (95% CI: 1.23% − 1.28%) in the UKB, very close to the expectation based on previous trio studies of DNMs [32].

After excluding GIAB difficult regions only, we found that approximately 1.28% (95% CI: 1.22%−1.33%) of true autosomal DNMs in [32] lie in such DMC1 hotspots. In comparison, of the DNMs identified using the trio approach, 1.52% (95% CI: 1.43% − 1.61%) fell within these hotspots in AoU and 1.53% (95% CI: 1.42% − 1.64%) in the UKB.

The resulting DNM calls in siblings after IBD2 quality control, removing clustered mutations, and applying an allele frequency filter showed 89.7% overlap with trio-based calls in the UKB and 93.3% in AoU. The 6.7-10.3% that do not overlap include false positive DNMs in the sibling calls as well as missed DNMs in the trio. We note further that results from the two approaches should not line up perfectly, given that the standard approach to analyzing trios excludes most early embryonic mutations in the parents, whereas calls based on siblings do not [30].

### 8 *De novo* mutations identified from sibling pairs

Having established the validity of our approach in quad families, we next applied it to the full set of sibling pairs in the UKB and AoU, where parental data are not available for the vast majority.

#### Removing sibling pairs with unexpectedly large differences in DNM calls

Given that mutation rates are properties of parents, both siblings could inherit a large number of DNMs, but one sibling is unlikely to inherit a much greater number of DNMs compared to the other, after accounting for their age difference. In principle, this observation could arise, for instance, if a parent was exposed to a mutagen between conceptions, but it is more probable that one of the siblings has high levels of sequencing errors or clonal hematopoiesis. We relied on this consideration to filter out cases with high error rates in one of the siblings. Because the number of DNMs depends linearly on maternal and paternal age [3, 5], older siblings (born to younger parents) are expected to carry fewer DNMs than their younger siblings. At a given paternal and maternal age, the number of mutations inherited by a child is roughly Poisson-distributed [90]. Given that the effect of parental ages is linear with an intercept close to 0 [1], we modeled the difference in the number of mutations between siblings as the difference between two Poisson variables, which follows a Skellam distribution. We approximated the Skellam distribution with a Normal distribution, using the difference in the mean number of putative DNMs between younger and older siblings as the mean, and the sum of these means as the variance. We filtered out data points that deviated more than three standard deviations from the mean for each age difference (Supplementary Figure 1).

#### UK Biobank

We used the subsetted population-level WGS data described in Methods 5. For each of the 22,344 sibling pairs (after excluding 14 pairs where individuals withdrew), we generated a list of all autosomal sites within appropriately trimmed (see Methods 6 and 7) IBD2 segments using PLINK with the `plink2 --exclude` option and focused on sites at which one sibling was heterozygous and the other was homozygous reference (0*/*1 and 0*/*0). We applied quality control filtering to these differences as described in Methods 4. We assigned each single base pair difference between the pair to the person with the heterozygous genotype. This procedure resulted in an average of 44.8 putative SNP differences per sibling pair. We then applied a 100-Kb sliding window and excluded variants in any window containing ≥ 3 differences (Methods 7); this step reduced the number of putative SNP differences to 31.7 per sibling pair. We further removed all variants with allele frequency *>* 0.1% in the unrelated set (Methods 3), resulting in 27.4 putative SNP differences per sibling pair on average. Finally, we removed sibling pairs with a larger difference in DNM calls than expected given their age difference, as described in Methods 8, leading to the exclusion of 684 sibling pairs out of 22,344. In the end, we identified 25.9 putative SNP DNMs per sibling pair on average, across 21,660 pairs. These sibling pairs came from 19,295 nuclear families (see Methods 2), of which 35 overlapped with the parent-offspring families, yielding 19,260 additional nuclear families not represented in the trios.

#### All of Us

For each of the 7,692 sibling pairs, we used the Hail MatrixTable to identify all loci in IBD2 segments at which one sibling was homozygous reference (0*/*0), the other sibling was heterozygous for one alternative allele (0*/*1) and both genotype calls passed quality control filters (Methods 4). We then filtered out GIAB stratifications with high error rates (Methods 4, Supplementary Table 10) using the *intersect* function in *bedtoolsr* [87], resulting in an average number of 59.9 putative DNMs per sibling pair. Next, we applied a 100-kb sliding window and excluded variants in any window containing ≥ 3 differences, which reduced the number of differences to an average of 35.4 per sibling pair. From the total set, we removed 123 variants (0.05% of all variants) with more than 50 alleles in the entire AoU cohort, which were not included in the Variant Annotation Table and likely represent sequencing errors [81] (Methods 3). We further removed all variants with allele frequency *>*0.1% in the unrelated set of 392, 514 individuals, resulting in 29.9 putative DNMs per sibling pair on average. Finally, we removed sibling pairs with a larger difference in DNM calls than expected given their age difference, as described in Methods 8; this led us to exclude 367 sibling pairs and resulted in the identification of 27.0 mutations per sibling pair on average, across 7,325 pairs, involved in 6,735 nuclear families, of which 6,580 were not already included in the trio families.

Building on the error characterization from twin and duplicate differences (see Methods 4), we also excluded clusters of three or more mutations within 100 kb (see Methods 7). Under the assumption that all the remaining differences are errors, our filtering reduced the error rate per bp to 6.75 × 10*^−^*^10^ in the UKB and 1.14 × 10*^−^*^9^ in AoU in the ∼ 79.8% of the autosomal genome that remains accessible. Given mutation rates of 1.28 × 10*^−^*^8^ across siblings in the UKB and 1.34 × 10*^−^*^8^ in AoU, these error rates correspond to 5.3% and 8.5% of DNM calls being errors, respectively.

### 9 Transmission of *de novo* mutation calls

We identified 357 parent–offspring pairs in the UKB and 2,109 in AoU in which DNMs were called in the offspring using the sibling approach (see Methods 2 and Methods 8), and data were available for only one parent. We used these pairs to verify that DNMs identified in the offspring are absent in the parent. Specifically, in the UKB, of the 3,614 DNMs identified across these offspring, 2.2% (95% CI: 1.8% − 2.7%) were observed in the corresponding parent (i.e., with *>* 0 reads supporting the alternative allele prior to DP and GQ filtering; see Methods 4). In AoU, of the 30,268 DNMs identified across these offspring, 2.7% (95% CI: 2.6% − 2.8%) were observed in the corresponding parent (i.e., non-reference genotype call prior to FT and GQ filtering; see Methods 4).

In addition, we identified 489 parent–offspring pairs in the UKB and 892 in AoU in which DNMs were called in the parent using the sibling approach (see Methods 2 and 8). We used these pairs to verify that parental DNMs are transmitted to the offspring with the expected 50% probability. Specifically, in the UKB, of the 5,021 DNMs detected across these parents that had quality-control passing genotype calls in the offspring, 46.3% (95% CI: 44.9% − 47.7%) were transmitted. In AoU, of the 14,139 DNMs detected across these parents that had quality-control passing genotype calls in the offspring, 47.6% (95% CI: 47.2%-48.0%) were transmitted. Under the (conservative) assumption of complete power to detect transmission, these estimates suggest that ∼4.8-7.4% of DNM calls were due to sequencing errors in the sibling (i.e., two times the fraction not transmitted).

Summing up these two sources of error yields an estimated false discovery rate for DNMs based on siblings of 9.7% (95% CI: 6.9%-12.5%) in the UKB and 7.5% (95% CI: 6.5%-8.5%) in AoU.

### 10 Relationship of *de novo* mutation rates to genomic features

As a sanity check on the DNMs called from sibling pairs, we examined the relationship of the DNM rates to features known to influence germline mutation rates: CpG methylation in the male germline and DNA replication timing. To this end, we considered testis CpG methylation levels from ENCODE bisulfite sequencing data (accession ENCFF053FUK) [91, 92], which was previously shown to predict CpG transition rates [5, 36]. We excluded sites overlapping GIAB difficult regions and stratified the remaining CpG sites into five methylation bins (0–20%, 20–40%, 40–60%, 60–80%, and 80–100%) based on the fraction of reads methylated at each site. For each bin, we calculated the CpG*>*TpG mutation rate as the number of CpG*>*TpG DNMs falling within sites in that bin divided by the total number of CpG sites in that bin, normalized per sibling pair in the UKB and AoU, respectively.

We further obtained replication timing estimates from [93], which predict germline mutation rates at the megabase scale, and notably the fraction of C*>*A mutations [36, 94]. Replication timing in 2.5 kb windows was measured in lymphoblastoid cell lines; timing values are Z-scores with a mean of 0 and a standard deviation of 1, with more negative values denoting later replication timing [93]. We generated 1 Mb windows across the genome and calculated mean replication timing in each window. We then divided the windows into ten bins of equal size with increasing replication timing, from earliest (bin 1) to latest (bin 10). For each window, we calculated the C*>*A mutation opportunity as the count of C (or G on the other strand) within the window after masking positions overlapping GIAB difficult regions in the GRCh38 reference genome. We calculated the C*>*A DNM rate per window normalized per sibling pair in the UKB and AoU, respectively.

To assess whether the genomic distribution of DNMs across replication timing differed between *REV1* carriers and non-carriers, we annotated each DNM in *REV1* carrier (for which the number of DNMs = 874) and non-carrier families (DNMs = 873,674) with replication timing estimates from the source described above. 98.5% of DNMs in carriers and 98.9% in non-carriers had fell within the 2.5 kb windows with replication timing information. We then carried out a two-sided Kolmogorov-Smirnov test using the `ks.test()` function in R (p = 0.16). Similarly, to test whether CpG methylation differed between *LIG1* carriers and non-carriers, we annotated each CpG*>*TpG mutation in *LIG1* carrier (for which the number of DNMs = 609) and non-carrier families (DNMs = 157,898) with testis CpG methylation levels from the source described above. 96.9% of DNMs in carriers and 96.8% in non-carriers had methylation information available. The two-sided Kolmogorov-Smirnov test had a p-value of 0.27 (Supplementary Figure 8).

### 11 Comparison of germline mutations across genetic ancestries

#### Genetic ancestry classifications

Since we were analyzing two different population cohorts and wanted to assess how their genetic ancestries were drawn from worldwide structure, we projected UKB and AoU samples onto a common principal component (PC) space, generated using HGDP-1KG samples [40]. To this end, we identified autosomal, biallelic SNPs with allele frequency *>*0.1% and call rate *>*99%, which we LD-pruned using a cutoff r2 of 0.1 across all three datasets. 130,660 sites that passed these filters in AoU and HGDP-1KG were provided by AoU in a sites-only VCF format [79]. Of these SNPs, 130,623 were also in the UKB, passed DRAGEN-ML filtering, and 110,028 were again biallelic. We retained variants with allele frequency *>*0.1% and call rate *>*99% in the entire UKB population-level WGS dataset using `plink2 --maf 0.001 --geno 0.01`, which resulted in 62,738 SNPs. Finally, we LD-pruned these SNPs with a cutoff r2 of 0.1 in the entire UKB population-level WGS dataset using `plink2 --indep-pairwise 50 5 0.1`, resulting in 57,098 SNPs. For AoU, we worked with the provided Hail MatrixTable, which included all variants with “population-specific allele frequency (AF) *>*1% OR population-specific allele count (AC) *>*100, in any computed ancestry subpopulations” (ACAF) based on AoU’s own PCA and ancestry classification [79]. This matrix table included 57,025 of the above SNPs, which we used as our final set of SNPs for PCA.

Next, we subsetted the HGDP-1KG srWGS callset in dense MatrixTable format provided by gnomAD 3.1.2 to the 3400 QC-passing and unrelated individuals used in gnomAD’s PCA (samples were provided in gnomAD release 3.1 under secondary analyses) [72]. We ran PCA using Hail’s hwe normalized pca with *k* = 20 (variant loadings can be found in Supplementary Table S2). We projected all AoU and UKB samples onto this PC-space by applying Hail’s `experimental.pc project` to the entire UKB population-level WGS dataset converted to Hail MatrixTable format and the ACAF Hail MatrixTable provided by AoU (described above), respectively. For all subsequent analyses, we used the first six PCs, which together explained 82.6% of the variance explained by the 20 PCs (Supplementary Figure 13a).

For the purpose of assigning individuals to genetic ancestry groups, we used k-means clustering on the HGDP-1KG PCs. We calculated average silhouette scores for *k* = 1 through 15 with 1000 random starts each using R’s kmeans function and the silhouette function from the R package cluster [95]. The k with the highest silhouette score was 10 (Supplementary Figure 13b). The HGDP and 1KG population labels for each HGDP and 1KG sample across the 10 clusters can be found in Supplementary Figure 14. Membership in cluster 1 is entirely drawn from the EAS (East Asian) continental label in gnomAD [40]. Similarly, cluster 2 corresponds to the AFR (African/African American) label, cluster 3 the MID (Middle Eastern) label, cluster 7 the SAS (South Asian) label, and cluster 10 the OCE (Oceanian) label. Clusters 4 and 6 are comprised of individuals with the AMR (Admixed American) label, with cluster 4 corresponding to individuals closer to the PEL (Peruvian in Lima) label in 1KG, and cluster 6 the CLM (Colombian in Medellin) and and PUR (Puerto Rican in Puerto Rico) labels. We thus refer to Clusters 4 and 6 as AMR1 and AMR2. Clusters 5, 8, and 9 include individuals with the EUR label. Cluster 5 mainly corresponds to the IBS (Iberian populations in Spain) and TSI (Toscani in Italia) labels in 1KG, cluster 8 corresponds to the FIN (Finnish) and Russian labels, and cluster 9 is mainly represented by CEU (Northern Europeans from Utah) and GBR (British from England and Scotland). We thus refer to Clusters 5, 8, and 9 as EUR1, EUR2 and EUR3.

Next, we assigned 40,645 offspring (from trios or sibling pairs) in the UKB and 14,190 from AoU to these clusters. We calculated the squared Euclidean distance of each of the chosen offspring to the centroid of every cluster and assigned each individual to the closest genetic ancestry cluster. For the purposes of our analyses, we removed genomes in UKB and AoU with distance greater than the 95th percentile of HGDP-1KG distances for each cluster, thus likely excluding individuals who draw substantial recent ancestry from more than one of these clusters. By this approach, 99.1% of UKB and 86.7% of AoU offspring were assigned genetic ancestries (Supplementary Figure 15).

#### Comparison of germline mutation rates across genetic ancestries

When comparing DNM rates across ancestry groups, we relied on data from both sibling pairs and trios. In the small subset of cases in which there was more than one sibling pair or offspring of the same parents, we considered each offspring as an independent data point (as did [15]). For trios, the length of the callable genome was calculated as described in Methods 5. For each individual in a sibling pair, the length of the callable genome was half of what was calculated for the pair in Methods 5. If an individual was a member of multiple sibling pairs, we kept the sibling differences for the pair with the largest callable IBD2 segments. In considering the impact of ancestry on mutation rates, we included three covariates: parental ages (when available), biobank and source (siblings or trios).

We first tested for differences across ancestries in trios only, where we had both paternal and maternal ages. We included ancestry groups with ≥ 60 individuals, which were AFR (*n* = 137), AMR1 (*n* = 82), EUR1 (*n* = 130), AMR2 (*n* = 294) and EUR3 (*n* = 1, 446). We tested for multiplicative differences in germline mutation rates across ancestries, accounting for batch effects (i.e., we included an indicator variable for biobank) using a quasi-Poisson family (due to mean-variance overdispersion in our DNM counts) and the log link with the R function `glm(number_of_dnms ∼ father_age + mother_age + biobank + cluster + offset(log(callable_genome)), family = quasipoisson(link=‘‘log’’))`. We included the length of the callable genome as an offset to model the Poisson rate parameter as mutation rate per bp rather than the total genome-wide DNM count. We assessed the contribution of each variable using single-term deletion (Type III) analysis of deviance with the drop1() function in R. Effects of father’s age at conception, mother’s age at conception and biobank were significantly different from 0 at the 5% level (F = 289.4, 36.5, 101.7 and p = 7.1 × 10*^−^*^61^, 1.8 × 10*^−^*^9^, 2.2 × 10*^−^*^23^, df = 1) whereas that of cluster was not (F = 0.69, p = 0.60, df = 4). In particular, samples from the UKB had an 11% lower mutation rate than samples from the AoU for the same mean parental ages. We used this model to generate predicted numbers of DNMs for each cluster at the sample mean paternal age (27.2 years), maternal age (25.0 years), and biobank set to UKB for the diploid autosomal genome using the `R predict()` function with `se.fit = TRUE`. The 95% confidence intervals for the clusters overlapped among ancestry groups, consistent with the non-significant F-test for cluster (Supplementary Figure 16).

Next, we tested for differences across ancestries in individuals from both trios and siblings with paternal age information. We included paternal but not maternal ages to maximize the number of individuals that we could analyze and because 75-80% of germline mutations are paternal in origin [1]. We again included clusters which we had ≥ 60 individuals, which were EAS (*n* = 64), AFR (*n* = 282), AMR1 (*n* = 97), EUR1 (*n* = 283), AMR2 (*n* = 350), SAS (*n* = 137) and EUR3 (*n* = 9,293). We used the same approach as above with a slightly modified model `glm(number_of_dnms ∼ father_age + source + biobank + cluster + offset(log(callable_genome)), family = quasipoisson(link=‘‘log’’)`) where the variable `maternal_age` is not included and the variable source is an indicator that equals 0 if the individual came from a sibling pair and 1 if they came from a trio. Effects of father’s age, biobank, and source were significant at the 5% level (F = 2,519.5, 177.5, 131.1 and p = *<* 10*^−^*^100^, 3.6 × 10*^−^*^40^, 3.6 × 10*^−^*^30^, df = 1) whereas that of cluster was not (F = 1.59, p = 0.15, df = 6). As before, samples from the UKB had an 11% lower mutation rate on average compared to those in the AoU at the same mean paternal age and for the same source. Moreover, samples at the same mean paternal age, cluster, and biobank that were based on trios had a 7.4% lower mutation rate compared to those from siblings. These differences could result from a number of possible sources, including a lower false positive rate in the trios compared to the siblings, the inclusion of some gene conversion events among sibling differences but not among those based on trios, or from true biological differences, such as a lower mean maternal age in trios compared to mothers of siblings (e.g. UKB trios have a mean maternal age of 22.8 and siblings 27.7) or the inclusion of early embryonic mutations in sibling differences but not trios [30].

As noted in the main text, [22] inferred based on substitution data that the mutation rate per year is roughly 5% higher per year in non-Africans compared to Africans. We compared mutation rates in AFR (*n* = 282) vs. Non-AFR (*n* = 10,234), accounting for paternal ages and batch effects using both trios and siblings (where “Non-AFR” pools all other ancestry groups). Samples in the Non-AFR group had a 2.4% higher germline mutation rate per generation than those in the AFR one (*p* = 0.06, Supplementary Figure 3). Rerunning the test using all samples (AFR *n* = 2,983 and non-AFR *n* = 49,606), even those for which we do not have parental age information (and therefore uncorrected for paternal age), the qualitative result is the same: the mutation rate is 4.7% higher in non-Africans than Africans.

At the same age of reproduction, a per year difference of 5% is equivalent to a 5% difference in the number of DNMs. Under this assumption, the probability of observing a difference of ≤ 2.4% is 2.3%. This estimate was obtained as follows: the standard error for the coefficient of the cluster indicator in our model was 0.0126, when under the alternative model, we expect that *β̂* ∼ *N* (log(1.05), 0.0126^2^). The probability of observing an effect such as we estimated or smaller is thus Pr(*β̂* ≤ log(1.024)) = 0.023). Moreover, given a standard error of 0.0126, the smallest effect we have 90% power to detect is a ±4.2% difference between AFR and non-AFR. Power was estimated from (*z*_0.975_ + *z*_0.90_) × *SE* = (1.960 + 1.286) × 0.0126 = 0.04084 and exp(0.04084) = 1.042.

#### Comparison of germline mutation spectra across genetic ancestries

For this analysis, we used individuals from both trios and siblings with paternal age information and clusters with ≥ 60 individuals, as described in Methods 11. We generated mutation spectra for each individual by categorizing their DNMs into seven mutation types representing strand-collapsed C*>*X and T*>*X mutations, with C*>*T mutations split depending on whether it is in a CpG context.

We tested for differences in the proportion of mutation type *j* across ancestries, accounting for batch effects using `glm((mutation_type_j, total_mutation_count) ∼ father_age + source + biobank + cluster, family = binomial(link = ‘‘log’’))`. We used single-term deletion likelihood-ratio tests to assess whether each predictor improved the fit using the `drop1()` function in R. We applied FDR correction for each predictor separately (seven tests per predictor). The results of these tests are in Supplementary Table 11. For visualization, we evaluated the fitted model on the response scale at the reference setting `father_age = 29, biobank = ukb and source = sibling` (Figure 3, Supplementary Figure 4).

Mutation type proportions differed as a function of source (Supplementary Table 11), possibly owing to unmodeled maternal age differences across sources and subtle influences of quality control steps interacting with mutation types. Moreover, the difference detected between trios and sibling pairs could reflect the imposition of an allele frequency filter for siblings but not trios, the effect of which will depend on the mutation type. Alternatively, it may stem from a greater contribution of early-embryonic mutations to the DNM calls in siblings than in trios [30].

Ancestry group was a significant predictor for C*>*T mutations after FDR correction (Supplementary Table 11). To identify the ancestry clusters that had significantly different proportions of C*>*T, we formed all pairwise ancestry contrasts via `pairs(emm, adjust=‘‘none’’, weights=‘‘proportional’’)`. Finally, we applied FDR correction to all tests (28 ancestry pair comparisons) using `p.adjust(p.value, method=‘‘fdr’’)`. We found a higher proportion of C*>*T mutations in AMR1 compared to EUR1 and EUR3 (Supplementary Table 1).

In principle, differences in maternal ages across ancestry groups could contribute to shifts in the spectra. Another source of difference among ancestries could be the allele frequency filter that we imposed on the DNMs called by the sibling approach, given that the proportion of true mutations excluded by this filter will depend on demographic history and the mutation rate of each specific type [41]. To ensure that the differences in C*>*T were not a result of unmodeled differences in maternal age or the allele frequency filter that we imposed on DNM calls in siblings, we tested if ancestry was a significant predictor for C*>*T mutations using trio DNMs only, including maternal age as a covariate. Even with the much smaller data set, cluster remained a significant predictor for C*>*T proportions (p = 0.007).

To home in on specific contexts that drive the observed C*>*T differences, we used mutation classifications that incorporate adjacent bases [1], and generated one 96-type mutation spectrum per cluster by combining all DNMs that came from individuals assigned to that cluster. To maximize the number of DNMs that we can compare without batch effects, we carried out tests using only AoU sibling DNMs (Supplementary Table 2). We then tested for differences in the two pairwise comparisons for C*>*T mutations that achieved statistical significance in the regression-based tests (Supplementary Table 1). We compared the resulting mutation spectra using the method described in [24], which uses ordered Chi-square tests to test for differences in counts across each triplet-context mutation type, accounting for the fact that mutation types are not independent from each other. We applied FDR correction to all tests using `p.adjust(p.value, method=‘‘fdr’’)`. None of the trinucleotide pairwise comparisons were significant after FDR correction; instead, the ancestry differences that we identified at C*>*T mutations appear to be distributed across triplet contexts.

To test if there is an enrichment of TCC*>*TTC in EUR, we again used AoU sibling DNMs. We generated a 96-type mutation spectrum by combining all AoU sibling DNMs that came from individuals assigned to EUR1 and EUR3, and another spectrum combining individuals labeled EAS and AFR. The proportion of TCC*>*TTC was higher in the EAS and AFR spectrum relative to EUR, and the comparison was not significant (p = 0.08).

For the comparison of CCG*>*CTG in SAS and EUR3, we used UKB siblings DNMs, as this maximized the number of DNMs from each ancestry group while avoiding batch effects (Supplementary Table 2). We generated 96-type mutation spectra by combining all UKB sibling DNMs that came from individuals assigned to the two ancestry groups, respectively.

### 12 The effect of smoking on germline mutations

#### Testing the effect of smoking on germline mutation rates

For this analysis, we used offspring from trios in AoU and the UKB. For the UKB, we generated the paternal and maternal smoking status by marking smoking as “Yes” if the UKB Data-Field 20116 was “Previous” or “Current”, “No” if it was “Never” and NA otherwise. For the AoU, we generated paternal and maternal smoking statuses by marking smoking as “Yes” if the answer to question concept ID 1585857 in the AoU Lifestyle Survey (smoked 100 cigarettes lifetime) was “Yes”, “No” if the answer was “No” and NA otherwise. We kept trios where both parents had a non-NA smoking status. For offspring sharing the same parents, we collapsed records into a single row per family, averaging DNM counts and parental ages. This resulted in DNM phenotypes from 1, 058 sets of parents from AoU and 984 from the UKB.

Since our DNMs are not phased to the maternal or paternal germline, we tested for a difference in germline mutation rates between parental sets in which neither smoked or both smoked, accounting for batch effects. Using a similar model as the one described above (Methods 11), we included an indicator both parents smoked. While father’s age, mother’s age and biobank improved the model fit (F = 204.8, 29.6, 103.8, p = 5.0 × 10*^−^*^43^, 6.4 × 10*^−^*^8^ and 1.9 × 10*^−^*^23^, df = 1), smoking status did not (F = 2.4 and p = 0.12, df = 1). Although the point estimate for smokers is higher, the 95% confidence intervals for the predicted number of DNMs at the sample mean paternal age (27.5), maternal age (25.3), and biobank set to UKB overlapped (Supplementary Figure 17). Since most germline mutations are paternal in origin [3], we also tested for a difference in DNM rates as a function of paternal smoking status, accounting for batch effects. As above, while father’s age, mother’s age and biobank improved the model fit, paternal smoking status did not (F = 2.0 and p = 0.16, df = 1).

We further tested whether the known tobacco-associated COSMIC SBS signatures (SBS4, SBS29, SBS92, SBS100, SBS109) contributed to germline mutation spectra as a function of smoking status. We generated two 96-type mutation spectra comprised of the DNMs from trios with parents that both smoked (13,309 DNMs) and with parents that did not (12,341 DNMs). We then modeled each 96-type mutation spectrum as a linear combination of COSMIC SBS signatures [39]. None of the signatures were found to contribute to mutations in smokers (or non-smokers) (*<* 0.01% contribution).

A previous study based on 9,033 trios [15] reported a 2.4% increase in the total (unphased) germline mutation rate when both parents were smokers compared to when both were not. We estimated the power of our test to detect an increase in the mutation rate of this size or greater. Under the null hypothesis, the Wald statistic can be approximated as a normal distribution *Z_H_*_0_ ∼ *N* (0, 1). The standard error for the coefficient of the smoking indicator in our model was 0.0099, meaning that, under the alternative hypothesis, the Wald statistic can be approximated as *Z_H_*_1_ ∼ *N* (log(1.024), 0.0099^2^). For a one-sided test at an alpha of 0.05, Power = Pr(*Z_H_*_1_ ≥ *z_0.95_*) = 0.77.

### 13 Identification of mutator alleles

To test for the presence of mutator alleles, we identified deleterious variants across 210 genes involved in DNA repair and maintenance [42] and obtained or inferred parental genotypes for 20,262 families in the UKB and 7,383 families in AoU (see Methods 5 and Methods 8). Because some genes overlapped extensively with GIAB regions, we excluded those with more than 33.5% overlap. This cutoff was defined using the Tukey fence criterion [96], calculated as the 75th percentile of the overlap distribution across genes plus 1.5 times the interquartile range, leaving 180 genes for burden testing. We then tested for associations with the total mutation rate and type-specific mutation rates (defined as the rates of CpG*>*TpG, C*>*G, non-CpG C*>*T, T*>*C, and C*>*A mutations).

#### Deleterious variants

In the UKB, we subsetted the population-level WGS data (see Methods 1) to variants that had a missing call rate ≤ 1% and fell within our chosen gene set using PLINK with the `plink2 --geno --extract` options. Gene coordinates were obtained from Ensembl BioMart and provided as input to `--extract`. Canonical functional consequences for the resulting variants were annotated using Ensembl VEP, implemented via HAIL-VEP on the UKB RAP. We identified 13,557 LoF and 7,189 deleterious missense variants across the 180 genes, with variants labeled as LoF if their predicted consequences contained transcript ablation, splice acceptor variant, splice donor variant, stop gained, or frameshift variant, and as deleterious missense if their predicted consequences contained missense variant and had a REVEL score *>* 0.75 [45]. We further excluded predicted LoF (pLoF) variants with allele frequency *>* 1%, as common pLoF variants are enriched for annotation artifacts [43, 44], resulting in 13,550 LoF variants, 6,429 SNPs and 7,121 indels (Supplementary Figure 18) across the UKB cohort.

In AoU, we subsetted the VAT to where the column gene id matched any of the 180 genes, is_canonical_transcript was true, and gvs_all_an was *>* 821,364 (corresponding to missing call rate ≤ 1%). We identified 7,870 deleterious missense variants, where the VAT column `consequence` was missense variant and `revel` was *>* 0.75. Moreover, we identified 14,740 LoF variants, where the VAT column consequence was equal to any of transcript ablation, splice acceptor variant, splice donor variant, stop gained, or frameshift variant. After removing LoF variants for which the column `gvs_all_af` was ≤ 1%, we were left with 14,729 LoF variants, of which 6,715 were SNPs and 8,014 indels (Supplementary Figure 18) across the AoU cohort.

#### Sum of parental genotypes

Where parental genotypes were available, we directly obtained the sum of parental genotypes at LoF and deleterious missense variants, with alleles encoded as 0 (reference) and 1 (alternative), so that the sum represents the total number of alternative alleles carried by both parents. Where parental genotypes were not available, we inferred that sum using the genotypes of sibling pairs and their local IBD state, following the approach of [27]. Specifically, for each variant and sibling pair, we first determined whether the variant fell within an IBD0, IBD1, or IBD2 segment between the two siblings (see Methods 6, prior to trimming), and the inference proceeded as follows:

##### IBD0

In IBD0 segments, the siblings share no parental haplotypes and therefore collectively carry all four parental alleles. The sum of parental genotypes is thus fully observed in the offspring and equals the sum of the two sibling genotypes.

##### IBD1

In IBD1 segments, the siblings share exactly one parental haplotype and jointly reveal three of the four parental alleles. If the sibling genotypes differed by two alleles (e.g., 0/0 and 1/1), the configuration is inconsistent with IBD1, and inference proceeded as described in Methods 13. Otherwise, the identity of the three observed parental alleles depends on the sibling genotype combination: if both siblings were homozygous reference (e.g., 0/0 and 0/0), all three observed parental alleles were reference; if one sibling was heterozygous and the other homozygous reference (e.g., 0/1 and 0/0), the observed alleles were two reference and one alternate; if one sibling was heterozygous and the other homozygous alternate (e.g., 0/1 and 1/1), the observed alleles were two alternate and one reference; and if both siblings were homozygous alternate (e.g., 1/1 and 1/1), all three observed alleles were alternate. The fourth parental allele was imputed using the cohort-wide allele frequency of the alternative allele *q*.

When both siblings were heterozygous (e.g., 0/1 and 0/1), three parental alleles were observed, but, without phasing information, it is ambiguous which allele is shared IBD. In this case, we estimated the sum of parental genotypes as the weighted average over the two possible shared alleles, conditional on the sibling genotypes and the IBD1 state. Following [27], the alternative allele with frequency *q* is shared with probability 1 − *q*, and the reference allele with frequency 1 − *q* is shared with probability *q*. Taking the weighted average over these two possibilities yields (1−*q*)(1+*q*)+*q*(2+*q*) = 1+2*q*.

##### IBD2

In IBD2 segments, the siblings share both parental haplotypes and therefore carry only two of the four parental alleles. If in practice the sibling genotypes differ, the configuration is inconsistent with IBD2, and the inference proceeded as described in Methods 13. Otherwise, the two observed alleles determine half of the sum of parental genotypes, and the remaining two alleles were imputed, i.e., the expected parental genotype sum was set to the shared sibling genotype plus 2*q* [27].

##### Missing or inconsistent IBD inference

When the IBD inference was missing or inconsistent, we imputed the sum of parental genotypes following the procedure laid out in [27] (Supplementary Table 12).

To infer the sum of parental genotypes at a variant in families with more than one sibling pair, we first prioritized pairs that were IBD0 at the site. If no IBD0 pair was available, we considered IBD1 and IBD2 pairs, selecting those that inherited the highest number of parental alternative alleles; in the case of ties, we prioritized IBD1 over IBD2. If no IBD0, IBD1, or IBD2 pairs were available, we selected pairs with missing or inconsistent IBD inference. Finally, if multiple pairs remained, we chose the pair with the maximum inferred sum.

The sums of parental genotypes at individual variants were aggregated across all LoF and deleterious missense variants within each gene to generate gene-level sums of parental genotypes. On average, parent sets carry 1.14 LoF and 0.66 deleterious missense variants across the set of 180 genes.

#### Validation of parental genotype imputation

To assess the accuracy of the imputation approach, we validated it in 155 AoU families and 35 UKB families containing parents and two or more offspring, in which we could both observe and impute sums of parental genotypes. Specifically, we defined putative mutator carriers as families in which the sum of parental LoF or deleterious missense genotypes at a gene is greater than or equal to 1, and non-carriers as those with a sum less than 1. For each gene, we compared the carrier status observed directly in the parents to the status inferred from the siblings.

In AoU, the false positive rate was only 0.01% (95% CI: 0.00% − 0.02%) and the false negative rate 21.4% (95% CI: 17.0% − 25.8%), for LoF carriers, and 0.01% (95% CI: 0.00%−0.01%) and 13.8% (95% CI: 7.3%−20.3%) for deleterious missense carriers, respectively (Supplementary Table 13). In the UKB, where we had fewer families on which to base these estimates, the false positive and false negative rates were 0.02% (95% CI: 0.00%−0.05%) and 36.8% (95% CI: 15.1%−58.5%) for LoF carriers, and 0% and 28.6% (95% CI: 8.6% − 48.6%) for deleterious missense carriers (Supplementary Table 13).

#### SBS comparisons

To quantify shifts in the contributions of COSMIC SBS signatures [38] in carriers and non-carriers (see Methods 13) in a set of genes previously shown to affect somatic mutation rates (*BRCA1*, *BRCA2*, *EXO1*, *PALB2*, *PAXIP1*, *RIF1*, *MLH1*), we performed signature decomposition using non-negative least squares (NNLS). We analyzed a set of COSMIC SBS signatures, including SBS5, SBS1, SBS39, SBS12, SBS16, and SBS30, the mutational signatures reported in the germline [37, 39], as well as SBS3, SBS6, SBS14, SBS15, SBS20, SBS21, SBS26, SBS44, SBS10a, SBS10b, SBS10c, SBS10d, SBS18, and SBS36, which are associated with DNA damage, repair, and polymerase proofreading [38]. For each group (i.e., carriers and non-carriers), the merged 96-type mutational spectrum (constructed from distinct DNMs per family) was approximated as a non-negative linear combination of these signatures, with NNLS used to estimate the contribution of each signature (`scipy.optimize.nnls`).

To test for shifts from the dominant germline signatures to signatures associated with DNA damage, repair, and polymerase proofreading, we compared the fraction of mutations assigned to signatures of interest between carriers and non-carriers. For each group, we calculated the contribution of one or more signatures divided by the total contributions of all signatures estimated by NNLS. To assess significance, carrier labels were randomly reassigned within biobank and source strata (UKB trios, UKB sibling pairs, AoU trios, or AoU sibling pairs) 1,000 times, preserving the observed number of carriers in each stratum. For each permutation, the difference in fractions between the two groups was calculated. The empirical p-value for enrichment was then obtained as the proportion of permutations in which the permuted difference was greater than or equal to the observed difference using the standard +1 adjustment for the numerator and denominator (Supplementary Table 3).

We took a similar approach for the two top genes identified in our burden tests (Methods 13): *REV1* and *LIG1* (Supplementary Tables 7 and 8). In this case, however, we reported two-tailed p-values, calculated as the proportion of permutations whose absolute difference was greater than or equal to the absolute value of the observed difference. Results were similar if we divided by the total number of mutations rather than only those assigned to signatures (Supplementary Tables 7 and 8). We also tried a different signature assignment method, based on the expectation–maximization framework described by [39]; to implement it, we considered all COSMIC SBS signatures without regularization (*β* = 0). This procedure led to similar results (Supplementary Tables 14 and 15).

#### Burden tests

For parent-offspring trios, we estimated mutation rates using the number of DNMs in the offspring as the numerator and the genome length (adjusted for GIAB, Methods 4) multiplied by 2 as the denominator. Estimates of parental rates were obtained by averaging across all offspring in the family. For sibling pairs, rates were calculated using the number of DNMs in the two siblings as the numerator and the length of their IBD2 segments (adjusted for GIAB) multiplied by 4 as the denominator. Estimates of parental rates were again obtained by averaging across all sibling pairs in the family. We estimated type-specific parental rates analogously for all 7 mutation types defined above, using the number of DNMs of the corresponding type in the offspring or the two siblings as the numerator, and the denominator defined above, multiplied by the fraction of sites representing the corresponding mutational opportunities genome-wide (i.e., the fraction of sites that are CpG dinucleotides on either strand for CpG*>*TpG, C for C*>*G and C*>*A, C not in CpG dinucleotides for non-CpG C*>*T, and T for T*>*C). Adjusted for GIAB, these comprise 1.7% of sites for CpG*>*TpG, 39.5% for C*>*G and C*>*A, 37.8% for non-CpG C*>*T, and 60.5% for T*>*C.

We conducted burden tests to assess associations between parental total and type-specific mutation rates and gene-level sums of parental genotypes at LoF and deleterious missense variants (see Methods 13). For each gene, we fit a linear model using ordinary least squares (OLS) with heteroscedasticity-robust standard errors (HC3) using `statsmodels.regression.linear_model.OLS`, with mutation rate as the response variable. Because of power considerations, we only considered 138 genes with at least 20 carriers. Under an *additive* model, the predictor of interest was the sum of parental LoF and deleterious missense genotypes at the gene for each family, capped at 2 (i.e., all sums greater than 2 were set to 2). All models included the first six principal components (see Methods 11), computed from one offspring in families derived from parent-offspring trios and from one sibling in families derived from sibling pairs, as covariates, to control for population structure. We also included paternal and maternal ages (see Methods 2); when these were unavailable, we imputed them using the mean of the available values within the corresponding biobank and source (UKB trios, UKB sibling pairs, AoU trios, or AoU sibling pairs). We note that the parental ages for siblings in AoU were based on cases in which the parent also participated in the cohort. Thus, the mean parental age at birth used for imputation is likely to be somewhat lower than the true mean age of all sibling pairs. This effect should be negligible for the UKB, in which only 2% of the parental ages come from cases in which the parent is also included in the dataset. Finally, we included a binary indicator variable for the biobank (UKB or AoU) and source (trios or sibling pairs) to control for batch effects. To test for an *increase* in mutation rate associated with parental genotypes, we used one-tailed p-values derived from `statsmodels.regression.linear_model.OLSResults.pvalues`, and applied a Benjamini–Hochberg FDR correction using `statsmodels.stats.multitest.multipletests` across the 138 genes, separately for each phenotype (Supplementary Table S1).

To control for residual inflation in test statistics, we also applied genomic control [68], and report the resulting p-values and q-values in Supplementary Table S1.

We additionally approximated a *recessive* model by redefining carriers as families in which the parental genotype sum at a gene is greater than or equal to 2, and non-carriers as those with a sum less than 2, and used this binary indicator as the predictor of interest in place of the parental genotype sum. Note that this approach will result in misclassifications, as a sum ≥ 2 can reflect either a homozygous or compound heterozygous parent, or two heterozygous parents, and we cannot distinguish between these cases. Due to the rarity of such carriers in this cohort, only 11 genes had at least 20 carriers and were thus included in the burden tests. All covariates and significance estimation procedures were as described above (Supplementary Table S1).

For the top associations, we performed three additional validation analyses (Supplementary Table 5). First, to examine whether the two annotations have similar effects on mutation phenotypes, we included parental genotype sums at LoF and deleterious missense variants as separate predictors and tested whether their coefficients differed significantly using an F-test (`statsmodels.regression.linear_model.OLSResults.f_test`; Supplementary Table 6). Second, to assess consistency across biobanks, we included a parental genotype sum × biobank (UKB or AoU) inter-action term in the model and obtained the interaction p-value from `statsmodels.regression.linear_model.OLSResults.pvalues` (Supplementary Table 9). Finally, to assess whether the associations might reflect clonal hematopoiesis rather than germline effects, we identified families in which at least one sibling carried a disruptive variant in *REV1* (16 families) or *LIG1* (57 families) and at least one did not, and compared mutation rates between carrier and non-carrier siblings using a two-sided Mann–Whitney U test (scipy.stats.mannwhitneyu). The resulting p-values were 0.4975 for *REV1* and 0.1346 for *LIG1*.

## Data and Code Availability

Variant loadings for PCA are included in Supplementary Table S2. Our pipeline to call *de novo* mutations as well as scripts used to run downstream analyses will be made available in a community workspace within the AoU Researcher Workbench. Scripts used to run analyses on UK Biobank data will be made available on GitHub and returned to UK Biobank in accordance with their data return policy.

## Acknowledgments

We thank Doc Edge, Will Milligan, Hakhamanesh Mostafavi, Simon Myers, Pier Palamara, Guy Sella, Simon Tavaŕe, and members of the Przeworski lab for helpful discussions and Will Milligan and Guy Sella for comments on a draft of the manuscript. This work was supported by R35 GM083098 and a Columbia University Research Initiative in Science and Engineering award to MP. This research has been conducted using the UK Biobank Resource under Application Number 11138. We thank the participants of the UK Biobank and the UK Biobank team for making available the data used in this study. We thank the participants of All of Us and the National Institutes of Health’s All of Us Research Program for making available the data used in this study.

## Declaration of generative AI and AI-assisted technologies in the manuscript preparation process

During the preparation of this work, the author(s) used Claude (Anthropic) for programming-related tasks (including code for figure generation), literature search, and for revising grammar and readability. The author(s) reviewed and edited the content as needed and take full responsibility for the content of the published article.

## Supplementary Materials

**Supplementary Figure 1.**
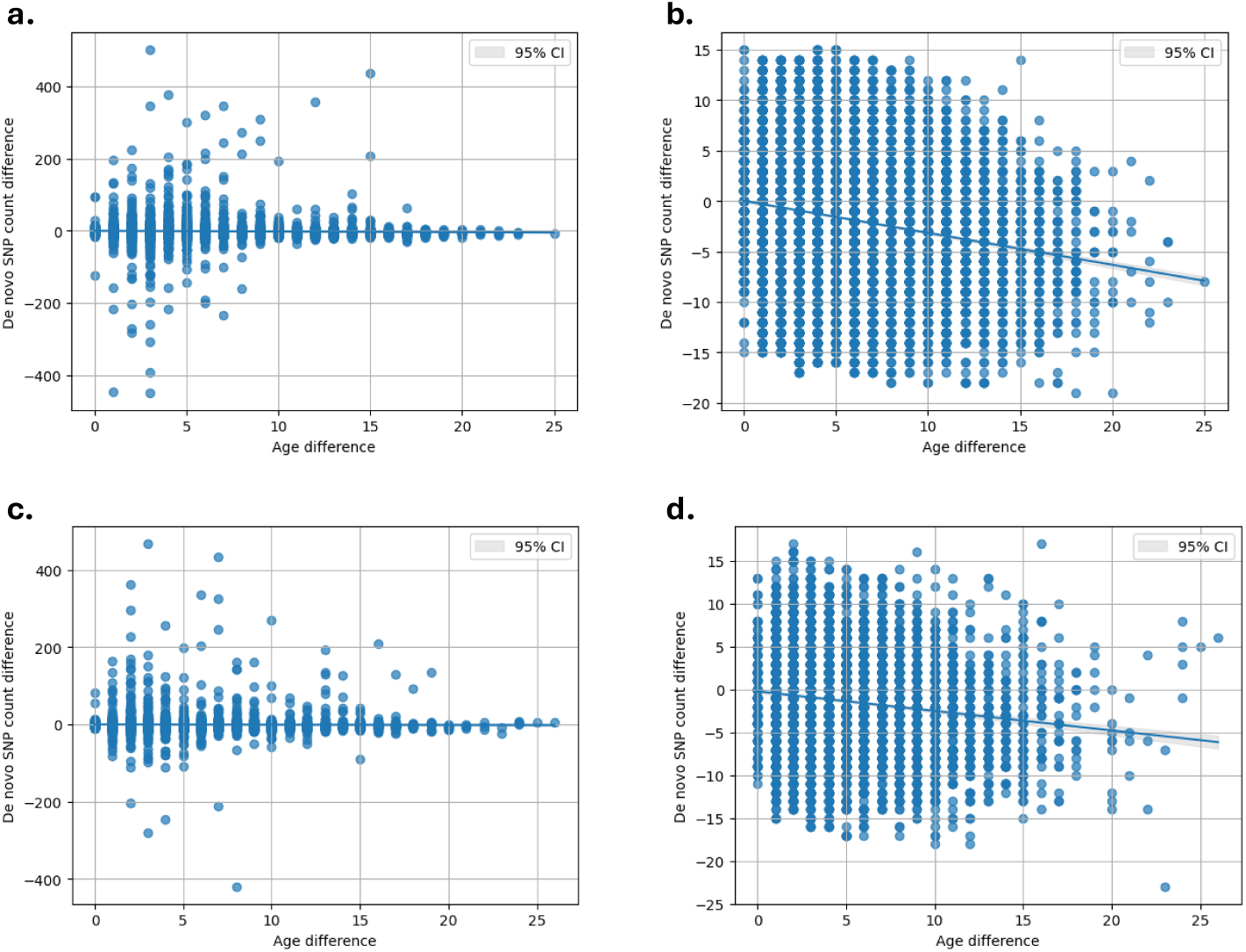
Outlier filtering based on differences in DNM counts between siblings. Difference in putative DNM counts as a function of age difference across (a) UKB sibling pairs, pre-filtering (slope = -0.1964, p*<* 10*^−^*^16^), (b) UKB sibling pairs, post-filtering (slope = -0.3165, p*<* 10*^−^*^16^), (c) AoU sibling pairs, pre-filtering (slope = -0.0678, p = 0.278), and (d) AoU sibling pairs, post-filtering (slope = -0.2273, p*<* 10*^−^*^16^). Outliers were defined as sibling pairs with differences in putative DNM counts exceeding three standard deviations from the expected mean under a Normal approximation to the Skellam distribution (Methods 8). Differences in putative DNM counts were calculated as the count in the older sibling minus the count in the younger sibling.

**Supplementary Figure 2.**
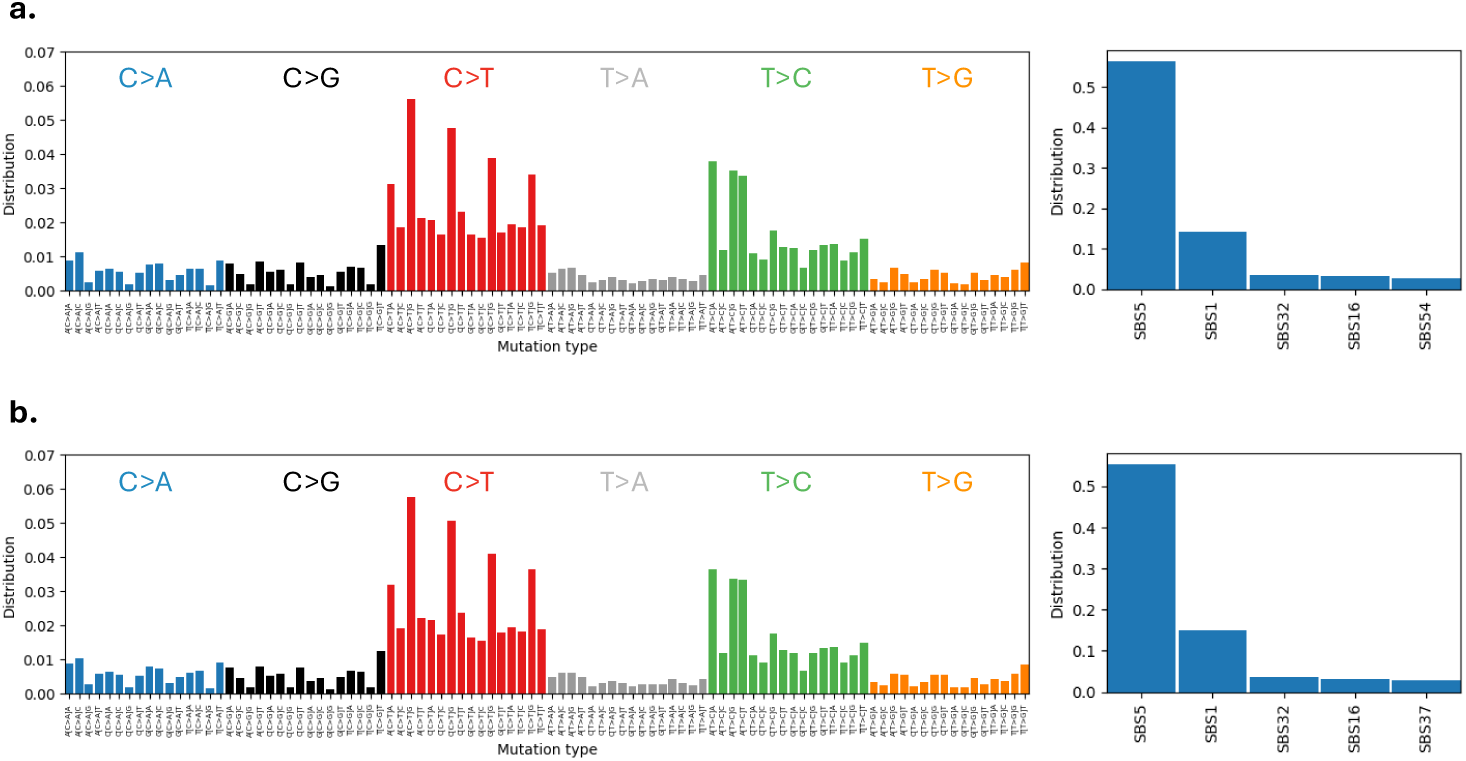
96-type mutational spectra and COSMIC SBS signature decompositions [39] for sibling-based DNMs in (a) the UKB and (b) AoU. We note that without regularization, ∼4% of DNMs are assigned to signatures associated with sequencing errors (e.g., SBS54) in both cohorts. This fraction is comparable to what is obtained using DNMs from trios, considered a gold standard: ∼4% in UKB and ∼5% in AoU (see Methods 5).

**Table S1.** Shown are estimates of the C*>*T mutation proportions for pairs of ancestry groups that differed significantly at the 5% level, after FDR correction (Methods 11). The estimated ratio of the mutation proportions in the two ancestry groups is shown, along with the standard error. The columns LCL and UCL indicate the lower and upper 95% confidence limits as calculated by the R package emmeans (Methods 11).

| Mutation | Contrast | LCL | Ratio | UCL | SE | FDR |
| --- | --- | --- | --- | --- | --- | --- |
| C>T | EUR3 / AMR1 | 0.88 | 0.92 | 0.98 | 0.025 | 0.046 |
| C>T | AMR1 / EUR1 | 1.03 | 1.07 | 1.17 | 0.035 | 0.046 |

**Supplementary Figure 3.**
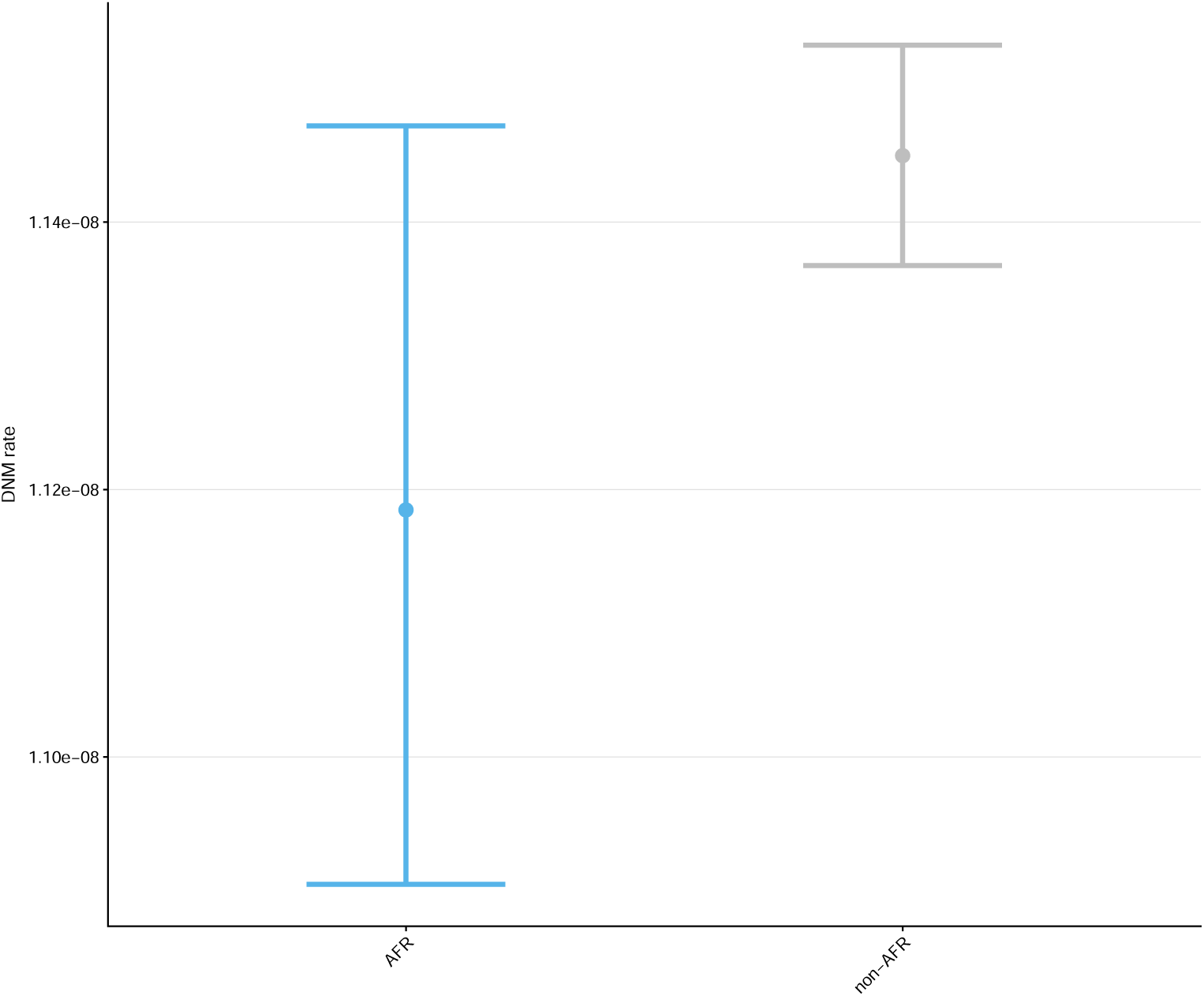
Estimates of autosomal DNM rates of individuals assigned to AFR and Non-AFR at the sample mean paternal age at conception (29.0 years), with source set to sibling and biobank set to UKB; whiskers denote 95% confidence intervals. (Methods 11)

**Supplementary Figure 4.**
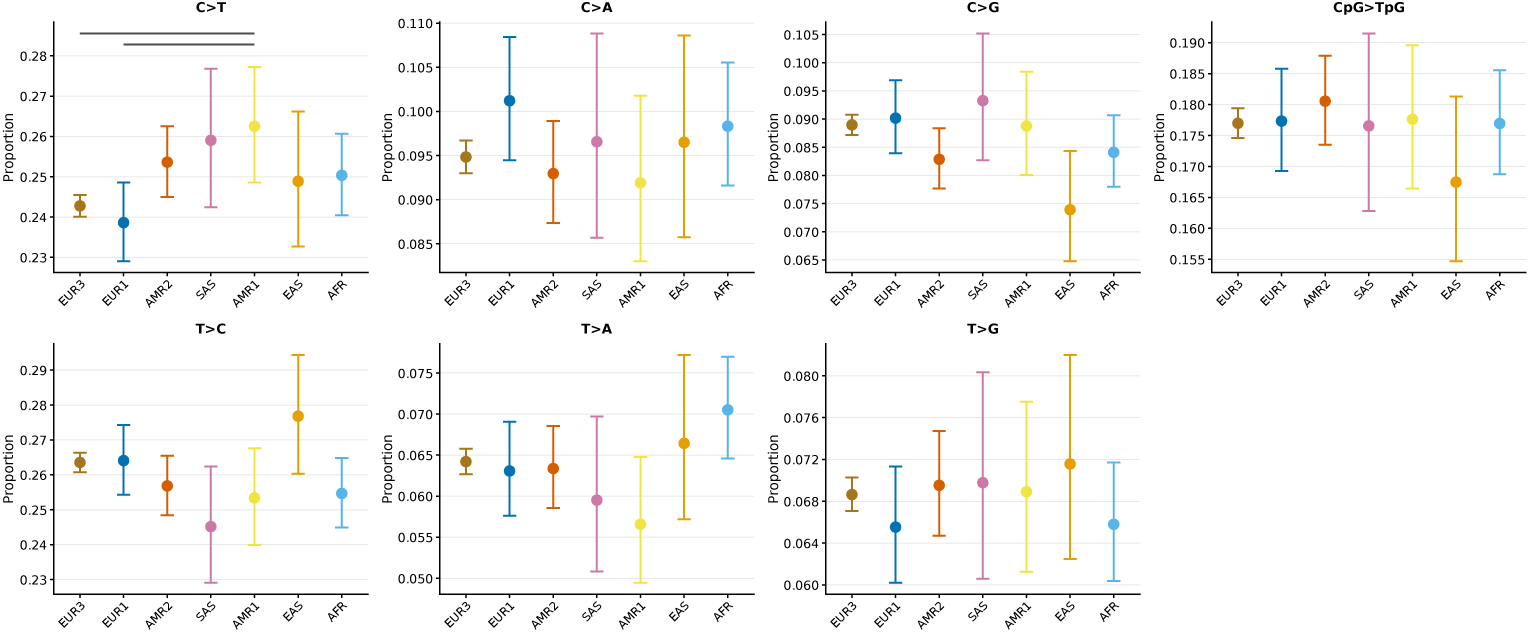
Shown are the proportion of each of 7 mutation types predicted for UKB siblings, at paternal age 29; whiskers denote 95% confidence intervals. The F-test for cluster was significant for C*>*T mutations. Brackets indicate significant pairwise comparisons after FDR correction for 7 mutation types. A previous study of DNMs did not find significant C*>*T differences among ancestry groups, possibly because of smaller number of DNMs identified in their AMR group [15] (Supplementary Table 2). In turn, they reported a lower proportion of C*>*A in the group that they label EUR than in SAS (with an estimate ratio of 0.93) as well as a lower proportion of CpG*>*TpG (for which the ratio was 0.94). Our estimates are consistent, however, as theirs fall within our (wide) 95% confidence interval for EUR3/SAS (CI: 0.92-1.09 for CpG*>*TpG and CI: 0.87-1.11 for C*>*A)

**Supplementary Table 2.** Shown are DNM counts that we used in the ancestry rate and spectra comparisons (Figure 3, Methods 11, Methods 11). For comparison, counts for [15] were estimated by multiplying average DNM counts by sample sizes provided.

| Ancestry | Dataset | DNMs |
| --- | --- | --- |
| AFR | AoU | 7,224 |
| AFR | UKB | 1,682 |
| AFR | Combined | 8,906 |
| AFR | Garcia Salinas et al. 2025 | 13,208 |
| AMR1 | AoU | 4,288 |
| AMR1 | Combined | 4,288 |
| AMR2 | AoU | 16,374 |
| AMR2 | UKB | 43 |
| AMR2 | Combined | 16,417 |
| AMR | Garcia Salinas et al. 2025 | 13,757 |
| EAS | AoU | 2,678 |
| EAS | UKB | 304 |
| EAS | Combined | 2,982 |
| EAS | Garcia Salinas et al. 2025 | 3,498 |
| EUR1 | AoU | 7,049 |
| EUR1 | UKB | 2,234 |
| EUR1 | Combined | 9,283 |
| EUR3 | AoU | 25,531 |
| EUR3 | UKB | 140,020 |
| EUR3 | Combined | 165,551 |
| EUR | Garcia Salinas et al. 2025 | 520,276 |
| MID | AoU | 812 |
| MID | UKB | 105 |
| MID | Combined | 917 |
| SAS | AoU | 990 |
| SAS | UKB | 1,611 |
| SAS | Combined | 2,601 |
| SAS | Garcia Salinas et al. 2025 | 80,825 |

**Supplementary Table 3.** Genes associated with elevated mutation rates in some somatic tissues, the corresponding COSMIC SBS signatures and the germline proportions in carriers and non-carriers of disrupting variants in those genes. Signature proportions and their differences between carriers and non-carriers (see Methods 13) were assessed using the NNLS approach described in Methods 13, under an additive model. We note that for fewer than 20 carriers, we are prohibited from providing the exact number by the AoU dissemination policy.

| Gene | Somatic signatures | Number of carriers | Number of non-carriers | Proportion in carriers | Proportion in non-carriers | p-value |
| --- | --- | --- | --- | --- | --- | --- |
| *BRCA1* | SBS3 | 112 | 27,533 | 0% | 0% | 1 |
| *BRCA2* | SBS3 | 982 | 26,663 | 0% | 0% | 1 |
| *EXO1* | SBS3, SBS6, SBS20, SBS44 | 317 | 27,328 | 0% | 0% | 1 |
| *PALB2* | SBS3 | 67 | 27,578 | 0% | 0% | 1 |
| *PAXIP1* | SBS3 | <20 | >27,625 | 19% | 0% | 0.0549 |
| *RIF1* | SBS3 | 21 | 27,624 | 1% | 0% | 0.3506 |
| *MLH1* | SBS6, SBS20, SBS44 | 599 | 27,046 | 0% | 0% | 1 |

**Supplementary Table 4.** Germline mutation rates in carriers versus non-carriers of disruptive variants in DNA repair genes with known mutator effects in the soma or germline. Significance of the elevation in mutation rate in carriers was assessed using the burden test framework described in Methods 13.

| Gene | Model | Number of carriers | Number of non-carriers | Mean rate in carriers | Mean rate in non-carriers | p-value |
| --- | --- | --- | --- | --- | --- | --- |
| *BRCA1* | Additive | 112 | 27,533 | $1.34 \times 10^{-8}$ | $1.28 \times 10^{-8}$ | 0.1283 |
| *BRCA2* | Additive | 982 | 26,663 | $1.29 \times 10^{-8}$ | $1.28 \times 10^{-8}$ | 0.2264 |
| *EXO1* | Additive | 317 | 27,328 | $1.28 \times 10^{-8}$ | $1.28 \times 10^{-8}$ | 0.4199 |
| *PALB2* | Additive | 67 | 27,578 | $1.23 \times 10^{-8}$ | $1.28 \times 10^{-8}$ | 0.7553 |
| *PAXIP1* | Additive | <20 | >27,625 | $1.40 \times 10^{-8}$ | $1.28 \times 10^{-8}$ | 0.1408 |
| *RIF1* | Additive | 21 | 27,624 | $1.25 \times 10^{-8}$ | $1.28 \times 10^{-8}$ | 0.4995 |
| *MLH1* | Additive | 599 | 27,046 | $1.25 \times 10^{-8}$ | $1.28 \times 10^{-8}$ | 0.7296 |
| *POLE* | Additive | 62 | 27,583 | $1.27 \times 10^{-8}$ | $1.28 \times 10^{-8}$ | 0.557 |
| *POLD1* | Additive | 27 | 27,618 | $1.23 \times 10^{-8}$ | $1.28 \times 10^{-8}$ | 0.6514 |
| *MUTYH* | Additive | 807 | 26,838 | $1.27 \times 10^{-8}$ | $1.28 \times 10^{-8}$ | 0.5379 |
| *MPG* | Additive | 88 | 27,557 | $1.26 \times 10^{-8}$ | $1.28 \times 10^{-8}$ | 0.6883 |
| *XPC* | Additive | 64 | 27,581 | $1.32 \times 10^{-8}$ | $1.28 \times 10^{-8}$ | 0.1537 |
| *MUTYH* | Recessive | <20 | >27,625 | $1.28 \times 10^{-8}$ | $1.28 \times 10^{-8}$ | 0.29 |

**Supplementary Figure 5.**
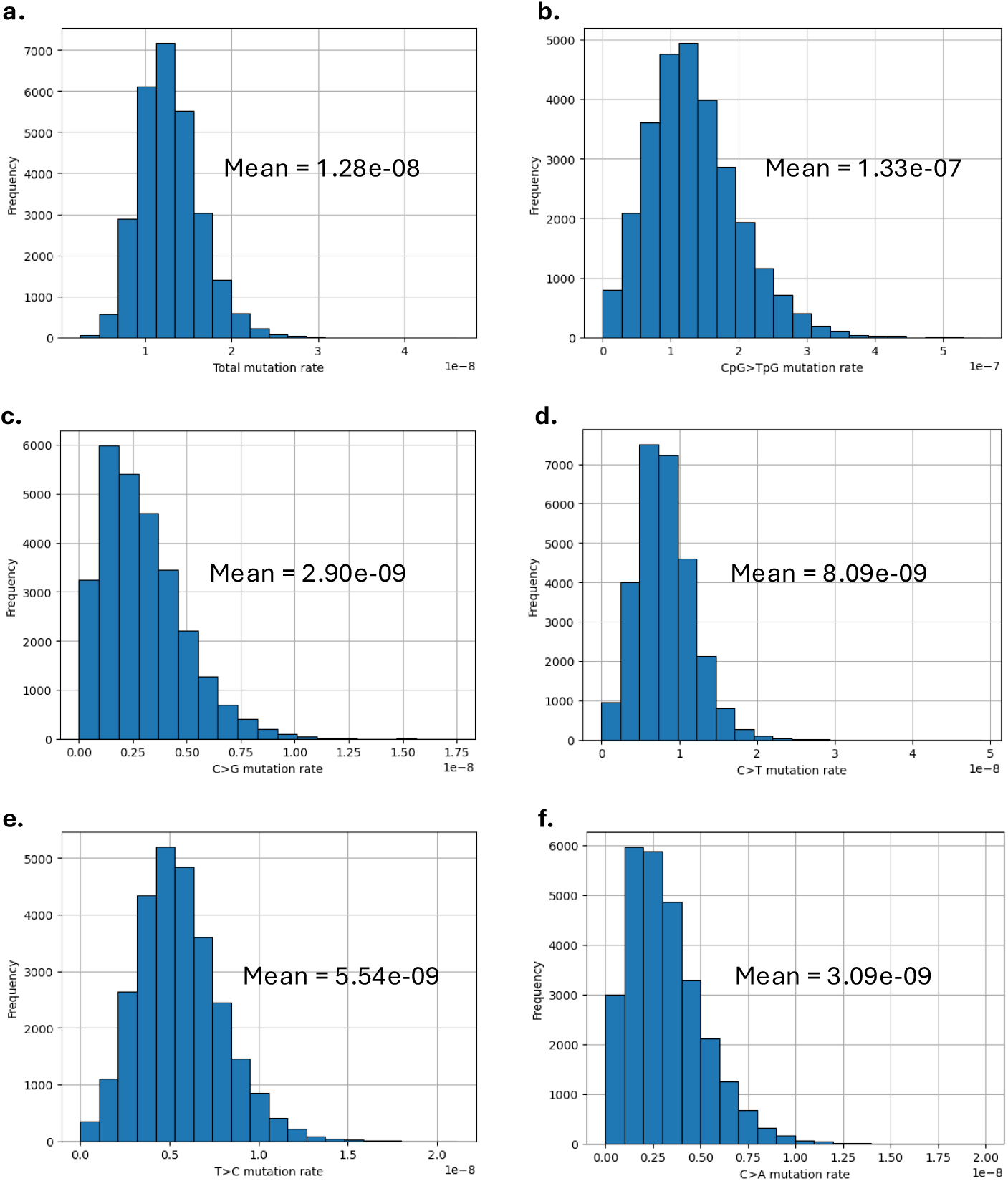
Distribution of raw mutation rate phenotypes used in burden tests. Histograms show the distribution of (a) the total autosomal DNM rate, (b) the autosomal rate of CpG*>*TpG, (c) of C*>*G, (d) of C*>*T outside a CpG context, (e) of T*>*C, and (f) of C*>*A across families in the UKB and AoU, calculated as described in Methods 13. Mean values are indicated.

**Supplementary Table 5.** Association of loss-of-function and deleterious missense variants in DNA repair genes with total DNM rates and mutation rates for specific subtypes, based on the approach described in Methods 13. Shown are genes for which q*<* 0.2, after applying a Benjamini-Hochberg FDR across 138 genes for each phenotype separately.

| Gene | Model | Mutation rate phenotype | Number of carriers | Number of non-carriers | Mean rate in carriers | Mean rate in non-carriers | p-value | q-value |
| --- | --- | --- | --- | --- | --- | --- | --- | --- |
| *REV1* | Additive | Total | 26 | 27,619 | $1.48 \times 10^{-8}$ | $1.28 \times 10^{-8}$ | 0.0006 | 0.0806 |
| *LIG1* | Additive | CpG>TpG | 85 | 27,560 | $1.65 \times 10^{-7}$ | $1.33 \times 10^{-7}$ | 0.00003 | 0.0045 |
| *RAD51C* | Additive | CpG>TpG | 24 | 27,621 | $1.72 \times 10^{-7}$ | $1.33 \times 10^{-7}$ | 0.0022 | 0.1507 |
| *SLX4* | Additive | C>T | 64 | 27,581 | $9.4 \times 10^{-9}$ | $8.09 \times 10^{-9}$ | 0.0011 | 0.1536 |
| *DNA2* | Recessive | T>C | 22 | 27,623 | $6.66 \times 10^{-9}$ | $5.54 \times 10^{-9}$ | 0.0166 | 0.1828 |

**Supplementary Table 6.** Comparison of the effect of LoF versus deleterious missense variants on mutation rate phenotypes for the genes reported in Table 5. The F-test p-value, computed as described in Methods 13, tests whether LoF and deleterious missense variants have significantly different effects on the mutation rate phenotype. For *DNA2*, when LoF and missense variants are modeled separately, families carrying ∼1 LoF and ∼1 missense variant become non-carriers under the recessive model, reducing the total carrier count from 22 to 19. A LoF p-value cannot be computed in the absence of LoF carriers; results for *DNA2* are therefore excluded from this analysis.

| Gene | Model | Mutation rate phenotype | Number of LoF carriers | Number of missense carriers | LoF p-value | Missense p-value | F-test p-value |
| --- | --- | --- | --- | --- | --- | --- | --- |
| *REV1* | Additive | Total | 5 | 21 | $1.55 \times 10^{-5}$ | 0.0108 | 0.7457 |
| *LIG1* | Additive | CpG>TpG | 38 | 47 | 0.0344 | 0.000496 | 0.6077 |
| *RAD51C* | Additive | CpG>TpG | 21 | 3 | 0.0385 | $3.86 \times 10^{-12}$ | $2.49 \times 10^{-5}$ |
| *SLX4* | Additive | C>T | 63 | 1 | 0.00241 | 0.9995 | 0.9995 |
| *DNA2* | Recessive | T>C | 0 | 19 | — | — | — |

**Supplementary Table 7.** COSMIC SBS signature decompositions for carriers versus non-carriers of disruptive variants in *REV1* under an additive model (see Methods 13). Signature proportions were estimated using the NNLS framework described in Methods 13, normalized either by the total contribution of all signatures estimated by NNLS (columns 2–3) or by the total mutation count (columns 4–5). Two-tailed p-values are reported for the former.

| Signature | Fraction in carriers | Fraction in non-carriers | Two-tailed p-value | Fraction in carriers (total mutation count) | Fraction in non-carriers (total mutation count) |
| --- | --- | --- | --- | --- | --- |
| SBS5 | 0.5826 | 0.8312 | 0.0669 | 0.5766 | 0.8370 |
| SBS1 | 0.1442 | 0.1445 | 0.9840 | 0.1427 | 0.1455 |
| SBS39 | 0.0416 | 0.0000 | 0.1499 | 0.0412 | 0.0000 |
| SBS16 | 0.0174 | 0.0000 | 0.3427 | 0.0172 | 0.0000 |
| SBS3 | 0.0920 | 0.0000 | 0.0340 | 0.0911 | 0.0000 |
| SBS6 | 0.0191 | 0.0000 | 0.2478 | 0.0189 | 0.0000 |
| SBS15 | 0.0175 | 0.0000 | 0.0609 | 0.0173 | 0.0000 |
| SBS21 | 0.0003 | 0.0000 | 0.3247 | 0.0003 | 0.0000 |
| SBS26 | 0.0167 | 0.0021 | 0.2957 | 0.0165 | 0.0021 |
| SBS10b | 0.0128 | 0.0079 | 0.7203 | 0.0127 | 0.0080 |
| SBS30 | 0.0558 | 0.0142 | 0.1499 | 0.0553 | 0.0143 |

**Supplementary Figure 6.**
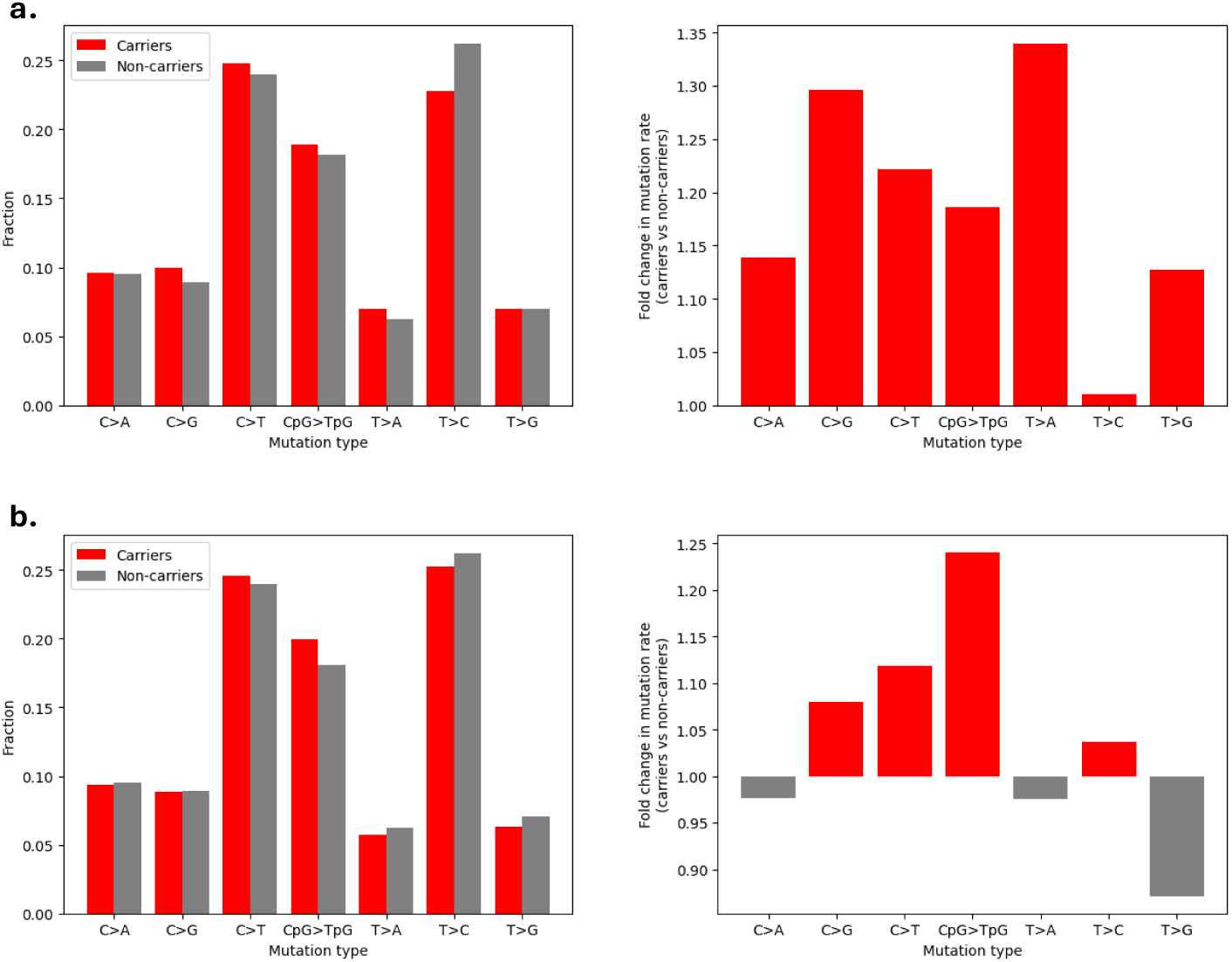
Mutational spectra and fold change in mutation rates in carriers vs non-carriers of LoF and missense variants for (a) *REV1* and (b) *LIG1*, estimated under an additive model (see Methods 13).

**Supplementary Table 8.** COSMIC SBS signature decompositions for carriers versus non-carriers of disruptive variants in *LIG1* under an additive model (see Methods 13). Signature proportions were estimated using the NNLS framework described in Methods 13, normalized either by the total contribution of all signatures estimated by NNLS (columns 2–3) or by the total mutation count (columns 4–5). Two-tailed p-values are reported for the former.

| Signature | Fraction in carriers | Fraction in non-carriers | Two-tailed p-value | Fraction in carriers (total mutation count) | Fraction in non-carriers (total mutation count) |
| --- | --- | --- | --- | --- | --- |
| SBS5 | 0.7183 | 0.8312 | 0.0669 | 0.7250 | 0.8370 |
| SBS1 | 0.1530 | 0.1445 | 0.4436 | 0.1544 | 0.1455 |
| SBS39 | 0.0180 | 0.0000 | 0.2368 | 0.0181 | 0.0000 |
| SBS16 | 0.0286 | 0.0000 | 0.0500 | 0.0289 | 0.0000 |
| SBS6 | 0.0237 | 0.0000 | 0.0599 | 0.0240 | 0.0000 |
| SBS21 | 0.0107 | 0.0000 | 0.0420 | 0.0108 | 0.0000 |
| SBS26 | 0.0000 | 0.0021 | 0.5914 | 0.0000 | 0.0021 |
| SBS10b | 0.0164 | 0.0079 | 0.1858 | 0.0165 | 0.0080 |
| SBS30 | 0.0313 | 0.0143 | 0.2168 | 0.0316 | 0.0144 |

**Supplementary Figure 7.**
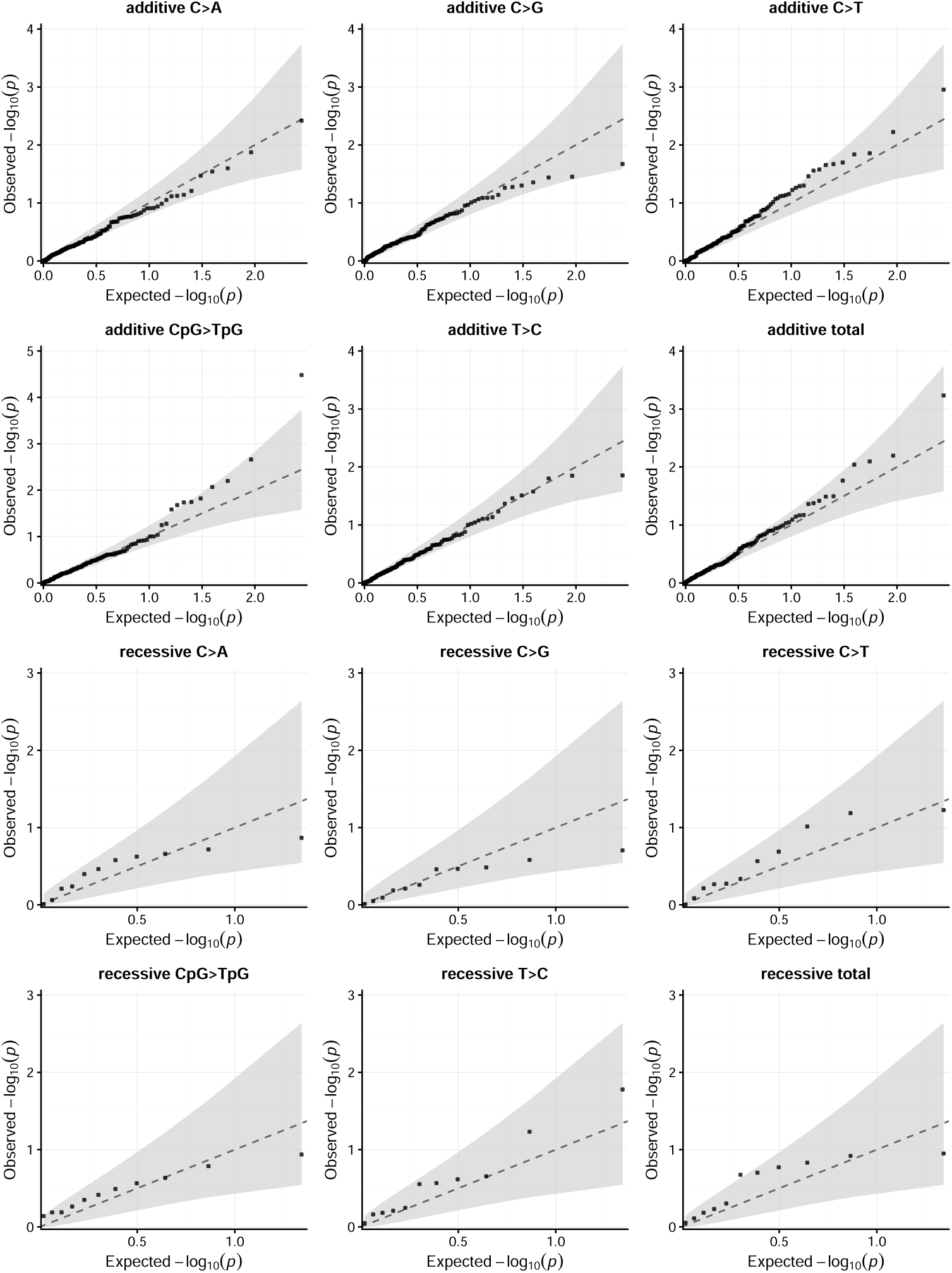
Q-Q plots for DNA repair genes with at least 20 carrier families in the UKB and AoU combined. The gray band represents the 95% CI under the null (see Methods 13)

**Supplementary Figure 8.**
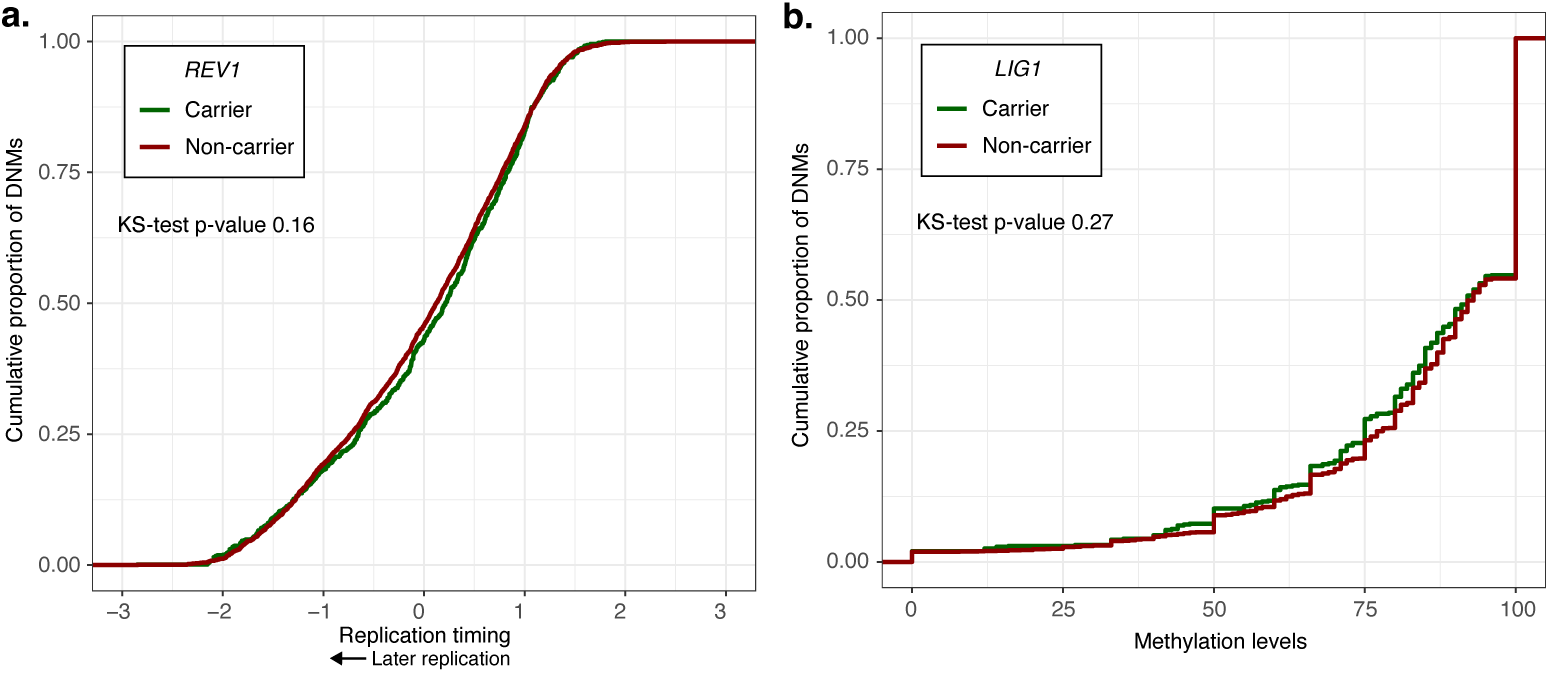
(a) Cumulative proportion of DNMs in carriers and non-carriers of LoF and deleterious missense mutations in *REV1* as a function of replication timing. Replication timing was measured in lymphoblastoid cell lines; the mean is set to 0 and later replication timing corresponds to lower values (see Methods 10). (b) Cumulative proportion of CpG*>*TpG mutations in carriers and non-carriers of LoF and deleterious missense mutations in *LIG1* as a function of methylation levels in bulk testis samples (see Methods 10). The p-values for the Kolmogorov-Smirnov test are two-tailed.

**Supplementary Table 9.**
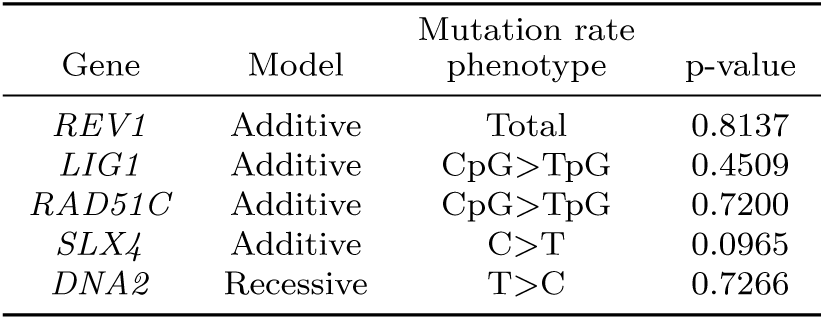
Test of genotype-by-biobank interaction for the genes reported in Table 13. The p-value, computed as described in Methods 13, tests whether the effect of LoF and deleterious missense variants on mutation rate phenotypes differs significantly across UKB and AoU.

| Gene | Model | Mutation rate phenotype | p-value |
| --- | --- | --- | --- |
| *REV1* | Additive | Total | 0.8137 |
| *LIG1* | Additive | CpG>TpG | 0.4509 |
| *RAD51C* | Additive | CpG>TpG | 0.7200 |
| *SLX4* | Additive | C>T | 0.0965 |
| *DNA2* | Recessive | T>C | 0.7266 |

**Supplementary Figure 9.**
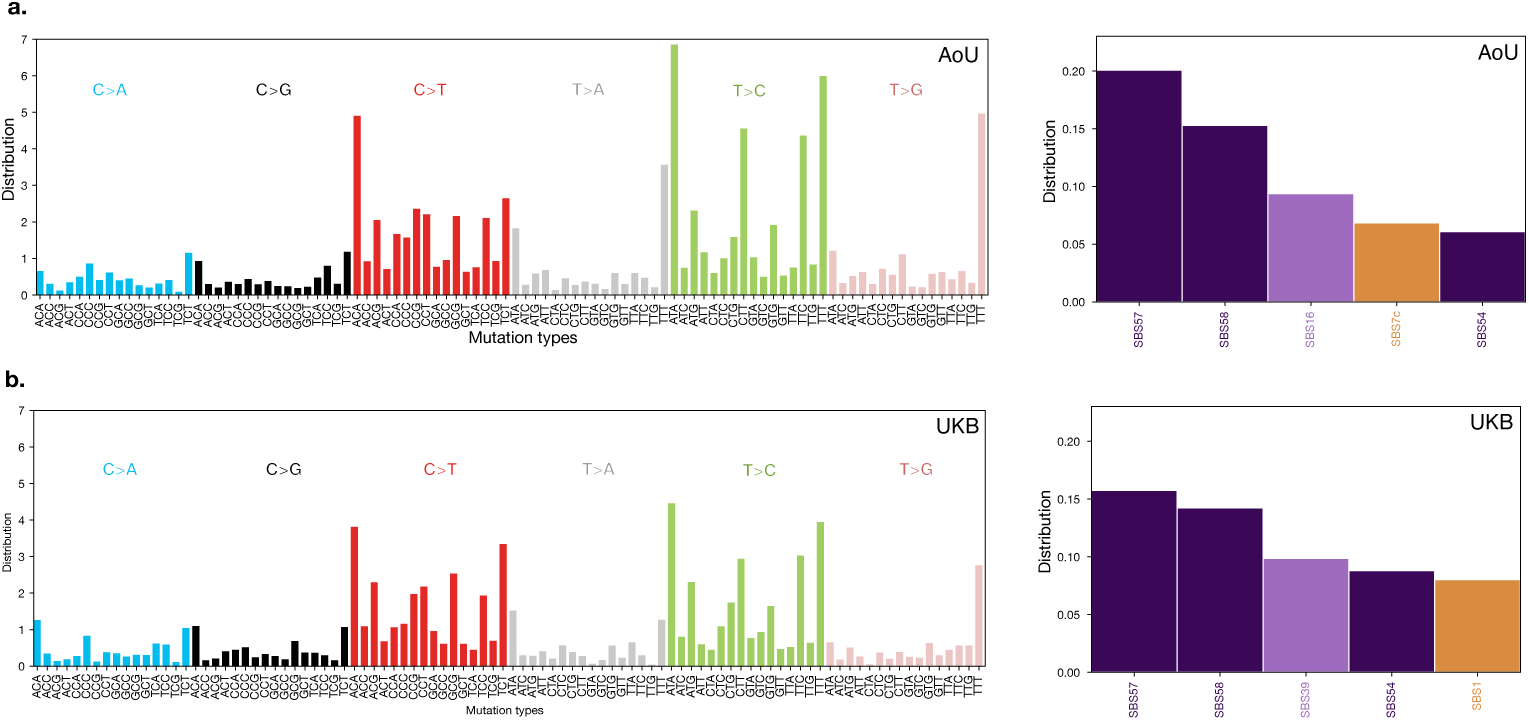
96-type mutational spectra and COSMIC SBS signature decompositions [39] for differences between monozygotic twins or genomic duplicates in (a) AoU and (b) the UKB before removing GIAB difficult regions. COSMIC SBS signatures that are possible sequencing artifacts are in dark purple, those that are annotated as unknown are in light purple, and signatures with known etiologies are in orange.

**Supplementary Figure 10.**
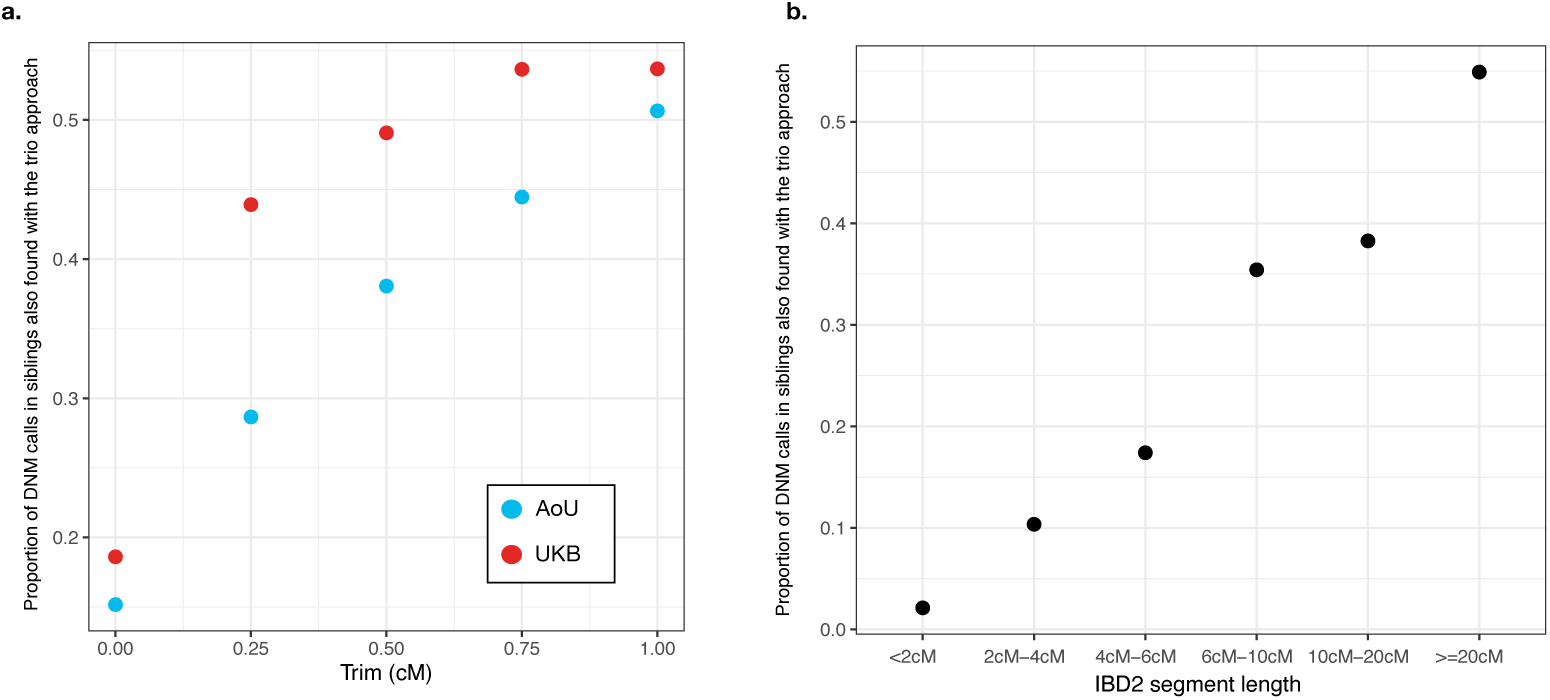
The overlap of SNP DNM calls made using the sibling and the trio approaches in 35 quads in the UKB and 192 quads after excluding GIAB difficult regions across (a) IBD2 trims and (b) IBD2 segment lengths (Methods 7). We note that the overlaps reported are before application of the allele frequency and the distance filters.

**Supplementary Table 10.**
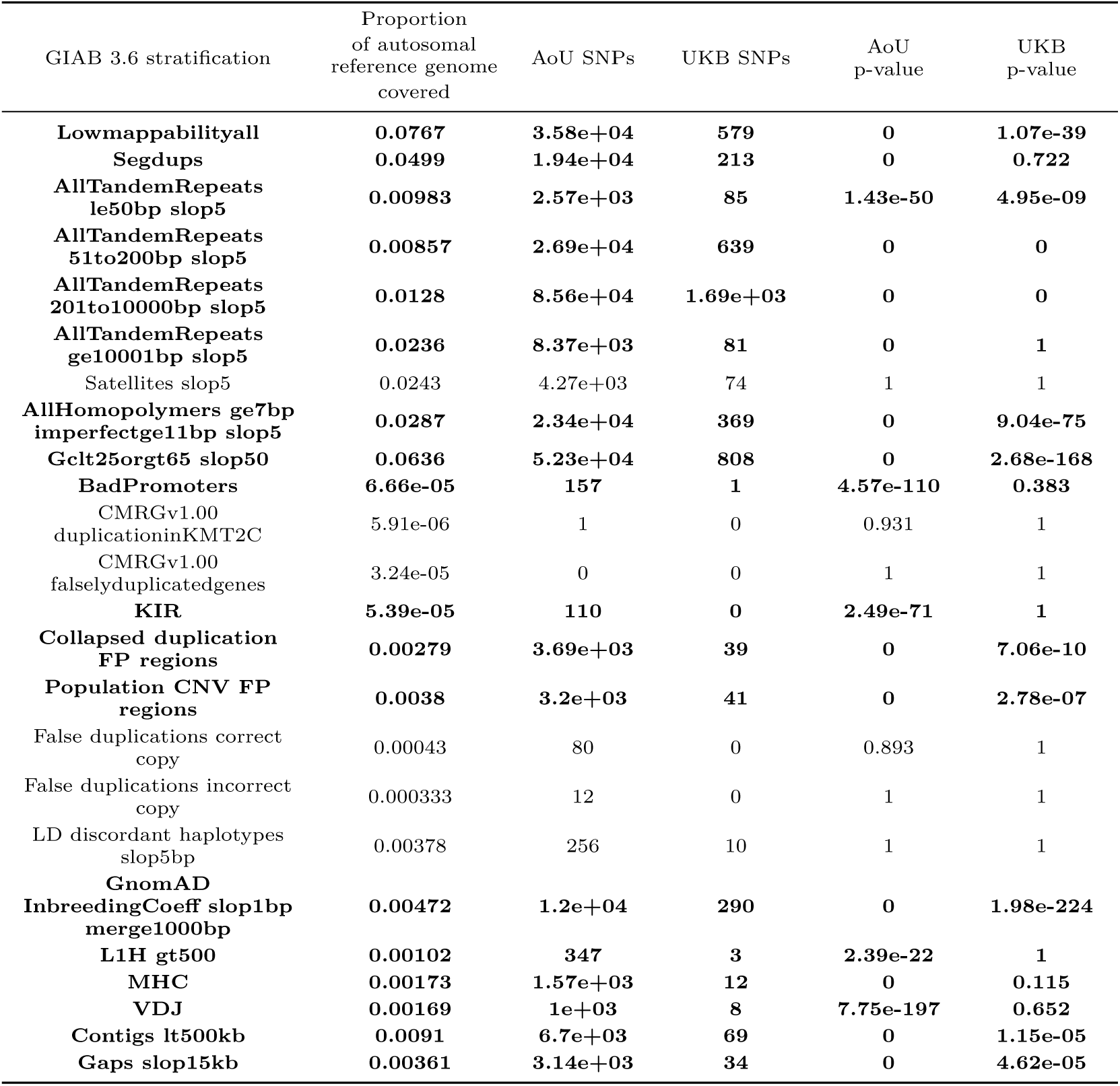
The proportion of the autosomal reference genome in each “difficult” GIAB 3.6 stratification, the number of differences between monozygotic twins or genomic duplicates in the UKB and AoU that fell within each stratification, and the FDR corrected p-values for the null hypothesis that the number of differences that fall within each stratification is proportional to the amount of the autosomal reference genome covered by that stratification. Stratifications for which the FDR corrected p-value was ≤ 0.05 for either the AoU or the UKB differences are in bold; these regions of the genome were excluded from further analyses.

**Supplementary Figure 11.**
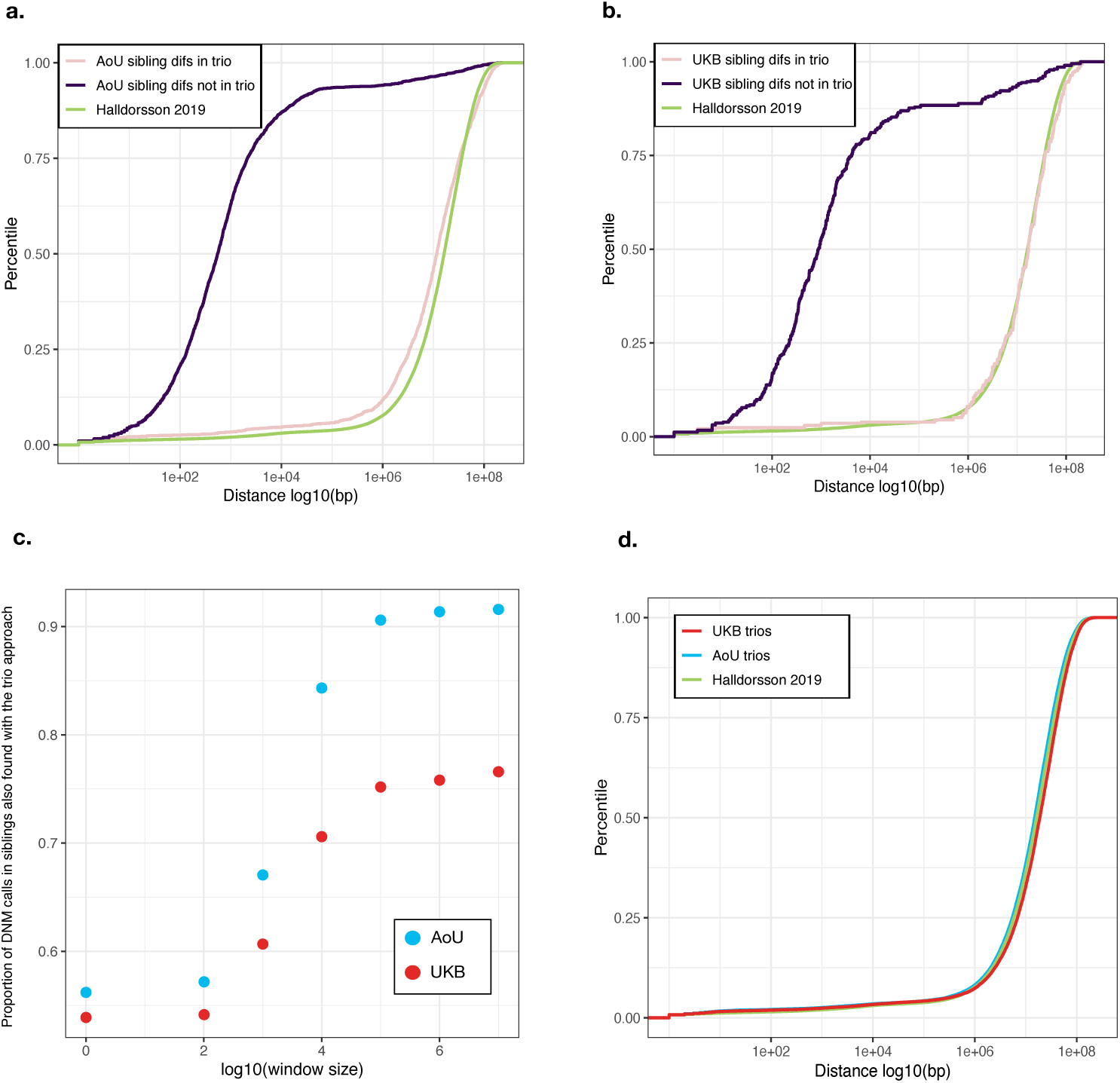
Evaluation of distance-based filters in UKB and AoU quads based on the overlap of DNM calls based on sibling versus trios (a-b) The empirical CDF of the distance to the nearest DNM of DNMs called only using siblings (purple), DNMs identified using both the sibling and the trio approaches (pink) and DNMs identified based on nuclear families in [32], for (a) the AoU quads and (b) the UKB quads. This comparison is made in IBD2 segments after trimming ends, removing small segments, and filtering for error-prone GIAB regions (Methods 7). (c) The proportion of DNMs identified in siblings that were also found using the trio approach, after applying varying window sizes to filter out clustered mutations. (d) The empirical CDF of the distances to the nearest DNM of DNMs identified across all trios in the UKB (red), AoU (blue) and [32] (green), after removing error-prone GIAB regions.

**Supplementary Figure 12.**
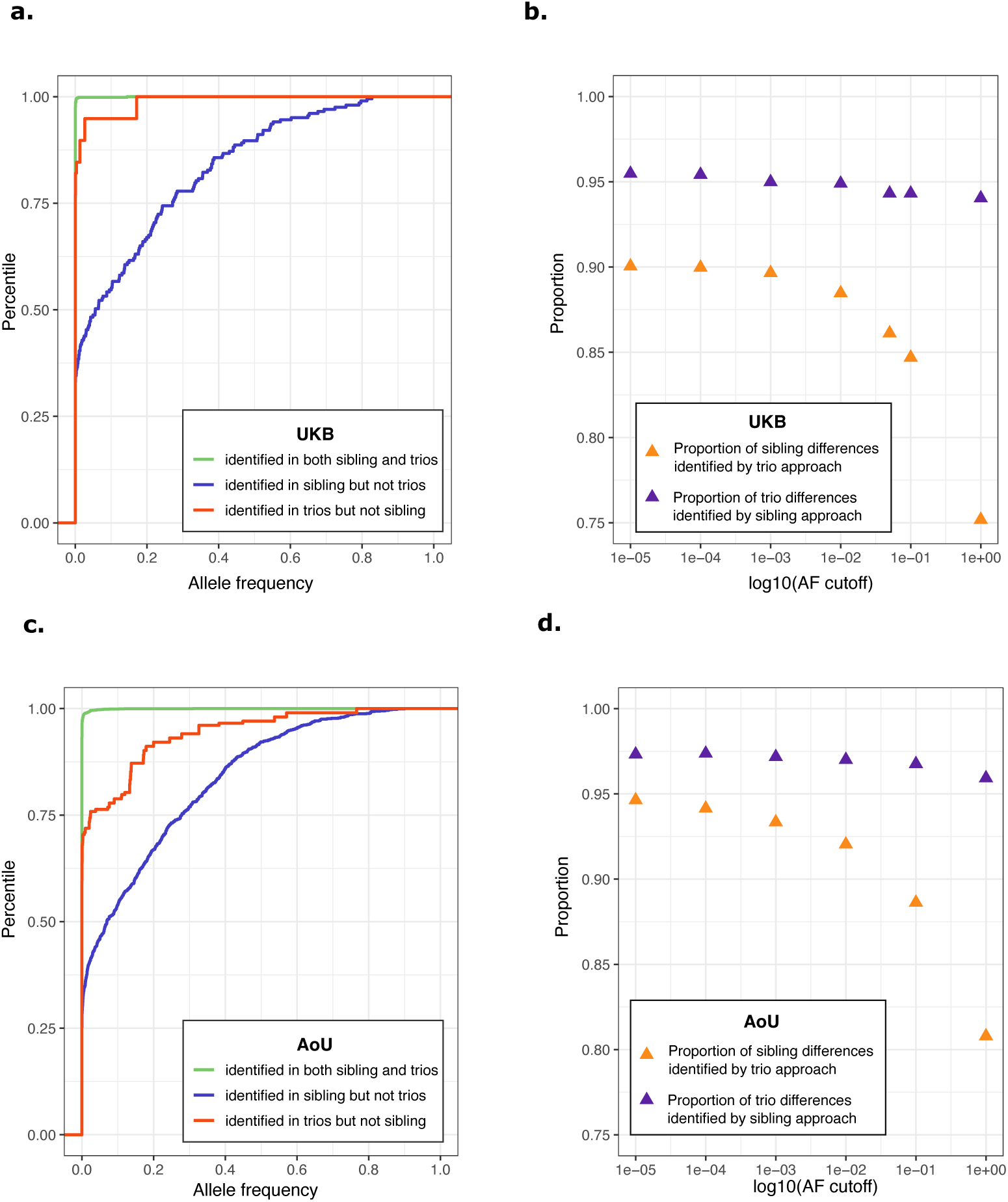
(a,c) The empirical CDF of the allele frequencies of mutations identified from both the sibling and the trio approaches (green), mutations identified using siblings but not from the trio approach (blue), and mutations that were identified by the trio approach but not in siblings. (b,d) The proportion of sibling differences that were identified by the trio approach and *vice versa*, for different allele frequency cutoffs.

**Supplementary Figure 13.**
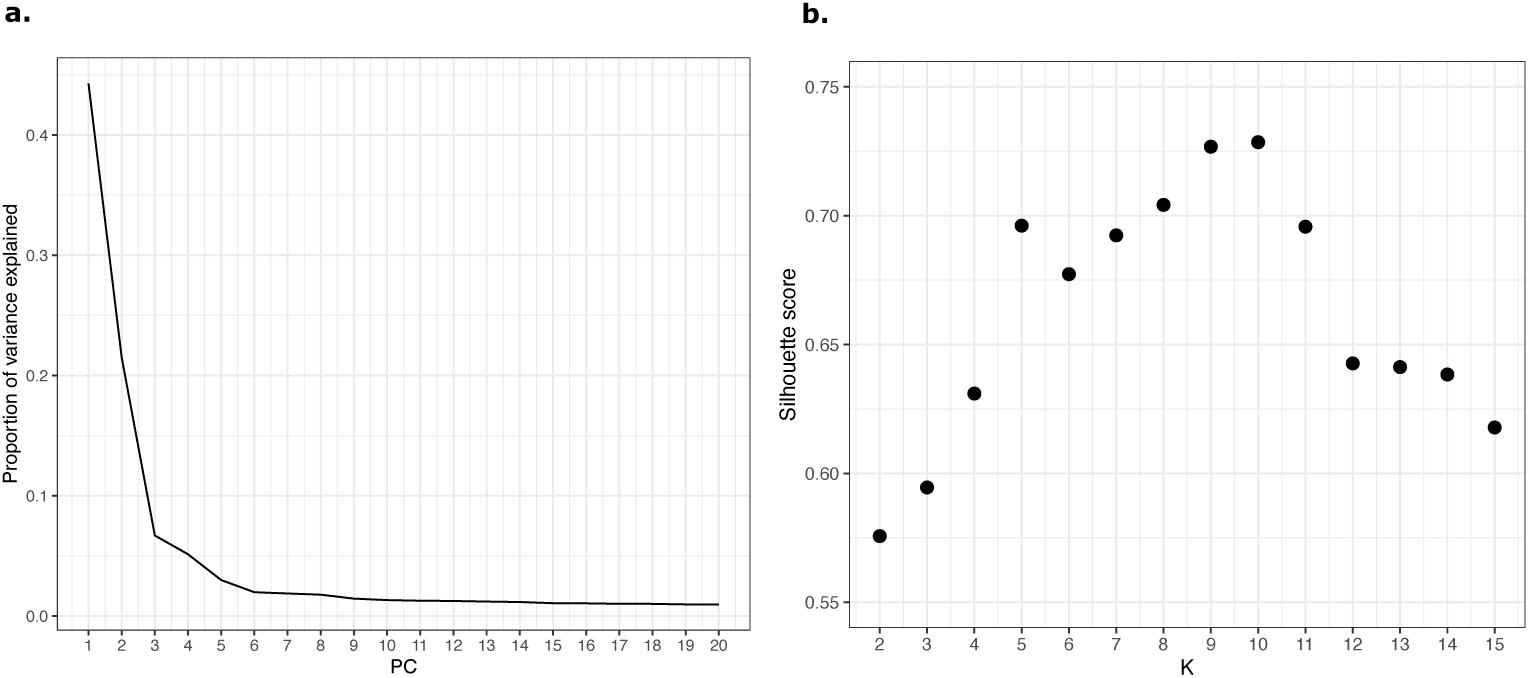
Genetic ancestry PCs obtained for HGDP-1KG samples (Methods 11). (a) The proportion of variance explained by each PC, out of the total variance explained by the first 20 PCs. (b) Silhouette scores for *k* = 1 through 15.

**Supplementary Figure 14.**
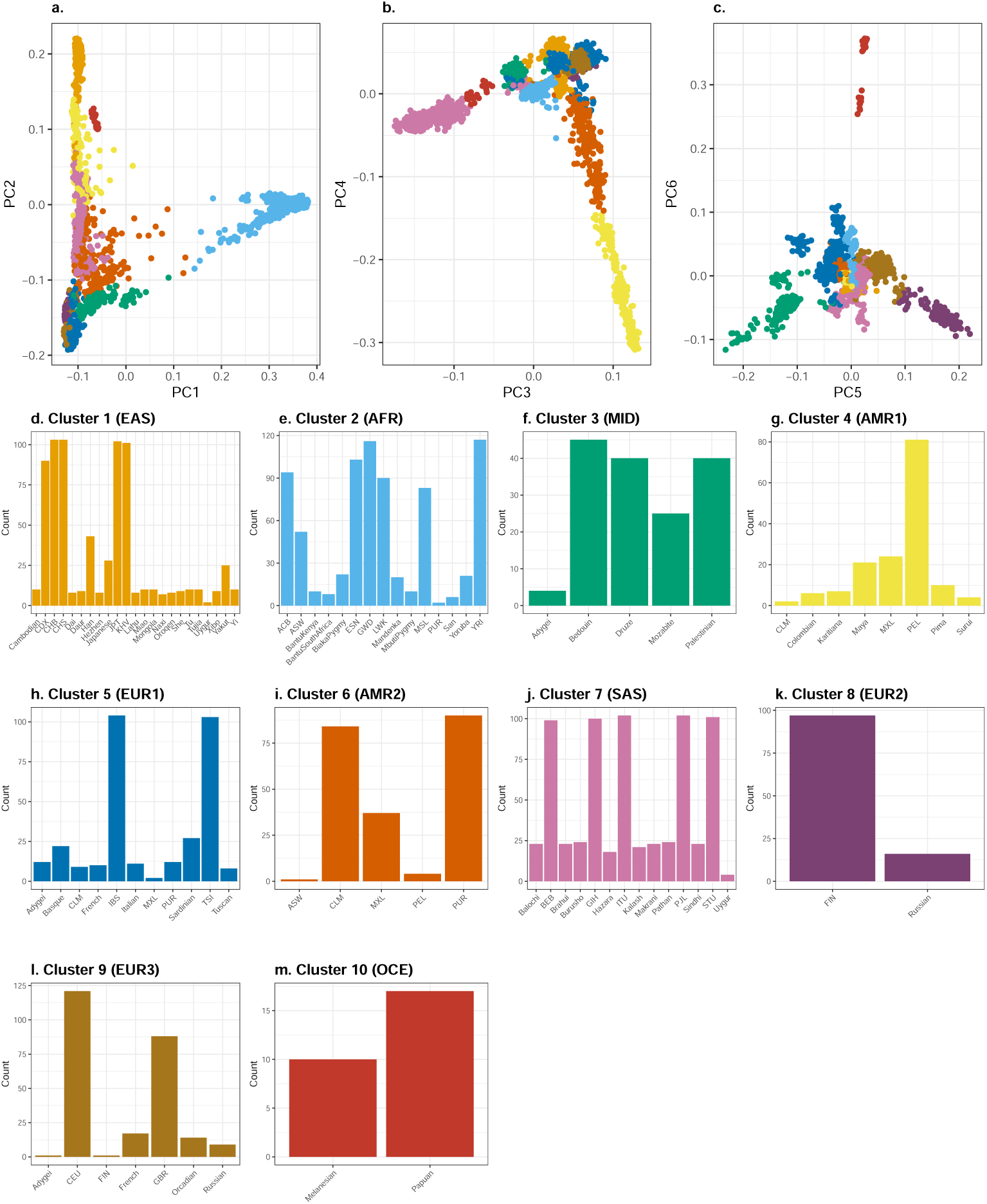
Genetic ancestry clusters with *k* = 10 (Methods 11). (a-c) Each point represents a sample from HGDP-1KG. The color of each point corresponds to a cluster (referred to as a genetic ancestry group in the text). (d-m) The number of people with a given HGDP and 1KG population label within each of our ancestry clusters.

**Supplementary Figure 15.**
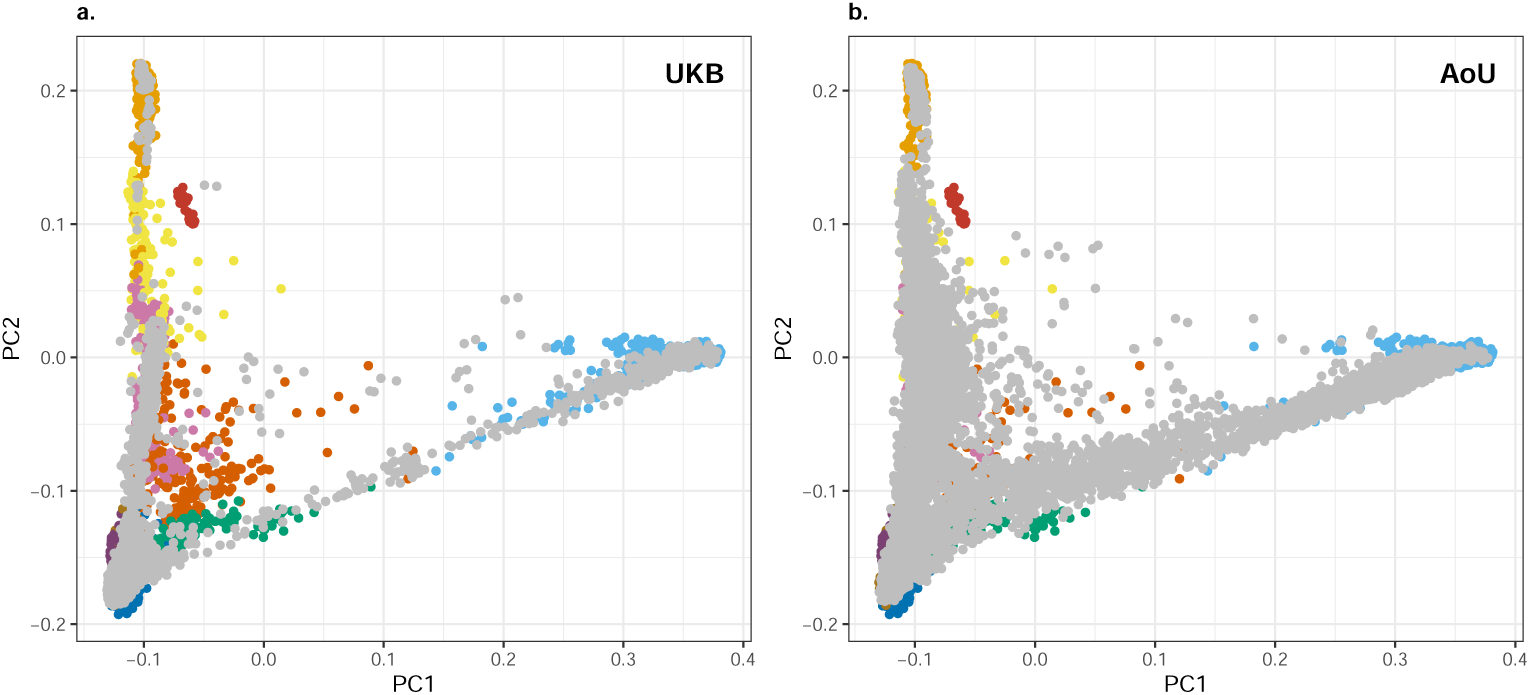
Genetic ancestries of siblings and trios in (a) UKB and (b) AoU. Colors denote the 10 ancestry clusters that we identified using the HGDP-1KG samples. Each gray point represents a person in a sibling pair or trio.

**Supplementary Figure 16.**
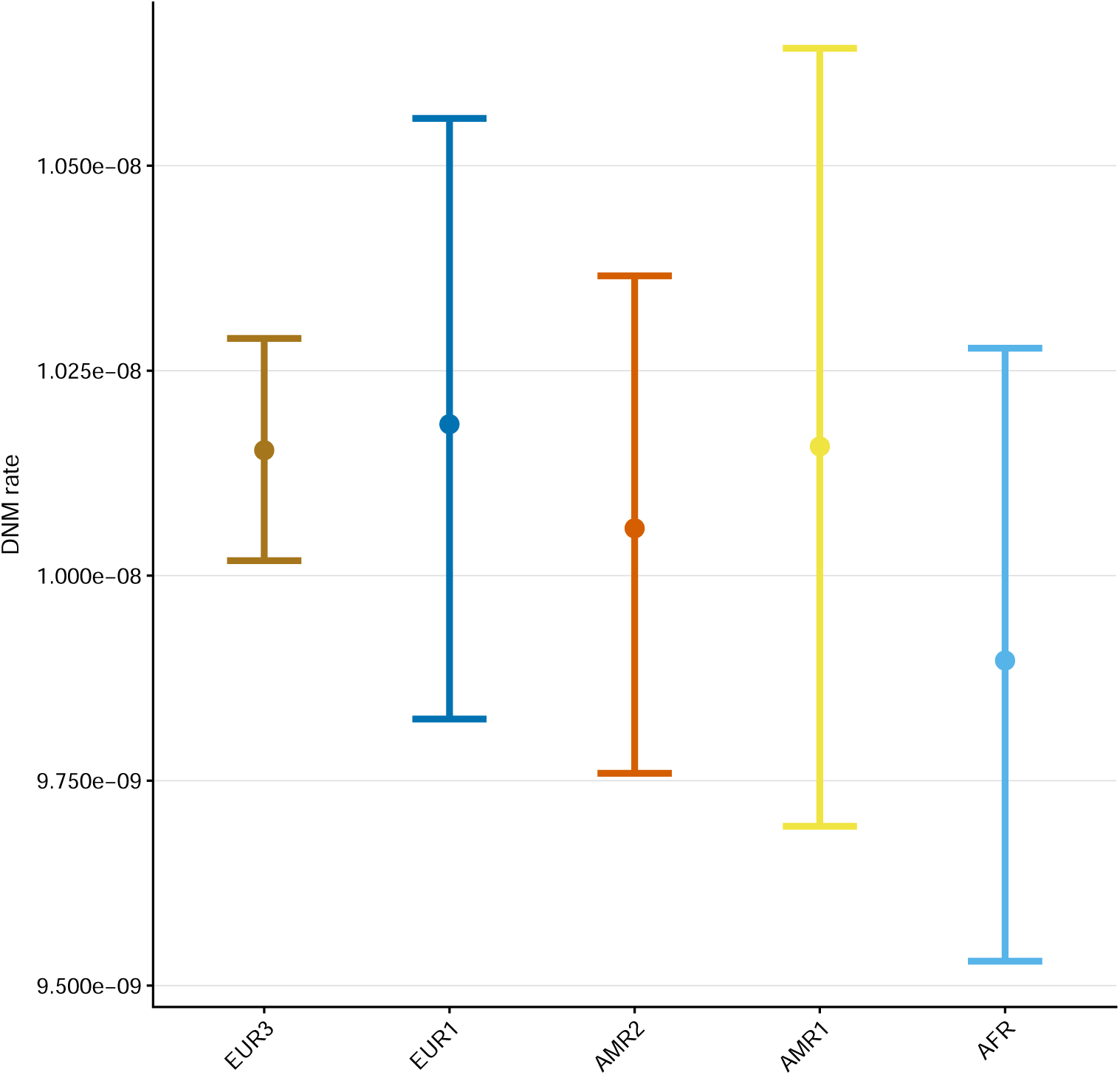
Estimates of autosomal DNM rates for each ancestry cluster at the sample mean paternal age at conception (27.2 years of age) and maternal age (25.0), with the biobank set to UKB. Shown are mean rates for the autosomal genome, with whiskers denoting 95% confidence intervals (Methods 11).

**Supplementary Table 11.**
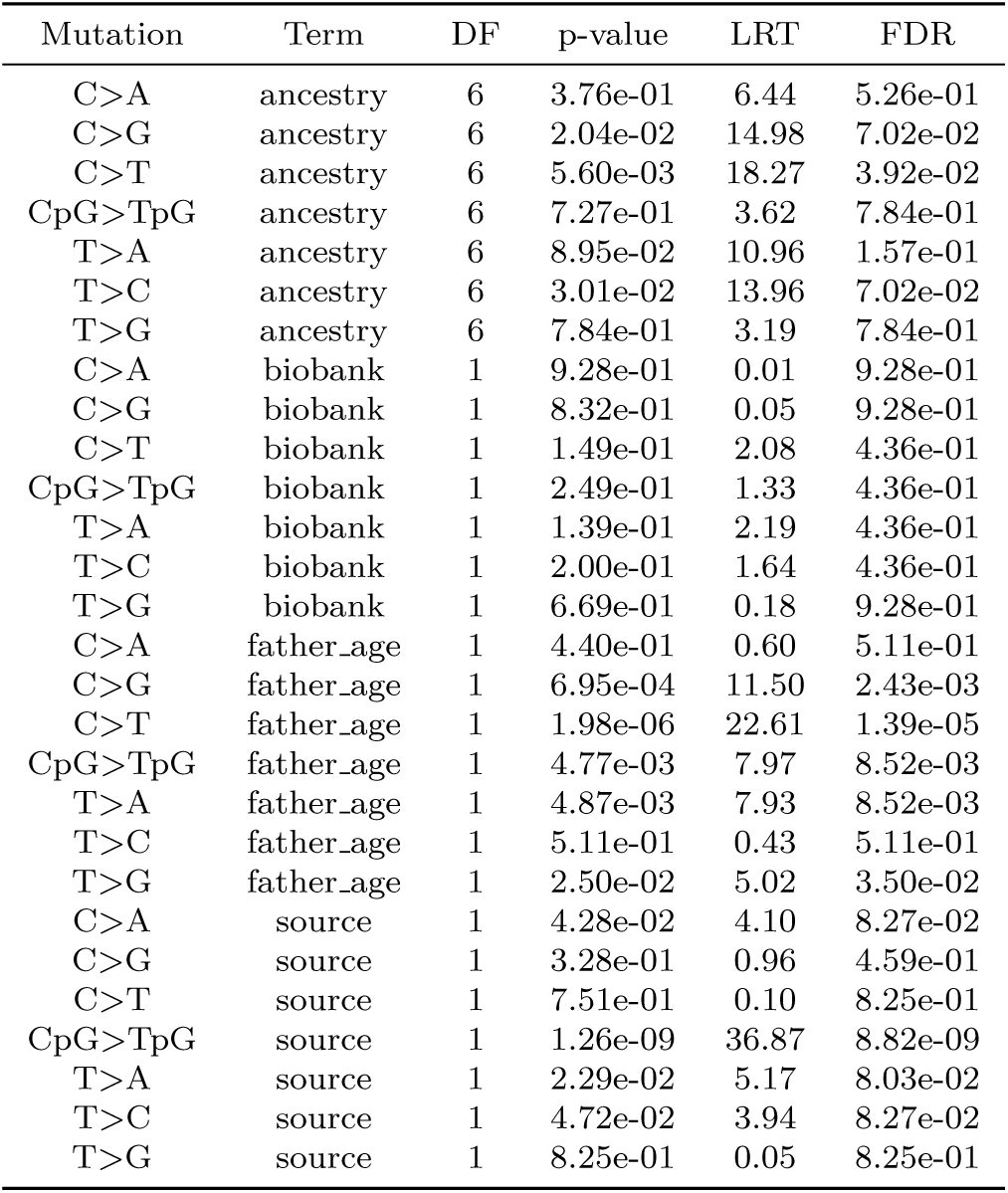
Assessing the significance of predictors for each mutation type. We used single-term deletion likelihood-ratio tests to assess whether each predictor improved the fit using the drop1() function in R. We applied an FDR correction for each predictor separately (seven tests per predictor, Methods 11).

| Mutation | Term | DF | p-value | LRT | FDR |
| --- | --- | --- | --- | --- | --- |
| C>A | ancestry | 6 | 3.76e-01 | 6.44 | 5.26e-01 |
| C>G | ancestry | 6 | 2.04e-02 | 14.98 | 7.02e-02 |
| C>T | ancestry | 6 | 5.60e-03 | 18.27 | 3.92e-02 |
| CpG>TpG | ancestry | 6 | 7.27e-01 | 3.62 | 7.84e-01 |
| T>A | ancestry | 6 | 8.95e-02 | 10.96 | 1.57e-01 |
| T>C | ancestry | 6 | 3.01e-02 | 13.96 | 7.02e-02 |
| T>G | ancestry | 6 | 7.84e-01 | 3.19 | 7.84e-01 |
| C>A | biobank | 1 | 9.28e-01 | 0.01 | 9.28e-01 |
| C>G | biobank | 1 | 8.32e-01 | 0.05 | 9.28e-01 |
| C>T | biobank | 1 | 1.49e-01 | 2.08 | 4.36e-01 |
| CpG>TpG | biobank | 1 | 2.49e-01 | 1.33 | 4.36e-01 |
| T>A | biobank | 1 | 1.39e-01 | 2.19 | 4.36e-01 |
| T>C | biobank | 1 | 2.00e-01 | 1.64 | 4.36e-01 |
| T>G | biobank | 1 | 6.69e-01 | 0.18 | 9.28e-01 |
| C>A | father_age | 1 | 4.40e-01 | 0.60 | 5.11e-01 |
| C>G | father_age | 1 | 6.95e-04 | 11.50 | 2.43e-03 |
| C>T | father_age | 1 | 1.98e-06 | 22.61 | 1.39e-05 |
| CpG>TpG | father_age | 1 | 4.77e-03 | 7.97 | 8.52e-03 |
| T>A | father_age | 1 | 4.87e-03 | 7.93 | 8.52e-03 |
| T>C | father_age | 1 | 5.11e-01 | 0.43 | 5.11e-01 |
| T>G | father_age | 1 | 2.50e-02 | 5.02 | 3.50e-02 |
| C>A | source | 1 | 4.28e-02 | 4.10 | 8.27e-02 |
| C>G | source | 1 | 3.28e-01 | 0.96 | 4.59e-01 |
| C>T | source | 1 | 7.51e-01 | 0.10 | 8.25e-01 |
| CpG>TpG | source | 1 | 1.26e-09 | 36.87 | 8.82e-09 |
| T>A | source | 1 | 2.29e-02 | 5.17 | 8.03e-02 |
| T>C | source | 1 | 4.72e-02 | 3.94 | 8.27e-02 |
| T>G | source | 1 | 8.25e-01 | 0.05 | 8.25e-01 |

**Supplementary Table 12.** Imputed sum of parental genotypes based on the number of alternative alleles in siblings when IBD inference is missing or inconsistent, as described in [27] (see Methods 13). *q* denotes the alternative allele frequency in the respective cohort.

| Number of alternative alleles in sibling 1 | Number of alternative alleles in sibling 2 | Imputed sum of parental genotypes |
| --- | --- | --- |
| 0 | 0 | $2q/(2 - q)$ |
| 0 | 1 | $2/(2 - q)$ |
| 0 | 2 | 2 |
| 1 | 1 | $2 + (2q - 1)/(1 + q(1 - q))$ |
| 1 | 2 | $2(1 + 2q)/(1 + q)$ |
| 2 | 2 | $2(1 + 3q)/(1 + q)$ |

**Supplementary Figure 17.**
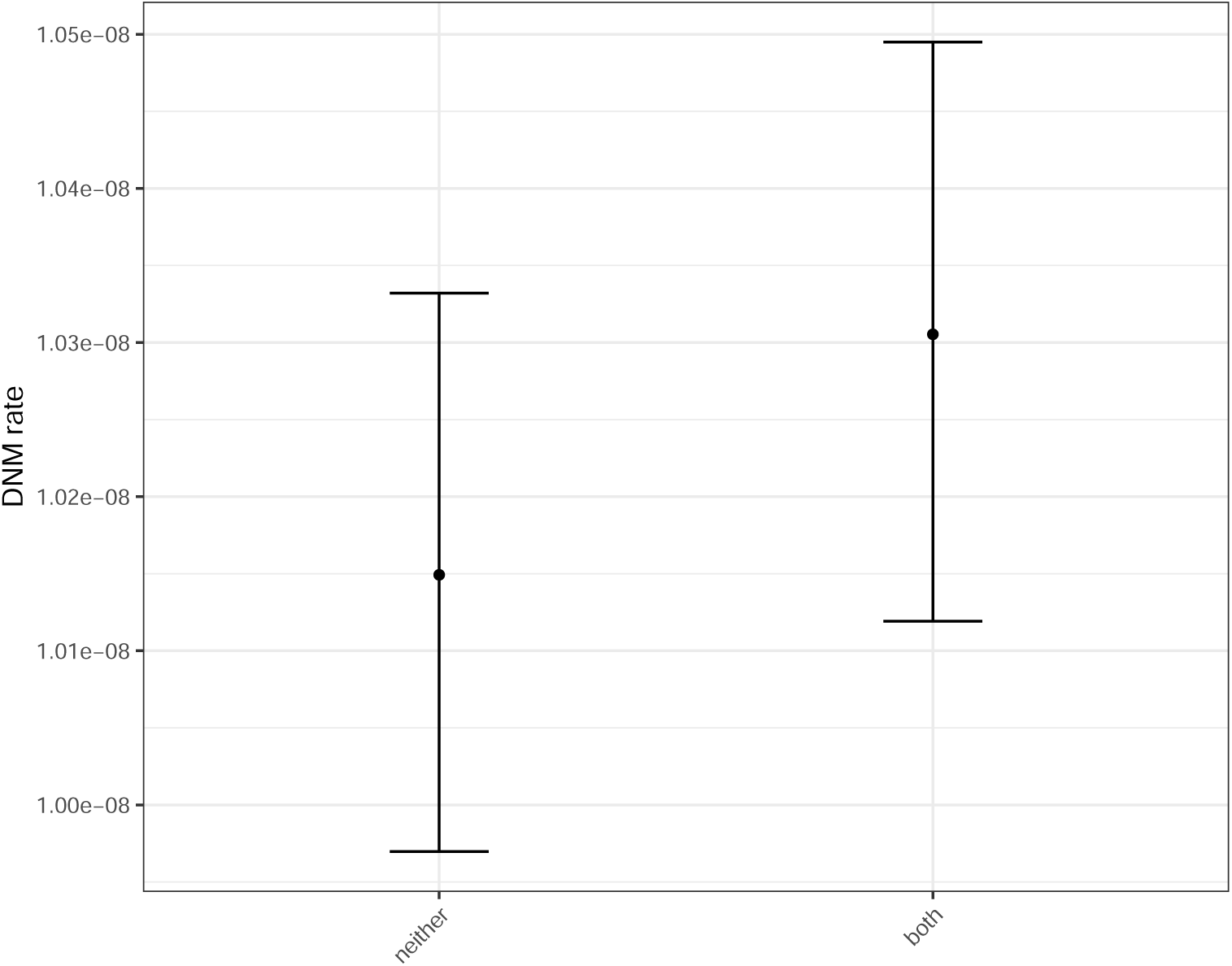
Estimates of autosomal DNM rates by parental smoking status at the sample mean paternal age at conception (27.7 years) and maternal age (25.4), with biobank set to UKB; shown are mean rates, with whiskers denoting 95% confidence intervals. Sample sizes were 829 for “neither” and 512 for “both”.

**Supplementary Figure 18.**
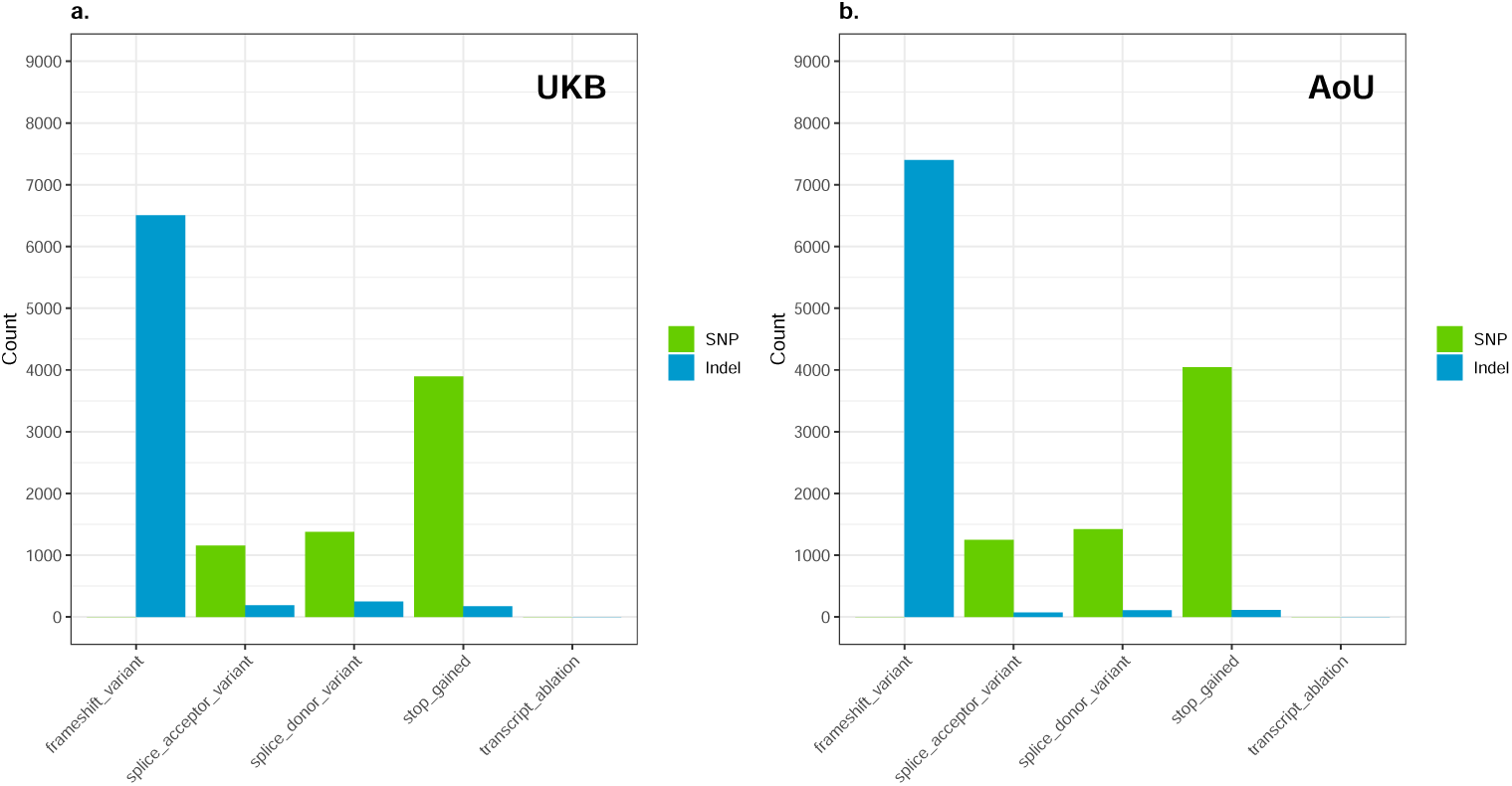
Functional annotations of the effects of LoF variants (see Methods 13) in UKB and AoU, for SNPs and indels.

**Supplementary Table 13.** Distribution of observed and imputed gene-level sums of parental genotypes for LoF and missense variants across 180 genes, based on 155 families with both parental and sibling data in AoU and UKB, where each case represents a gene-family pair.

| Cohort | Annotation | Observed sum | Imputed sum | Number of observed cases | Number of imputed cases | Proportion (%) |
| --- | --- | --- | --- | --- | --- | --- |
| AoU | LoF | 0 | < 1 | 27,563 | 27,560 | 99.99 |
| AoU | LoF | 0 | ≥ 1 | 27,563 | 3 | 0.01 |
| AoU | LoF | ≥ 1 | ≥ 1 | 337 | 265 | 78.6 |
| AoU | LoF | ≥ 1 | < 1 | 337 | 72 | 21.4 |
| AoU | Missense | 0 | < 1 | 27,791 | 27,789 | 99.99 |
| AoU | Missense | 0 | ≥ 1 | 27,791 | 2 | 0.01 |
| AoU | Missense | ≥ 1 | ≥ 1 | 109 | 94 | 86.2 |
| AoU | Missense | ≥ 1 | < 1 | 109 | 15 | 13.8 |
| UKB | LoF | 0 | < 1 | 6,281 | 6,280 | 100 |
| UKB | LoF | 0 | ≥ 1 | 6,281 | 1 | 0.02 |
| UKB | LoF | ≥ 1 | ≥ 1 | 19 | 12 | 63.2 |
| UKB | LoF | ≥ 1 | < 1 | 19 | 7 | 36.8 |
| UKB | Missense | 0 | < 1 | 6,279 | 6,279 | 100 |
| UKB | Missense | 0 | ≥ 1 | 6,279 | 0 | 0 |
| UKB | Missense | ≥ 1 | ≥ 1 | 21 | 15 | 71.4 |
| UKB | Missense | ≥ 1 | < 1 | 21 | 6 | 28.6 |
| AoU & UKB | LoF & Missense | 0 | < 1 | 67,914 | 67,908 | 99.99 |
| AoU & UKB | LoF & Missense | 0 | ≥ 1 | 67,914 | 6 | 0.01 |
| AoU & UKB | LoF & Missense | ≥ 1 | ≥ 1 | 486 | 386 | 79.4 |
| AoU & UKB | LoF & Missense | ≥ 1 | < 1 | 486 | 100 | 20.6 |

**Supplementary Table 14.** COSMIC SBS signature decompositions for carriers versus non-carriers of disruptive variants in *REV1* under an additive model (see Methods 13). Signature proportions were estimated using the EM framework of [39] as described in Methods 13.

| Signature | Fraction<br>in carriers | Fraction in<br>non-carriers |
| --- | --- | --- |
| SBS5 | 0.1617 | 0.5606 |
| SBS1 | 0.1505 | 0.1453 |
| SBS26 | 0.0778 | 0.0343 |
| SBS3 | 0.0731 | 0.0314 |
| SBS16 | 0.0661 | 0.0273 |
| SBS39 | 0.0574 | 0.0264 |
| SBS6 | 0.0510 | 0.0213 |
| SBS110 | 0.0507 | 0.0210 |
| SBS19 | 0.0366 | 0.0179 |
| SBS37 | 0.0356 | 0.0175 |
| SBS30 | 0.0282 | 0.0142 |
| SBS89 | 0.0271 | 0.0114 |
| SBS2 | 0.0258 | 0.0099 |
| SBS8 | 0.0236 | 0.0090 |
| SBS50 | 0.0190 | 0.0082 |
| SBS98 | 0.0162 | 0.0079 |
| SBS32 | 0.0151 | 0.0054 |
| SBS104 | 0.0138 | 0.0052 |
| SBS31 | 0.0137 | 0.0047 |
| SBS40b | 0.0106 | 0.0047 |
| SBS13 | 0.0074 | 0.0031 |
| SBS55 | 0.0067 | 0.0030 |
| SBS24 | 0.0058 | 0.0024 |
| SBS54 | 0.0052 | 0.0017 |
| SBS10b | 0.0051 | 0.0016 |
| SBS57 | 0.0042 | 0.0012 |
| SBS97 | 0.0031 | 0.0011 |
| SBS106 | 0.0022 | 0.0008 |
| SBS91 | 0.0020 | 0.0005 |
| SBS17a | 0.0015 | 0.0004 |
| SBS47 | 0.0010 | 0.0002 |
| SBS7a | 0.0009 | 0.0001 |
| SBS60 | 0.0008 | 0.0001 |
| SBS33 | 0.0006 | 0.0001 |
| SBS40a | 0.0001 | 0.0001 |
| SBS92 | 0.0001 | 0.0000 |

**Supplementary Table 15.** COSMIC SBS signature decompositions for carriers versus non-carriers of disruptive variants in *LIG1* under an additive model (see Methods 13). Signature proportions were estimated using the EM framework of [39] as described in Methods 13.

| Signature | Fraction<br>in carriers | Fraction in<br>non-carriers |
| --- | --- | --- |
| SBS5 | 0.4998 | 0.5602 |
| SBS1 | 0.1463 | 0.1453 |
| SBS16 | 0.0548 | 0.0343 |
| SBS6 | 0.0351 | 0.0314 |
| SBS86 | 0.0285 | 0.0274 |
| SBS32 | 0.0281 | 0.0266 |
| SBS30 | 0.0262 | 0.0215 |
| SBS24 | 0.0203 | 0.0210 |
| SBS12 | 0.0178 | 0.0179 |
| SBS87 | 0.0152 | 0.0175 |
| SBS10b | 0.0140 | 0.0142 |
| SBS92 | 0.0134 | 0.0115 |
| SBS21 | 0.0131 | 0.0099 |
| SBS102 | 0.0128 | 0.0090 |
| SBS55 | 0.0126 | 0.0081 |
| SBS19 | 0.0089 | 0.0078 |
| SBS97 | 0.0084 | 0.0055 |
| SBS46 | 0.0078 | 0.0052 |
| SBS106 | 0.0075 | 0.0047 |
| SBS54 | 0.0067 | 0.0046 |
| SBS17a | 0.0065 | 0.0032 |
| SBS2 | 0.0036 | 0.0030 |
| SBS23 | 0.0027 | 0.0024 |
| SBS98 | 0.0025 | 0.0017 |
| SBS99 | 0.0017 | 0.0016 |
| SBS13 | 0.0015 | 0.0012 |
| SBS110 | 0.0010 | 0.0011 |
| SBS48 | 0.0009 | 0.0008 |
| SBS7d | 0.0007 | 0.0005 |
| SBS26 | 0.0007 | 0.0004 |
| SBS29 | 0.0007 | 0.0003 |
| SBS39 | 0.0004 | 0.0001 |
| SBS3 | 0.0000 | 0.0001 |
| SBS101 | 0.0000 | 0.0001 |
| SBS84 | 0.0000 | 0.0001 |

